# Emission of floral volatiles is facilitated by cell-wall non-specific lipid transfer proteins

**DOI:** 10.1101/2022.11.30.518598

**Authors:** Pan Liao, Itay Maoz, Meng-Ling Shih, Ji Hee Lee, Xing-Qi Huang, John A. Morgan, Natalia Dudareva

## Abstract

For volatile organic compounds (VOCs) to be released from the plant cell into the atmosphere, they have to cross the plasma membrane, the cell wall, and the cuticle. However, how these hydrophobic compounds cross the hydrophilic cell wall is largely unknown. Using biochemical and reverse-genetic approaches combined with mathematical modeling, we show that cell-wall localized non-specific lipid transfer proteins (nsLTPs) facilitate VOC emission. Out of three highly expressed nsLTPs in petunia petals, which emit high levels of phenylpropanoid/benzenoid compounds, only PhnsLTP3 contributes to the VOC export across the cell wall to the cuticle. A decrease in *PhnsLTP3* expression reduces volatile emission and leads to VOC redistribution with less VOCs reaching the cuticle without affecting their total pools. This intracellular build-up of VOCs lowers their biosynthesis by feedback downregulation of phenylalanine precursor supply to prevent self-intoxication. Overall, these results demonstrate that nsLTPs are intrinsic members of the VOC emission network, which facilitate VOC diffusion across the cell wall.

## Introduction

Plants direct a substantial amount of photosynthetically fixed carbon (up to 10%) to the biosynthesis of volatile organic compounds (VOCs)^1^, which are emitted globally at a rate of ∼10^9^ metric tons per year^2^. Released from all organs, these amazingly diverse lipophilic low-molecular-weight molecules (100-200 Da) are used by plants to communicate and interact with their environment^3^. VOCs play key roles in attracting pollinators and seed dispersers, in above- and below-ground herbivore defense, protection from pathogens, plant-plant and inter-organ signaling, plant allelopathy and abiotic stress responses^3–5^. Given the VOC multifunctionality, much progress has been made over the last couple of decades towards elucidating the biosynthesis of plant VOCs, yet transport within and out of cells is poorly understood.

For many years, the assumption was that VOCs passively diffuse from the site of their biosynthesis through the cytosol, plasma membrane, hydrophilic cell wall and cuticle into the atmosphere. However, this common concept was recently challenged by showing that attaining observed emission rates by diffusion alone would necessitate high intracellular levels of VOCs, which would preferentially accumulate in the hydrophobic cellular compartments detrimentally affecting membrane integrity and function^6^. Therefore, biological mechanisms were proposed to facilitate VOC emission and prevent their accumulation to toxic levels^6^. Indeed, it was soon thereafter shown that efficient transport of phenylpropanoid/benzenoid volatiles across the plasma membrane in petunia flowers relies on the ATP-binding cassette (ABC) transporter (PhABCG1), the action of which prevents intracellular VOC accumulation and cellular self-intoxication^7^. Moreover, it was recently shown that although VOCs passage through the cuticle, the final barrier for efflux, occurs solely by diffusion, the cuticle acts as a sink/concentrator for VOCs. As an inert sink proximal to the environment, the cuticle contributes to modulating VOC emission while protecting cells from toxic intracellular accumulation of these volatiles^8^. Despite the latest progress in deciphering the emission process, little is known about how VOCs, which are primarily lipophilic, are transported from the plasma membrane across the hydrophilic cell wall to the cuticle.

The ability of non-specific lipid transfer proteins (nsLTPs) to transport a variety of hydrophobic molecules^9^ puts them in the spotlight as potential key players facilitating VOC transport across the cell wall. Indeed, nsLTPs were shown to function in the export of wax lipids and sesqui- and diterpenes from the plasma membrane, across the cell wall, to the cuticle^10–14^. Being widely distributed in higher plants, these proteins are frequently encoded by large gene families ^9,15,16^. In general, nsLTPs are small proteins with molecular mass ranging between 6.5-10.5 kDa, which contain a conserved motif of eight cysteines^15^. This unique motif forms four disulfide bridges that together with four α-helices create an inner hydrophobic cavity capable of transferring different lipophilic molecules^17^. To date, the biological role of the nsLTPs is not fully understood. However, increasing evidence suggests that these proteins are associated with numerous physiological processes including plant growth and reproduction, signaling, defense, and adaptation to stresses^9,17,18^.

Based on analogy to intracellular trafficking of other hydrophobic compounds, it was proposed that nsLTPs participate in the shuttling of VOCs across the aqueous cell wall space to the cuticle^6^, however, until now, this has not been experimentally validated. This type of transport facilitates diffusion, in which the presence of a carrier protein enhances the net flux without requiring cellular energy. In this study, we identified and characterized three nsLTPs in petunia flowers and analyzed their involvement in VOC emission. Although all three nsLTPs were localized to the cell wall, our results show that only one of them, PhnsLTP3, contributes to the export of VOCs across the cell wall to the cuticle *in planta*.

## Results

### Identification and characterization of *nsLTPs* in *Petunia hybrid*

A search of the petunia de novo assembled transcriptome^19^ for sequences with homology to two nsLTPs shown to be involved in trafficking of hydrophobic compounds across the cell wall, *Nicotiana tabacum* nsLTP1 (AB625593.1) and *Artemisia annua* nsLTP3 (KY712428.1)^13,14,20,21^ identified nine putative PhnsLTPs (**Fig. 1a**). As *PhnsLTPs* involved in VOC emission likely follow the developmental expression profiles of scent biosynthetic genes, which have the highest transcript levels on day 2 postanthesis, the expression of identified candidates was analyzed in our petunia petal RNA-Seq datasets^19^. Out of nine putative *PhnsLTP* genes, only four *PhnsLTPs* (*Ph19536, Ph18755, Ph18086*, and *Ph45320*) exhibited relatively high expression levels on day 2 postanthesis (**Fig. 1a**).

**Fig. 1.**
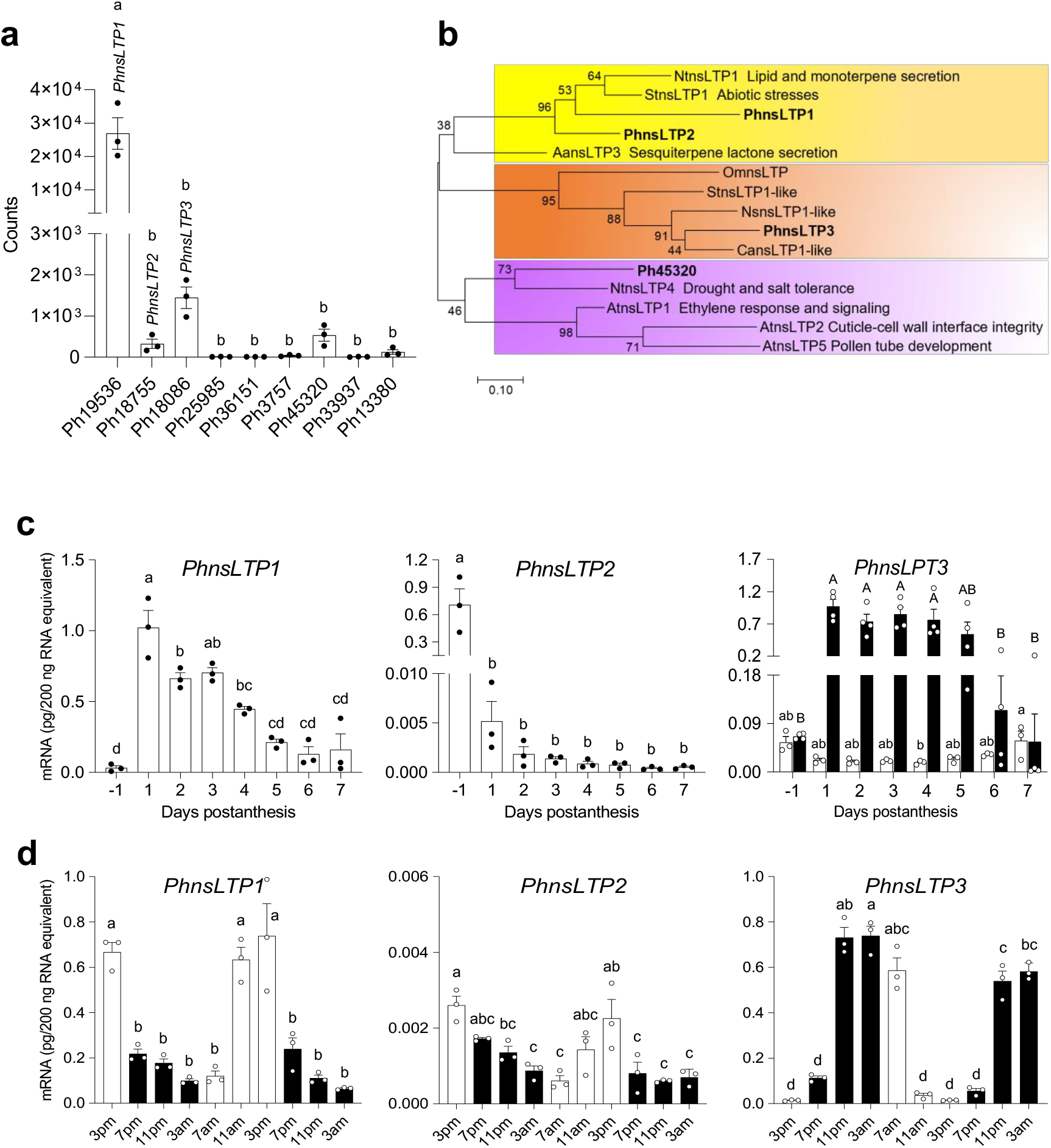
Expression analysis of petunia *nsLTP* candidates and phylogenetic analysis of PhnsLTP1, PhnsLTP2, PhnsLTP3 and Ph45320 in type I plant nsLTPs subfamily. **a**, Expression of nine *PhnsLTP* candidates in petunia flowers at 20:00 h on day 2 postanthesis. Gene expression was measured based on analysis of RNA seq datasets^19^ and represents an average of counts from three biological replicates ± S.E. **b**, The phylogenetic tree was built by the maximum-likelihood method in the MEGA7 package from the alignment of mature proteins using MUSCLE algorithm. At, *Arabidopsis thaliana*; Nt, *Nicotiana tabacum*; Aa, *Artemisia annua*; St, *Solanum tuberosum*; Ns, *Nicotiana sylvestris*; Ca, *Capsicum annuum*; and Om, *Oryza meyeriana var. granulate*. The scale bar indicates 0.1 amino acid substitutions per site. The percentage of replicate trees in which the associated taxa clustered together in the bootstrap test (1000 replicates) are shown next to the branches. **c**, Developmental expression profiles of *PhnsLTP1, PhnsLTP2* and *PhnsLTP3* in petunia corollas from bud stage through day 7 postanthesis determined at 15:00 h and at 23:00 h for *PhnsLTP3*. White columns represent the expression of *PhnsLTPs* at 15:00 h during flower development, black columns indicate the *PhnsLTP3* expression at 23:00 h during flower development. **d**, Changes in *PhnsLTPs* mRNA levels during a normal light/dark cycle in petunia corollas from day 1 to day 3 postanthesis. White and black columns correspond to light and dark periods, respectively. In (**c**) and (**d**) absolute mRNA levels were determined by qRT-PCR with gene-specific primers and are shown as pg/200 ng total RNA. Data are means ± S.E. (n = 3 biological replicates in **c** and **d**, and n = 4 biological replicates for *PhnsLTP3* at 23:00 h in **c**). Different letters indicate statistically significant differences (*P* < 0.05) determined by one-way ANOVA with the Tukey’s multiple comparisons test. In (c), capital letters were used to indicate significant differences among *PhnsLTP3* mRNA levels at 23:00 h during flower development.

Phylogenetic analysis of identified petunia PhnsLTPs along with nsLTPs from *A. thaliana* and *Oryza sativa* revealed that all nine petunia candidates belong to type I nsLTPs (Supplementary Figure 1). Out of four highly expressed candidates, *Ph19536, Ph18755, Ph18086*, and *Ph45320* (**Fig. 1a**), the latter was excluded from analysis as the annotated corresponding gene encodes a protein of 239 amino acids, which significantly exceeds the size of typical nsLTPs^15^ and clusters with plant nsLTPs known to affect cuticle properties^22,23^ (**Fig. 1b**). The other three candidates were used for further analysis and designated as *PhnsLTP1, PhnsLTP2* and *PhnsLTP3*, respectively. Quantitative RT-PCR (qRT-PCR) analysis with gene-specific primers revealed that *PhnsLTP1* displayed the highest level of expression in petunia petals on day 1 postanthesis and resembled the developmental expression profile typical for genes involved in the biosynthesis of volatiles^24,25^ (**Fig. 1c**). In contrast, *PhnsLTP2* expression was the highest in flower buds, which drastically decreased on day 1 postanthesis with a further reduction during the lifespan of the flower, while *PhnsLTP3* mRNA levels in corolla decreased after flower opening on day 1 postanthesis and remained relatively stable till day 7 when it recovered back to the level in the buds (**Fig. 1c**). Out of three candidates, rhythmic expression of only *PhnsLTP3* was the highest at night (**Fig. 1d**), which corresponds to the peak of emission of phenylalanine-derived petunia volatiles over a daily light/dark cycle^24,25^. Since the expression of *PhnsLTP* candidates was analyzed during flower development at 15:00 h, a time point with the highest expression for many scent biosynthetic genes^24,25^, when *PhnsLTP3* exhibited its lowest mRNA levels, the expression of the latter was also evaluated at 23:00 h. The *PhnsLTP3* developmental expression profile resembled that of scent biosynthetic genes^24,25^ being highest on day 1 postanthesis and decreasing afterward (**Fig. 1c**).

*PhnsLTP1* and *PhnsLTP2* encode proteins of 116 amino acids with calculated molecular masses for mature proteins of 9.35 kDa and 9.09 kDa, respectively (Supplementary Table 1). They share 58/69% identity/similarity with each other (Supplementary Table 2) and cluster together with other functional nsLTPs (**Fig. 1b**) including NtLTP1, which is involved in lipid secretion in tobacco and monoterpenes transport upon its overexpression in mint^13,26^, and AaLTP3 that is responsible for sesquiterpene lactone secretion^21^. *PhnsLTP3* encodes a protein of 115 amino acids with calculated molecular mass for mature protein of 9.65 kDa that shares 40/60% and 41/60% identity/similarity with PhnsLTP1 and PhnsLTP2, respectively (Supplementary Tables 1 and 2). In contrast to PhnsLTP1 and PhnsLTP2, PhnsLTP3 resides in a sister clade within the type I group containing several nsLTP1-like proteins with unknown functions (**Fig. 1b**).

### PhnsLTP1, PhnsLTP2 and PhnsLTP3 reside in the cell wall

In plants, nsLTPs have been detected in multiple subcellular locations including cytosol, plastids, plasma membranes, lipid-containing vesicles, and extracellularly in cell wall^18,27^. A signal peptide prediction program, SignalP, predicted that all three selected PhnsLTP candidates have a secretion signal (Supplementary Figure 2), suggesting their possible localization in the extracellular space. To experimentally determine their subcellular localization and potential involvement in the trafficking of VOCs from plasma membrane to the cuticle, the full-length coding region of each gene was fused to the red fluorescent protein (RFP) coding sequence and transiently expressed in *Nicotiana benthamiana* leaves. The Arabidopsis aquaporin1 (PIP1) fused with the green fluorescent protein (GFP) was used as a plasma membrane marker^28^. The RFP fluorescence signals from all PhnsLTP:RFP constructs were observed at the cell periphery, but they were almost indistinguishable from the signal from the co-transformed PIP1:GFP plasmid (Supplementary Figure 3). Therefore, prior to analysis, plasmolysis was used to separate the plasma membrane from the cell wall. Under these conditions, nsLTPs, as small and hydrophilic proteins, are expected to easily diffuse out of the cell wall. Indeed, after treatment of the detached transformed leaves with 20% sucrose for 60 min, signals from the PhnsLTP1:RFP, PhnsLTP2:RFP, and PhnsLTP3:RFP, but not empty RFP control, accumulated in free apoplastic place between the plasma membrane and cell wall demonstrating that all three PhnsLTPs are localized extracellularly, mostly in the cell wall (**Fig. 2**).

**Fig. 2.**
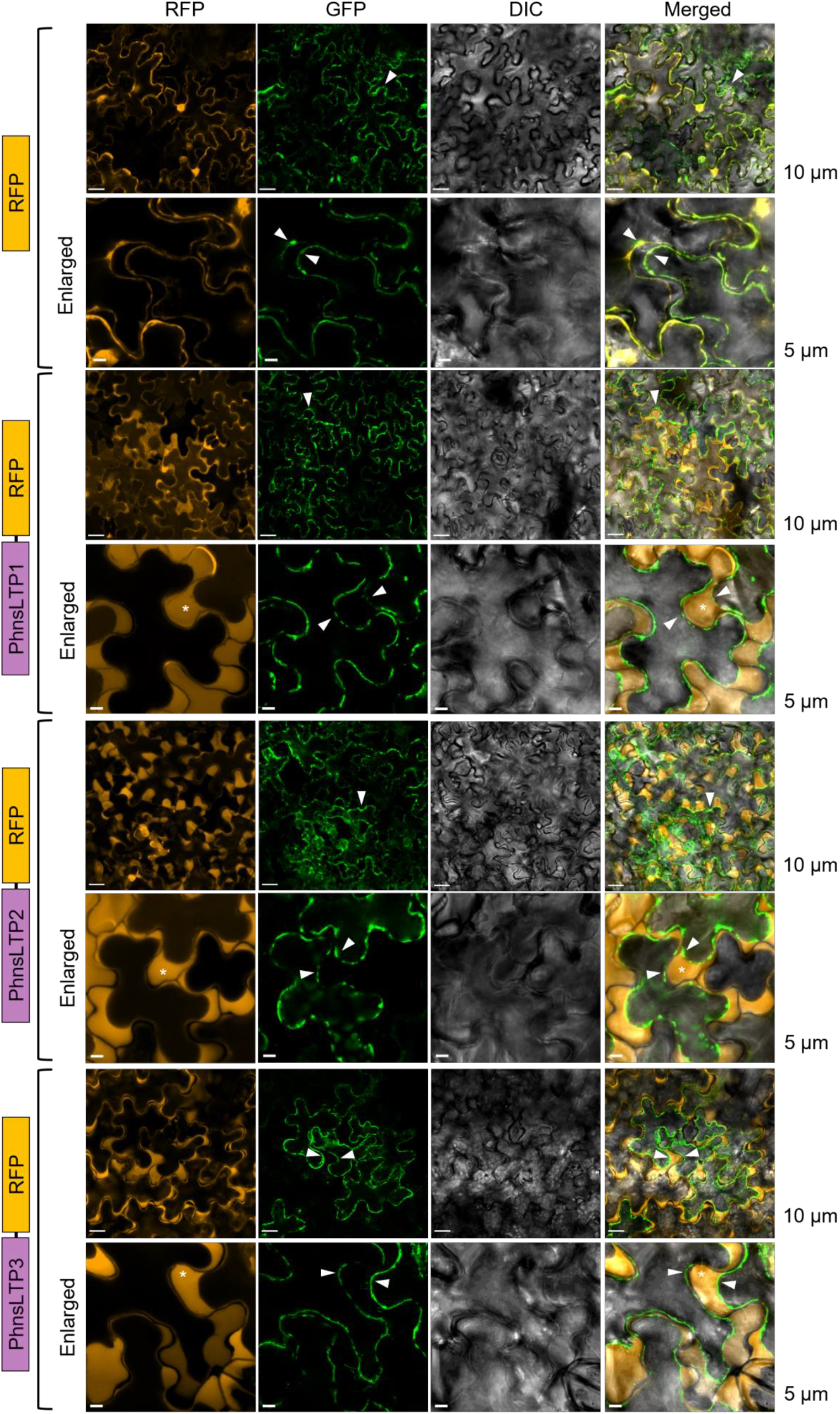
Subcellular localization of PhnsLTP1, PhnsLTP2 and PhnsLTP3. PhnsLTP1, PhnsLTP2, and PhnsLTP3 fusion constructs were expressed in *N. benthamiana* leaves and their corresponding transient expression was detected by confocal laser scanning microscopy. “RFP” panels (orange) represent signals of PhnsLTP1, PhnsLTP2, and PhnsLTP3 fused fluorescence proteins; the “GFP” panels (green) represent signals of the plasma membrane marker protein, AtPIP1:GFP; the DIC panels (dark grey) show images of differential-interference-contrast (DIC) microscopy; and the “Merged” panels show merged RFP, GFP and DIC signals. Prior to analysis, leaves were subjected to plasmolysis. Schematic diagrams of the PhnsLTP-RFP fusion proteins for each experiment are illustrated on the left. Triangles indicate the plasma membrane, and an asterisk shows the apoplastic space formed after plasmolysis. Images were captured using 20x (for low magnifications) and 63x oil immersion (for higher magnifications) objective lens. Scale bars were 5 μm or 10 μm and are shown on the right. The experiments were repeated three times with similar results.

### PhnsLTP3 is involved in emission of VOCs from petunia flowers

To assess the effect of *PhnsLTP1, PhnsLTP2* and *PhnsLTP3* on VOC emission, the expression of each gene was independently downregulated in petunia flowers using RNA interference (RNAi). In addition, transgenic plants with simultaneous downregulation of two genes *PhnsLTP1* and *PhnsLTP3*, the former exhibiting high expression on day 2 postanthesis (**Fig. 1a, c**), were generated. The RNAi constructs specifically targeted the desired genes and did not affect the expression of the other *PhnsLTPs*, which was verified by qRT-PCR with gene-specific primers (Supplementary Figures 4-6). For each construct, three independent lines with the greatest downregulation of a target gene were selected for further analysis (**Fig. 3**). Downregulation of *PhnsLTP1* expression by 52 to 77% (**Fig. 3a**) or *PhnsLTP2* mRNA levels by 47 to 61% (**Fig. 3d**) did not affect the emission of VOCs or their amount in petals (hereafter total pools) (**Fig. 3b, c, e, f**, and Supplementary Figures 7 and 8). In contrast, suppression of *PhnsLTP3* gene expression by 30 to 50% (**Fig. 3g**) led to the corresponding reduction in VOC emission by 33 to 49% (**Fig. 3h** and Supplementary Figure 9) without changing the total pool concentration (**Fig. 3j** and Supplementary Figure 10). No additive effect was observed when the transcript level of *PhnsLTP1*, a gene with the highest expression on day 2 postanthesis (**Fig. 1a, c**), was downregulated by 57 to 99% in addition to *PhnsLTP3* mRNA suppression by 58 to 79% (**Fig. 3i**). These *PhnsLTP1*+*PhnsLTP3-*RNAi transgenic plants displayed VOC emission and total pool profiles similar to that in *PhnsLTP3*-RNAi lines (**Fig. 3h, j;** Supplementary Figures 9 and 10), suggesting that PhnsLTP3, but not PhnsLTP1, contributes to VOC emission.

**Fig. 3.**
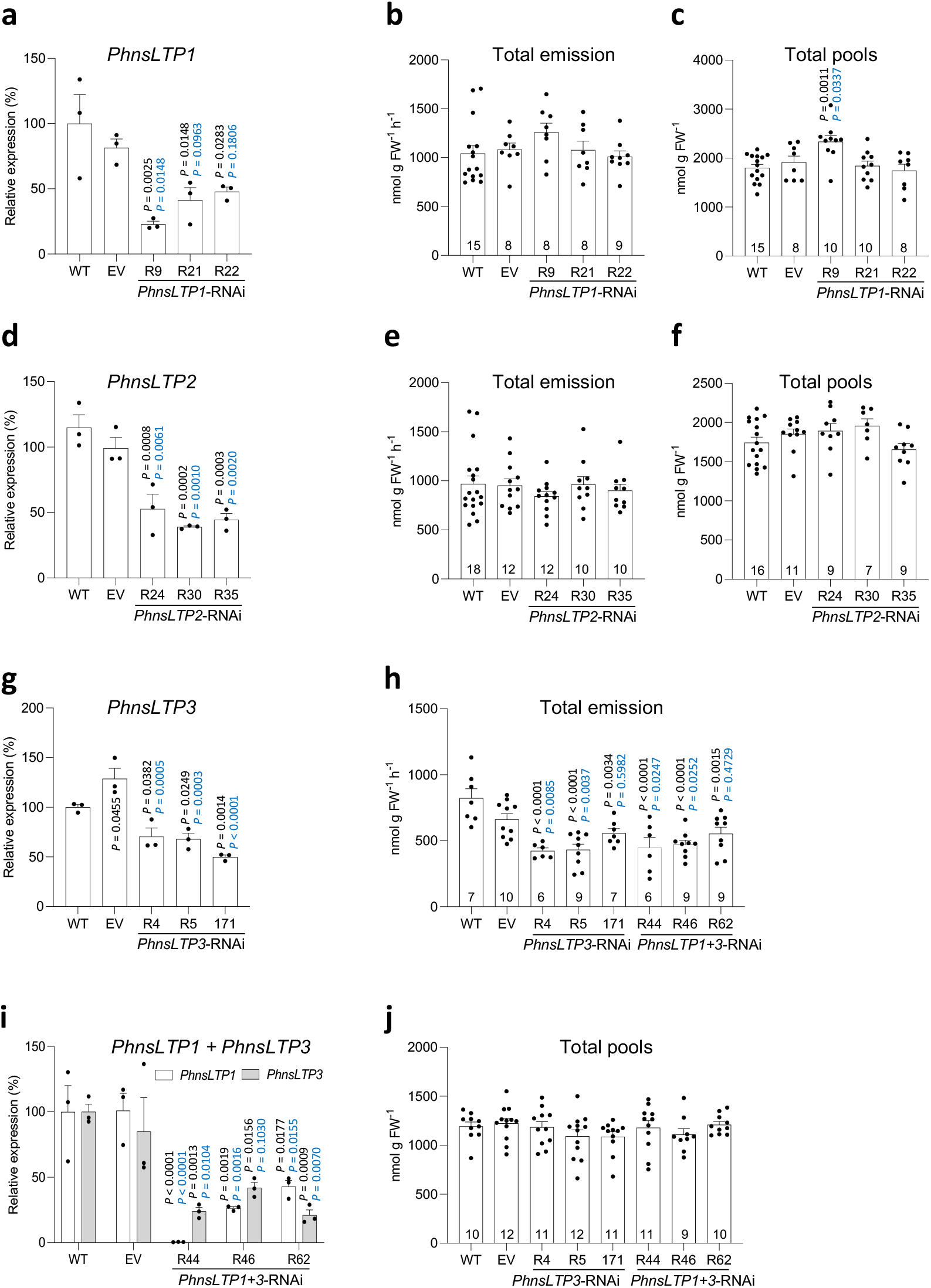
Effect of *PhnsLTP1, PhnsLTP2, PhnsLTP3* and *PhnsLTP1* + *PhnsLTP3* downregulation on emission of VOCs and their total pools in petunia flowers. Transcript levels of *PhnsLTP1* (**a**), *PhnsLTP2* (**d**), *PhnsLTP3* (**g**) and *PhnsLTP1* + *PhnsLTP3* (**i**) determined by qRT-PCR with gene-specific primers in 2-day-old wild-type petunia flowers (WT), empty vector control (EV) and the corresponding 3 independent transgenic lines with the greatest reduction in gene expression, harvested at 15:00 h. Data are shown as a percentage of a corresponding gene expression in WT set as 100% and are presented as means ± S.E. (n = 3 biological replicates). Total VOC emission (**b, e**, and **h**) and total pools (**c, f, j**) in 2-day-old flowers from WT, EV, and RNAi lines: *PhnsLTP1* (**b, c**), *PhnsLTP2* (**e, f**), *PhnsLTP3* and *PhnsLTP1* + *PhnsLTP3* (**h** and **j**). All emissions and total pools data are means ± S.E. Numbers at the bottoms of the columns in **b, c, e, f, h**, and **j** show the numbers of biological replicates. All *P* values were determined by one-way ANOVA with Dunnett’s multiple comparisons test relative to the WT (black) and EV (blue) controls except for **i**, for which *P* values were obtained by two-way ANOVA with Tukey’s multiple comparisons test. FW, fresh weight.

Total VOC pools, which remained unchanged in all transgenic plants except for *PhnsLTP1*-RNAi-R9 line (**Fig. 3**), do not provide information about VOC distribution within the cell plus cell wall and cuticle (hereafter cellular and cuticular pools, respectively). If PhnsLTP3 participates in the shuttling of VOCs across the cell wall to the cuticle, downregulation of its expression might lead to VOC redistribution between cellular and cuticular pools without affecting total VOC pools. Indeed, the cuticular abundance of all VOCs was decreased by 21 to 30% in *PhnsLTP3*-RNAi plants but not in *PhnsLTP1*-RNAi and *PhnsLTP2*-RNAi lines (**Fig. 4**), further supporting the involvement of PhnsLTP3 in VOC trafficking. The exception included *PhnsLTP2*-RNAi-R30 line, which displayed a statistically significant difference relative only to the wild-type control.

**Fig. 4.**
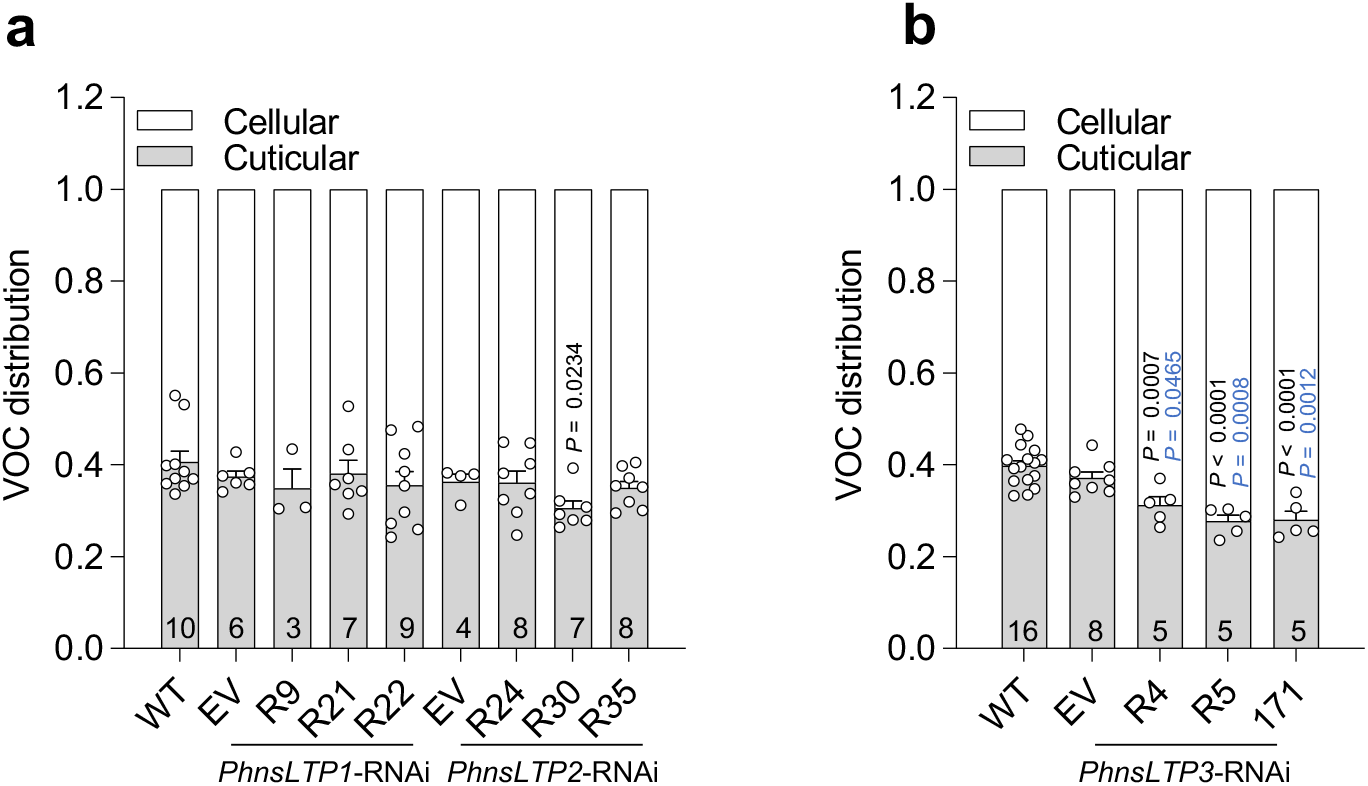
VOC distribution in 2-day-old petunia flowers. Cellular and cuticular VOC distribution in petunia flowers of WT, EV control, and three independent *PhnsLTP1*- and *PhnsLTP2*-RNAi lines (**a**) and *PhnsLTP3*-RNAi lines analyzed at 22:00 h (**b**). *P* values were determined by one-way ANOVA with Dunnett’s multiple comparisons test relative to the WT (black) and EV (blue) controls. All data are means ± S.E. Numbers at the bottoms of the columns show the numbers of biological replicates. Absolute VOCs’ amounts in cellular and cuticular fractions are presented in Supplementary Table 5.

nsLTPs are known to function in transporting cutin monomers across the cell wall to the cuticle^11–13,20^. It is possible that the observed VOC phenotypes (**Fig. 3b, c, e, f, h, j;** Supplementary Figures 9 and 10) are due to perturbations in cuticular wax. Thus, the total amount and chemical composition of wax were analyzed in wild-type and transgenic lines. No statistical differences in total wax amounts were found in *PhnsLTP1*-, *PhnsLTP2*-, *PhnsLTP3*-, and *PhnsLTP1*+*PhnsLTP3-* RNAi lines relative to the wild-type and empty-vector controls except for *PhnsLTP3*-RNAi-R5 line, which exhibited statistically significant difference relative to the empty-vector control only (Supplementary Figure 11). Small, but statistically significant differences, were found in wax constituents in individual lines, which were inconsistent among three transgenic lines generated for each *PhnsLTP1*-, *PhnsLTP2*-, *PhnsLTP3*-, and *PhnsLTP1*+*PhnsLTP3-*RNAi genotypes relative to the wild-type and empty-vector controls (Supplementary Figure 12). In addition, the petals of transgenic plants did not show staining by toluidine blue (Supplementary Figure 13), which detects increased cuticle permeability, retained water like that in the wild-type and empty vector controls (Supplementary Figure 14), and displayed consistent volatile emission and total pool phenotypes across transgenic genotypes (**Fig. 3**), suggesting that these small changes in cuticle composition did not affect the key cuticle properties.

### PhnsLTP3 displays petunia VOCs’ binding activity

To directly test whether PhnsLTP3 can bind petunia VOCs, in vitro displacement assays were performed using the recombinant PhnsLTP3 protein and fluorescent 6-(*p*-toluidino)-2-naphthalenesulfonate (TNS) in the presence of petunia volatiles. The coding region of *PhnsLTP3* lacking the N-terminal signal sequence was heterologously expressed in *Pichia pastoris GS115* as a 6× histidine-tagged protein, which was then purified by affinity chromatography on Ni-NTA agarose (Supplementary Figure 15a and b). TNS fluoresces strongly when bound to the hydrophobic regions of the protein, while it has weak fluorescence in an aqueous solution. Indeed, intense TNS fluorescence was detected in the presence of PhnsLTP3 protein relative to buffer and denatured and reduced PhnsLTP3 protein controls (Supplementary Figure 15c). Potential ligands of the protein will displace the bound TNS from the hydrophobic protein pocket into the aqueous environment leading to a decrease in the fluorescent signal, which was used to calculate the dissociation constant of the protein with ligand. Our results revealed that PhnsLTP3 binds VOCs, including benzaldehyde, benzyl alcohol, 2-phenylethanol, methylbenzoate, benzylbenzoate and vanillin with the dissociation constants (K_d_) ranging from 6.5 ± 2.7 μM for benzylbenzoate to 396 ± 131 μM for 2-phenylethanol (**Fig. 5**). However, we were unable to calculate PhnsLTP3 dissociation constants with phenylacetaldehyde, eugenol, and isoeugenol due to high background TNS fluorescence in the presence of these compounds. To understand why the downregulation of *PhnsLTP1* and *PhnsLTP2* does not affect VOC emission (**Fig. 3a-f** and Supplementary Figures 7 and 8), we also characterized the VOCs’ binding activity of the corresponding proteins (Supplementary Figure 15a-c). While PhnsLTP1 did not display TNS binding activity (Supplementary Figure 15c), the dissociation constants for PhnsLTP2.VOCs’ interactions were in the micromolar range as for PhnsLTP3 (Supplementary Figure 16), suggesting that PhnsLTP2 has the potential to contribute to VOC emission in petunia flowers. To test whether PhnsLTP2 and PhnsLTP3 can bind other VOCs, we assayed the terpenoids: limonene, linalool, and nerolidol, as well as methyl jasmonate in TNS displacement assays. Both PhnsLTP2 and PhnsLTP3 were able to bind tested volatile compounds with K_d_s in the micromolar range (Supplementary Figures 16 and 17).

**Fig. 5.**
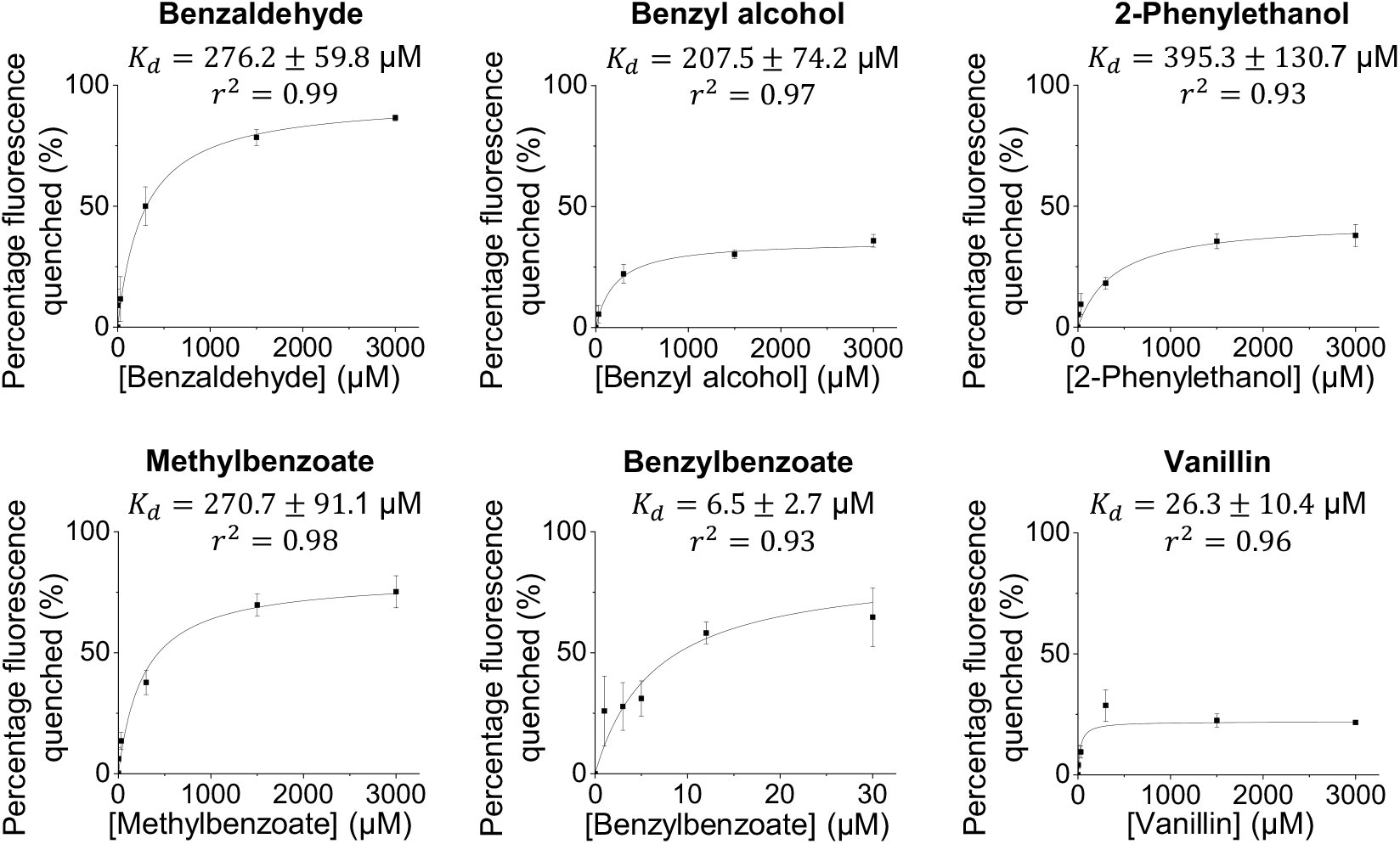
Binding kinetics of recombinant PhnsLTP3 with VOCs emitted by petunia flowers. TNS displacement assays were performed at fixed concentrations of PhnsLTP3 (1 μM) and TNS (3 μM) and the fluorescence was measured before and after the tested compounds were added at various concentrations. The results are expressed as percentage of fluorescence quenched with respect to the concentration of added compound. The binding curves were obtained by non-linear fitting to the Michaelis-Menten equation (solid line in the graph) and the dissociation constants *K*_*d*_ [μM] were obtained. Phenylacetaldehyde, eugenol, and isoeugenol had high fluorescence background with TNS, which did not allow obtaining the corresponding binding curves. All data are means ± S.E. (n = 3 biological replicates).

### Mathematical modeling of facilitated diffusion of VOCs by PhnsLTP3

To quantitatively assess the influence of PhnsLTP3 on VOC emission flux and estimate the flux changes in response to decreased PhnsLTP3 levels in the cell wall, a steady-state mathematical model based on facilitated diffusion^29^ (Supplementary Figure 18) was used. A steady-state model was selected here as it allows to reduce the simulation complexity and provides a valid approach to assess the effects of key parameters on VOC efflux. In contrast to active transport, facilitated diffusion is a spontaneous passive process, in which the molecules move down a concentration gradient across a biological compartment by specific carrier proteins. Thus, we hypothesized that the water-soluble nsLTPs localized in the cell wall could facilitate the transport of the non-polar molecules through this aqueous cellular compartment. According to the model, the effect on VOC flux depends on several parameters including (*i*) the cell wall nsLTP concentration, (*ii)* the nsLTP.VOC binding constant, and (*iii*) the difference in VOC concentrations between the apical facing side of the plasma membrane and the inner side of the cuticle (Supplementary Figure 18).

First, the effect of PhnsLTP3 concentrations in the cell wall on the emission flux was evaluated over a wide range of physiologically relevant concentrations from 1 μM to 1 mM (**Fig. 6a**) as we were unable to experimentally determine the PhnsLTP3 concentration within the cell wall of epidermal cells. As expected, the flux enhancement was linearly proportional to the PhnsLTP3 concentration. The highest extent of enhanced flux was observed for vanillin and the lowest for 2-phenyethanol, which correlated with the distance of the corresponding dissociation constant from the optimal value determined by the model (see below and **Fig. 6b**). Next, we compared the model predictions with the experimentally measured decrease in VOC emission in the *PhnsLTP3*-RNAi lines. For this, PhnsLTP3 concentration in wild type was set as 1 mM, a reasonable, high physiological value (**Fig. 6a**). The PhnsLTP3 concentration in *PhnsLTP3*-RNAi lines was estimated based on the 40% decrease in the corresponding gene transcript level assuming that protein abundance is proportional to gene expression. Using these parameters, model simulation revealed that the total VOC flux would be 18% lower in the *PhnsLTP3*-RNAi lines than in the wild type, thus underestimating the reduction in VOC emission flux relative to the measured fluxes in the RNAi lines.

**Fig. 6.**
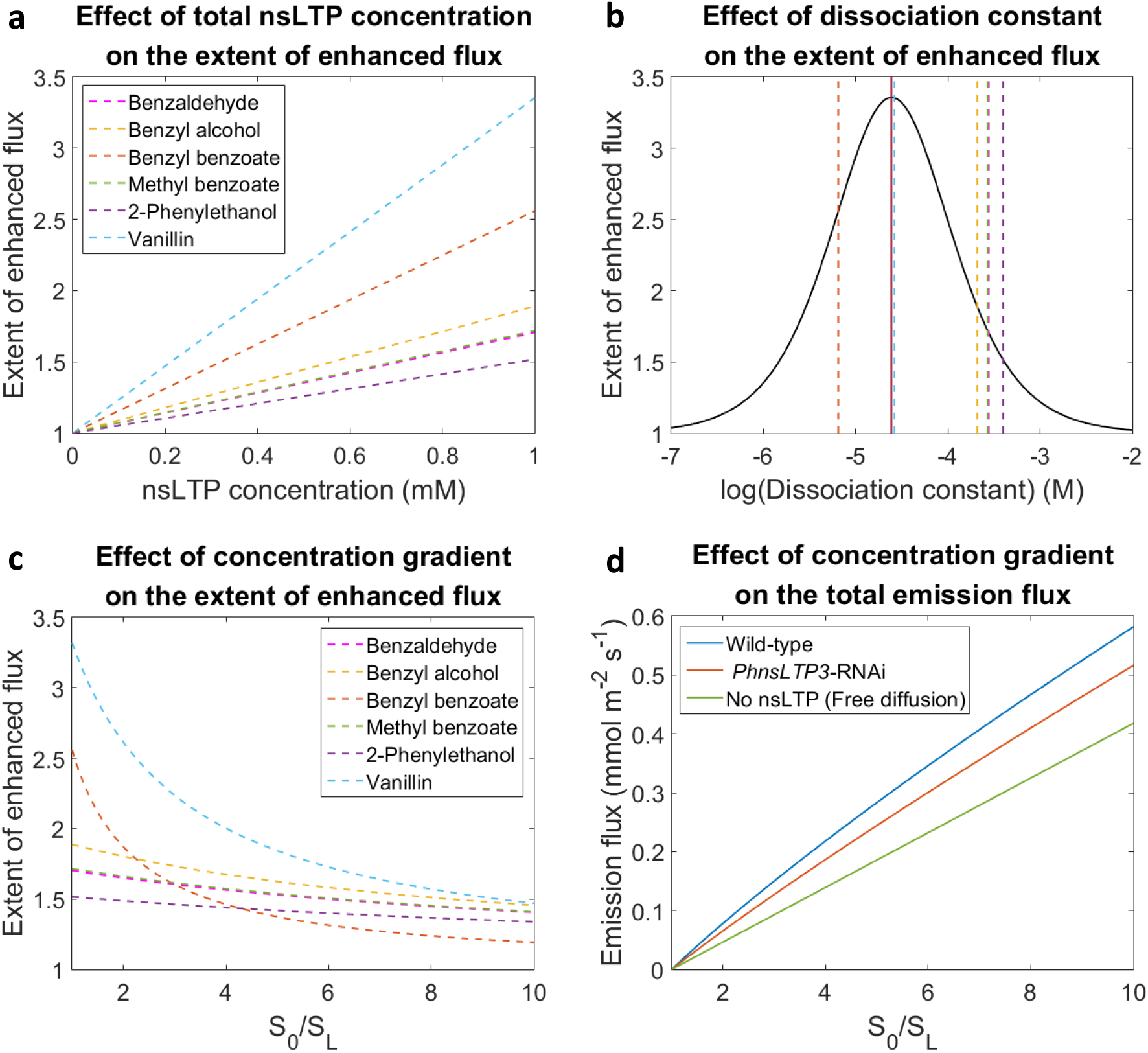
The effect of total nsLTP concentration (a) and dissociation constant (b) on the extent of enhanced flux, and concentration gradient across the cell wall (S_0_/S_L_) on the extent of enhanced flux (c) and the total VOC emission flux (d). **a**, The extent of enhanced flux was plotted against different nsLTP levels at a fixed S_0_/S_L_ ratio and measured respective dissociation constants. **b**, The extent of enhanced flux (black line) was plotted against different dissociation constants at a fixed S_0_/S_L_ ratio and nsLTP level (1 mM). The vertical red line indicates the predicted optimal dissociation constant, and the six vertical dotted lines show the experimentally obtained in the TNS binding assay dissociation constants of PhnsLTP3 with benzylbenzoate (orange), vanillin (light blue), benzyl alcohol (yellow), methylbenzoate (green), benzaldehyde (pink), and 2-phenylethanol (purple) from left to right, respectively. **c**, The extent of enhanced flux was plotted against different S_0_/S_L_ ratios at a fixed nsLTP level (1 mM) and measured respective dissociation constants. **d**, The total emission flux was plotted against different S_0_/S_L_ ratios assuming nsLTP levels in wild-type 1 mM (blue), in *PhnsLTP3*-RNAi flowers 0.6 mM (orange), and no nsLTP (green). Abbreviations: S_0_ and S_L_ are VOC concentrations at the apical facing side of the plasma membrane and the inner side of the cuticle, respectively (see Supplementary Figure 18). The extent of enhanced flux (E.o.F) in **a – c** and emission flux (*J*) in **d** were calculated based on Eq 9 and Eq 8, respectively (see Supplementary Material). The values of the parameters used are provided in Supplementary Table 6.

In addition, the mathematical model simulations were performed to calculate the enhancement of flux as a function of the nsLTP.VOC binding constant. The model parameters were set as follows: the nsLTP concentration constant at 1mM and the VOC gradient across the cell wall at an arbitrary concentration difference of 0.25 nM. Assuming a single VOC binding site model, the simulation shows an optimal dissociation constant of 24.4 μM, where the maximum enhancement of flux by nsLTP occurs (**Fig. 6b**). Similar to previous models of facilitated transport, the simulated curve has a bell shape^29^. The model predicted that if the dissociation constant is high (> 1 mM), the ability of the carrier protein to bind the ligands is limited hence leading to a low enhancement factor. In contrast, if the dissociation constant is too small (< 1 μM), meaning that the affinity between the protein and the ligand is strong, the protein will hold the ligand tightly, which restricts the release of the VOC to the cuticle and limits the overall mass transfer. Therefore, the maximal enhancement of flux by nsLTPs is best achieved when the balance between the binding and the unloading of VOC is reached.

VOC emission from petunia flowers changes rhythmically during a day/night cycle^24,25^, however, to date it is not possible to directly ascertain the VOC concentration gradient across the cell wall. Thus, a range of concentration gradients was set up in the model simulation to examine the effect on emission flux. The simulations demonstrate that the extent of enhanced VOC flux by nsLTP is the highest when the VOC concentration gradient across the cell wall is shallow being higher than one at all concentrations (**Fig. 6c**) meaning that the VOC transport across the hydrophilic cell wall always benefits from the presence of the nsLTPs. Consistent with this prediction, regardless of the VOC concentration gradient across the cell wall, the simulated absolute VOC flux in both wild-type and *PhnsLTP3*-RNAi line is always higher when compared to free diffusion (**Fig. 6d**) due to nsLTP facilitation of the VOC trafficking across this hydrophilic barrier.

### PhnsLTP3 facilitates VOC emission

The reduction in VOC emission in *PhnsLTP3*-RNAi lines without changes in their total pools indicates that VOC biosynthetic flux is reduced. Such reduction in VOC emission could be the result of diminished VOC transport across the cell wall, decreased VOC biosynthesis or both. Our results show that *PhnsLTP3* downregulation leads to VOC redistribution, resulting in ∼25% fewer VOCs on average in the cuticle (**Fig. 4b**). Recently, we have shown that the VOC redistribution and reduction of their levels in the cuticle decrease their release from the cell and reduce carbon flux to Phe, which is the precursor for petunia VOCs^8^. Thus, the expression of selected genes involved in Phe biosynthesis was analyzed and found, as in petunia flowers with thinner cuticle^8^, to be downregulated by 17 to 38 % in *PhnsLTP3*-RNAi flowers relative to the wild-type control (Supplementary Figure 19). Thus, to distinguish between the contribution of VOC transport and Phe limitation in VOC biosynthesis, control and *PhnsLTP3*-RNAi-R5 flowers were fed with Phe, and total VOC emission and total pools were measured from 19:00 to 22:00 h (Supplementary Figure 20). While total pools in transgenics were similar to that in wild type, Phe feeding was unable to restore emission flux in *PhnsLTP3-*RNAi-R5 flowers, which was overall 30.1 ± 8.9% (*P* = 0.0086, two-way ANOVA) lower than in the wild type. However, the absolute increase over 2 h in emission rate upon feeding was similar in transgenic and wild-type flowers (1,569 ± 76 nmol g FW^-1^ h^-1^ and 1,444 ± 298 nmol g FW^-1^ h^-1^ in *PhnsLTP3*-RNAi-R5 and wild-type flowers, respectively) (Supplementary Figure 20a), indicating that VOC biosynthetic capacity downstream of Phe was unaffected in transgenic flowers. Analysis of activities of enzymes involved in the VOC biosynthesis including phenylalanine ammonia lyase (PAL), benzoyl-CoA:benzylalcohol/2-phenylethanol benzoyltransferase (BPBT) and benzoic acid carboxyl methyltransferase (BAMT) at 20:00 h on day 2 postanthesis further supported that they were unaffected in *PhnsLTP3*-RNAi lines (Supplementary Figure 21), suggesting that both Phe supply and transport by PhnsLTP3 contribute to VOC emission.

Our model suggested that nsLTPs might facilitate the emission the most when the VOC concentration gradient across the cell wall is shallow. Since VOC emission in petunia flowers occurs in a rhythmic manner, with the maximum during the night^24,25,30^, which can potentially result in different concentration gradients across the cell wall, the nsLTP contribution to emission flux was additionally analyzed during the onset phase of nocturnal VOC emission from 16:00 to 19:00 h. VOC emission factors (VEFs), which account for reduced VOC biosynthesis and are a ratio of emission flux to the corresponding total biosynthetic flux^8^, were compared in wild-type and transgenic flowers under physiological conditions (without Phe feeding) at two different time periods during a day/night cycle. It should be noted that the total VOC pool is the balance between the incoming and outgoing fluxes at a single time point. To get an insight into the flux changes, the individual total pool measurements were first converted into the rate of change in the total pool by estimating the slope from linear regression. VEFs were calculated in wild-type and *PhnsLTP3-* RNAi-R5 transgenic flowers from 17:00 to 19:00 h and 20:00 to 22:00 h (**Fig. 7** and Supplementary Table 3). The VEFs continuously increase in both wild-type and transgenic flowers at both time periods, but the effect of PhnsLTP3 on VEFs became apparent only after emission flux reached ∼ 50% of biosynthetic flux at 19:00 h (VEF = 0.5) (**Fig. 7e** and **f**). The exact reasons for it are currently unknown but could be the result of combined changes in the nsLTP levels and VOC concentration gradient across the cell wall. VEFs in transgenics were lower than in the wild type suggesting the existence of VOC transport limitations in *PhnsLTP3-*RNAi flowers.

**Fig. 7.**
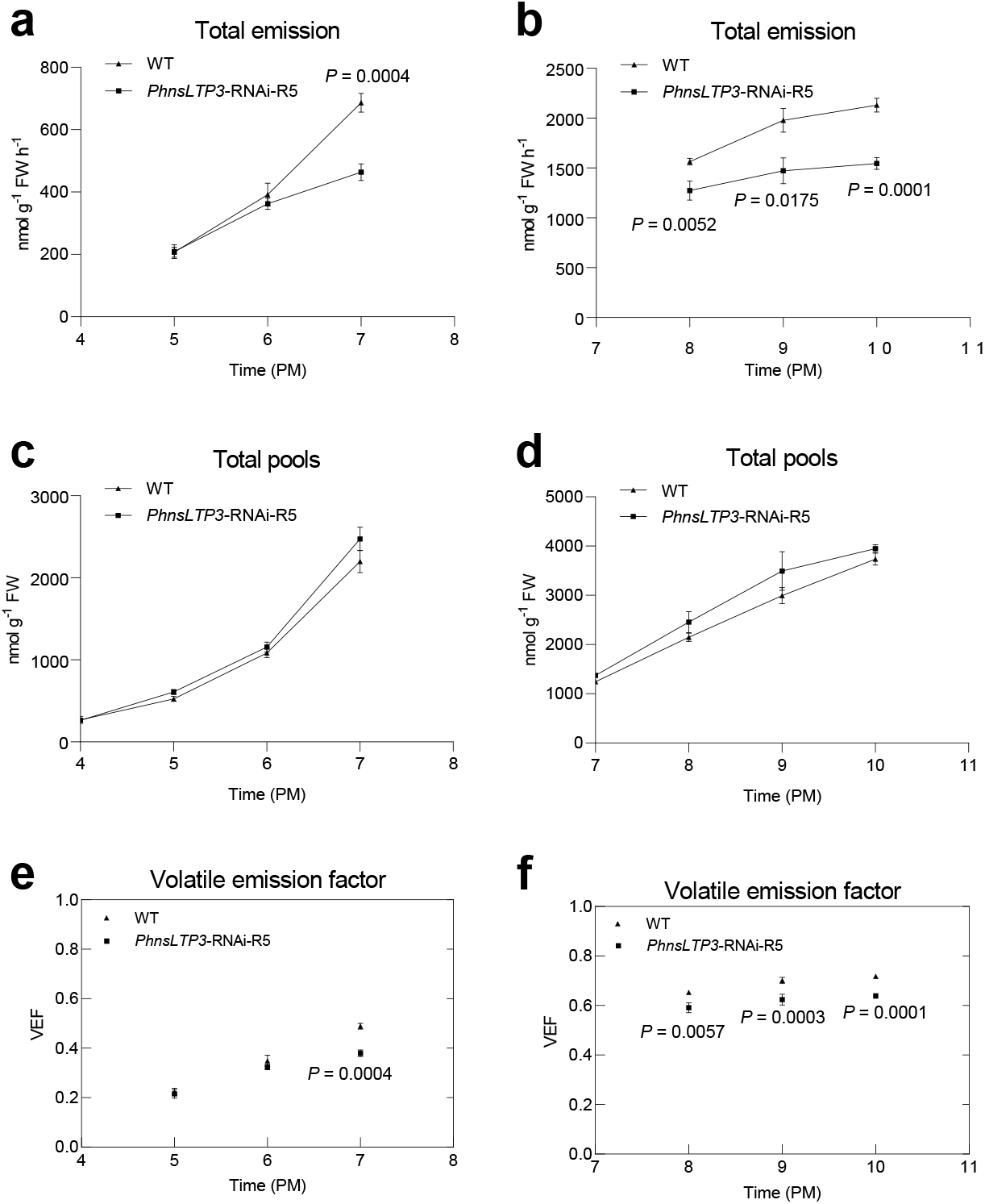
Total VOC emission, total pools, and the VOC VEFs in control and *PhnsLTP3*-RNAi petunia flowers. Total VOC emission in wild-type (WT) and *PhnsLTP3*-RNAi-R5 line from 16:00 to 19:00 h (**a**) and 19:00 to 22:00 h (**b**). Total pools in WT and *PhnsLTP3*-RNAi-R5 line from 16:00 to 19:00 h (**c**) and 19:00 to 22:00 h (**d**). VEFs in WT and *PhnsLTP3*-RNAi-R5 line from 16:00 to 19:00 h (**e**) and 19:00 to 22:00 h (**f**). In (**a**) and (**c**), data are means ± S.E. (n = 9 and 5 biological replicates for WT and *PhnsLTP3*-RNAi-R5 line, respectively at 17:00 h, 18:00 h, and 19:00 h, except that n = 3 biological replicates for WT and *PhnsLTP3*-RNAi-R5 line respectively at 16:00 h); in (**b**) and (**d**), n = 8, 8, 7 and 8 biological replicates for WT, and n = 4, 4, 5 and 5 biological replicates for *PhnsLTP3*-RNAi-R5 line, respectively, at 19:00 h, 20:00 h, 21:00 h, and 22:00 h. *P* values were determined by a two-tailed Student‘s *t*-test for the emission and total pools between WT and *PhnsLTP3*-RNAi-R5 line at each time point. In (**e**), data are means ± S.E. (n = 9 and 5 biological replicates for WT and *PhnsLTP3*-RNAi-R5 line, respectively at 17:00 h, 18:00 h, and 19:00 h). In (**f**), data are means ± S.E. (n = 8, 7 and 8 biological replicates for WT, and n = 4, 5 and 5 biological replicates for *PhnsLTP3*-RNAi-R5 line, respectively, at 20:00 h, 21:00 h, and 22:00 h. In (**e**) and (**f**), *P* values were determined by two-way ANOVA with the Sidak’s multiple comparisons test relative to the corresponding WT at each time point.

## Discussion

It has been shown that nsLTPs are involved in delivering wax components to the cuticle^11,12,20^, as well as in the secretion of monoterpenes^26^, sesquiterpenes^14,21^ and diterpenes^13^ from the secretory head cells to the subcuticular storage cavities of glandular trichomes. More than half a decade ago, it was proposed that nsLTPs could also facilitate VOC transport across the hydrophilic cell wall *en route* to the atmosphere^6^. Here, we provide genetic evidence that nsLTPs indeed participate in VOC trafficking across the cell wall to the cuticle thus directly contributing to VOC emission. Out of three highly expressed *PhnsLTP* genes (**Fig. 1a, c-d**) in petunia petals, which emit high levels of phenylpropanoid/benzenoid compounds, only downregulation of *PhnsLTP3* affects emitted VOCs (**Fig. 3** and Supplementary Figures 7-9) without consistently altering the total wax amount or its composition (Supplementary Figures 11 and 12). Reduced emission (**Fig. 3h**) in *PhnsLTP3*-RNAi lines as well as the inability of transgenic flowers to recover upon feeding with Phe to the total emission levels of wild-type flowers (Supplementary Figure 20) indicate the existence of transport limitation(s) upstream of the cuticle and the involvement of PhnsLTP3 in VOC trafficking. PhnsLTP3 resides in a sister clade within the type I nsLTPs containing several nsLTP1-like proteins with unknown functions (**Fig. 1b**) and shares only 40 to 41% amino acid identity with PhnsLTP1 and PhnsLTP2 (Supplementary Table 2). Both in vitro binding experiments (**Fig. 5**) and in vivo analysis of PhnsLTP3 function (**Fig. 3g-j and 4b**) provide evidence that PhnsLTP3 binds and transports phenylpropanoid/benzenoid compounds. The ability of nsLTPs to bind fatty acids and terpenoids has been demonstrated before^11–14,20^, but, to our knowledge, this is the first report of nsLTP binding to phenylpropanoid and benzenoid compounds. However, there was no correlation between PhnsLTP3 dissociation constants with individual VOCs and a reduction in the emission of the corresponding compound upon *PhnsLTP3* downregulation (Supplementary Figure 22). This could be explained by the fact that flowers produce a mixture of different VOCs at different levels, which compete for the LTP binding site, as well as *S*_0_ and *S*_L_ VOC concentrations being dependent on the efficiency of plasma membrane ABC transporter for individual compounds and their partitioning to the cuticle, respectively.

PhnsLTP2 and PhnsLTP3 have overlapping ligand specificity (**Fig. 5** and Supplementary Figures 16 and 17) and both are localized in the cell wall (**Fig. 2**), but only PhnsLTP3 is involved in VOC emission (**Fig. 3h**). The reasons why transgenic petunia flowers only with PhnsLTP3 suppression have reduced emission fluxes are currently unknown, but it could be partially explained by the differences in expression of *PhnsLTP3* and *PhnsLTP2*, as the mRNA level of the latter drastically decreases after flower opening on day 1 postanthesis (**Fig. 1c**). Additionally, only *PhnsLTP3* exhibits a rhythmic profile with the highest transcript level out of three *PhnsLTPs* at 23:00 h on day 1 postanthesis, which is > 500-fold higher than that of *PhnsLTP2* (**Fig. 1d**), and correlates with emission of petunia volatiles^24,25^. Moreover, PhnsLTP2 and PhnsLTP3 belong to two separate clades within the type I plant nsLTP (**Fig. 1b)**. Clustering of PhnsLTP2 with plant nsLTPs known to be involved in terpenoid secretion combined with its highest expression in flower buds (**Fig. 1c**) when petunia flowers produce and emit terpenoid compounds^31^, might suggest that PhnsLTP2 and PhnsLTP3 functions are temporally separated and they likely contribute to the transport of different classes of compounds, terpenoids and phenylpropanoids, respectively. Indeed, when *PhnsLTP2* and *PhnsLTP3* expressions were analyzed in petunia stigma, only *PhnsLTP2* mRNA levels resembled the expression of petunia terpene synthases^31^ (Supplementary Figure 23).

In general, VOCs are synthesized de novo in the tissues from which they are emitted. The biosynthesis of VOCs in flowers occurs almost exclusively in epidermal cells, which are in closest proximity to the atmosphere^32-35^ for release. Before being released, VOCs move from the site of their biosynthesis through the cytosol, plasma membrane, hydrophilic cell wall, and cuticle and each of these cellular barriers provides resistance to the overall emission rate^6^. In petunia, VOC transport across the plasma membrane is mediated by the PhABCG1 transporter^7^ and across the cell wall by PhnsLTP3 (**Fig. 3**), while VOCs move by passive diffusion through the cuticle, which imposes significant resistance to mass transfer and serves as VOC sink/concentrator^8^. An increase in a barrier’s resistance either via *PhABCG1*-RNAi or *PhnsLTP3*-RNAi downregulation leads to a decrease in VOC emission^7^ (**Fig. 3h** and Supplementary Figures 9), which also occurs upon reduction in the cuticle thickness as a result of RNAi downregulation of PhABCG12 transporter or dewaxing^8^. However, the alterations of the cellular barrier properties have different effects on total VOC pools and cell intoxication. Total pools increased in *PhABCG1*-RNAi flowers^7^ and decreased in flowers with reduced cuticle thickness^8^ but remained unchanged upon *PhnsLTP3*-RNAi downregulation (**Fig. 3j** and Supplementary Figure 10). Despite unaffected total pools, up to 30% fewer VOCs accumulated in the cuticle of *PhnsLTP3*-RNAi transgenic petunia flowers, leading, as in the case of flowers with thinner cuticles, to their build-up within the cell (**Fig. 4b**). Such intracellular accumulation of VOCs led, similar to the situation in flowers with reduced cuticle thickness^8^, to a reduction in overall VOC formation by transcriptional feedback downregulation of genes involved in the biosynthesis of Phe precursor (Supplementary Figure 19). In contrast to flowers with altered cuticle^8^ and downregulated *PhABCG1* transporter^7^, the intracellular build-up of VOCs in cells of *PhnsLTP3*-RNAi flowers did not affect membrane integrity, as staining of *PhnsLTP3*-RNAi and *PhnsLTP1+PhnsLTP3*-RNAi petals with propidium iodide, which diffuses into cells and stains nucleic acids only if plasma membrane is damaged^36^, showed no nuclei staining (Supplementary Figure 24). Taken together, these results imply that the effect of altered cellular barrier resistance on VOC biosynthesis, emission, and total pools is barrier specific. Moreover, these data also suggest that the release of individual volatiles to the atmosphere is determined by the substrate specificity of the plasma membrane transporter, as PhnsLTP3 has broad substrate specificity and can carry both phenylpropanoid and terpenoid compounds (**Fig. 5** and Supplementary Figure 17) and there is no specificity for volatile diffusion through the cuticle. Indeed, as PhABCG1 transports exclusively phenylpropanoid/benzenoid compounds^7^, the overexpression of *Clarkia breweri* (*S*)-linalool synthase in petunia did not result, despite its production, in the emission of linalool, which was sequestered as a nonvolatile linalool glycoside to prevent cell intoxication^37^. In addition, the rate of release through the cell wall and cuticle depends on the affinity of PhnsLTP3 towards individual VOCs and their physicochemical properties^38^.

Facilitated VOC transport by nsLTPs is analogous to that of O_2_ transport by myoglobin, the carrier protein in muscles that enhances the transport of poorly water-soluble molecules^39^. Therefore, a mathematical model based on facilitated diffusion^29^ (Supplementary Figure 18) was utilized to describe the nsLTP-mediated transport across the cell wall. The model predicted that the total VOC flux would decrease by 18% in the *PhnsLTP3*-RNAi lines relative to wild type assuming that PhnsLTP3 concentration in wild type is 1 mM (**Fig. 6a**) and is 40% lower in transgenic flowers proportionally to *PhnsLTP3* gene expression (**Fig. 3g**). However, the VOC emission flux was reduced by 33 to 49% in the *PhnsLTP3*-RNAi lines relative to wild type (**Fig. 3h** and Supplementary Figure 9). This discrepancy could be due to the fact that the model considered the same VOC production and the concentration gradient across the cell wall in both backgrounds. While in transgenic plants VOC biosynthesis was reduced as a result of a decrease in the supply of Phe precursor for petunia phenylpropanoid/benzenoid compounds^8^ (Supplementary Figure 19), currently, there are no experimental approaches to measure the VOC concentration gradient across the cell wall. However, when the shortage in precursor supply was eliminated by Phe feeding (Supplementary Figure 20a), emission in transgenic flowers was overall 30.1 ± 8.9% lower than in control flowers, which is closer to the model prediction. Additionally, we do not exclude that the PhnsLTP3 concentration in the cell wall could be different from 1 mM, which was used in the modeling. Moreover, the dynamic changes in VOC gradients across the cell wall between the transgenic and wild-type flowers were not taken into account as they cannot be captured by a steady-state model.

The model predicted that larger enhancement of flux relies more on LTP concentration than on VOC gradient across the cell wall (**Fig. 6a, c** and **d**). Several crystal structures of plant nsLTPs show that more than one ligand could bind in their hydrophobic pocket^17^. Since the facilitation of the total flux would be higher when the carrier proteins can bind multiple ligands^40^, the capability of PhnsLTP3 to bind more than one VOC was considered by fitting the binding curve to the two-binding-site model, in which the binding sites were assumed to be independent. Among the experimentally tested phenylpropanoids/benzenoid compounds (**Fig. 5**), only PhnsLTP3 with methylbenzoate showed a statistically better fit to a two-binding-site model (*P* value < 0.05; Supplementary Figure 25). Assuming the total PhnsLTP3 concentration is 1 mM and the VOC concentration difference between the cell wall boundaries is 0.25 nM, the extent of the enhancement in methylbenzoate flux would increase from 1.67 in the single-binding-site model to 1.98 in the two-binding-site model. The lack of statistical improvement in fitting with the two-binding-site model for other tested compounds does not rule out that PhnsLTP3 can bind more than one VOC. Certainly, protein-ligand crystallization studies would help to determine the number of VOCs that PhnsLTP3 can bind.

Overall, analysis of the role of nsLTPs in VOC emission revealed that nsLTPs, specifically PhnsLTP3, contribute to the export of VOCs from the plasma membrane, across the cell wall, to the cuticle. The obtained information will not only allow us to identify new metabolic engineering targets for altering VOCs released from the plant but can also be potentially adapted to reduce the release of undesired odor from plants.

## Material and methods

### Chemicals and reagents

Benzyl alcohol, benzaldehyde, benzylbenzoate, methylbenzoate, 2-phenylethanol, phenylacetaldehyde, eugenol, isoeugenol, vanillin, 6-(*p*-toluidino)-2-naphthalenesulfonic acid sodium salt (TNS), toluidine blue, *n*-tetracosane, L-phenylalanine and naphthalene were purchased from Sigma-Aldrich (http://www.sigmaaldrich.com).

### Plant material and growth conditions

Wild-type and transgenic *P. hybrida* cv. Mitchell diploid (W115, Ball Seed Co.) plants were grown under standard greenhouse conditions with a light period from 6:00 to 21:00 h. All four *PhnsLTP1*-, *PhnsLTP2*-, *PhnsLTP3*-, *PhnsLTP1+ PhnsLTP3*-RNAi constructs were synthesized by Genscript. Before synthesis, the constructs (Supplementary Figures 4 - 6) were verified to target only the desired genes and not to lead to off-target interference using the Sol Genomics Network VIGS Tool (http://vigs.solgenomics.net/). The analysis was performed by comparing the *PhnsLTP1-, PhnsLTP2-* and *PhnsLTP3*-RNAi target sequences against the *P. axillaris* and *P. inflata* (parents of *P. hybrida*) genomes^41^ using default parameters, except that the n-mer setting was adjusted to 24 nucleotides, which is generally considered to be below the threshold to efficiently trigger downregulation^42,43^. This analysis revealed that no other genes would be targeted by the constructs’ double-stranded RNA triggers. Except for *PhnsLTP1*, each of the synthetic RNAi fragments was placed under the control of the CaMV-35S promoter in the binary destination vector pK2GW7 using Gateway LR Clonase II kit (Invitrogen). *PhnsLTP1*-RNAi construct was subcloned into a modified pRNA69 vector containing the *Clarkia breweri* linalool synthase (LIS) promoter^44^ and the entire cassette containing the *LIS* promoter, *PhnsLTP1* hairpin structure, and the OCS terminator was cut off and ligated into pART27 binary vector. Transgenic *P. hybrida* plants were obtained via *Agrobacterium tumefaciens* (strain GV3101 carrying the final constructs) leaf disc transformation using a standard transformation protocol^45^. Further analysis of RNAi downregulation of a target gene on expression of the other *PhnsLTP* homologs determined by qRT–PCR with gene-specific primers revealed that their expression remained unchanged (Supplementary Figures 4 - 6).

### Analysis of RNA-Seq datasets for *PhnsLTP* genes

In general, *nsLTP* genes are small in size and share relatively high sequence homology^15^, therefore, to accurately calculate *PhnsLTPs’* expression levels in previously generated RNA-Seq datasets^19^, we utilized the HISAT2-HTSeq workflow^46^. The amino acid sequence of *Nicotiana tabacum* non-specific lipid-transfer protein (BAK19150) and *Artemisia annua* nsLTP3 (AST22305) were used as queries against *P. hybrida* protein sequence dataset obtained from the de novo transcriptome assembly of our RNA-Seq data^19^. The corresponding nucleotide sequence of each hit was BLAST against the current version of *Petunia axillaris* genome (v1.6.2) to retrieve the correct genomic sequences manually. While most of the identified *PhnsLTPs* have already been annotated in the *P. axillaris* genome, this analysis enabled us to annotate one additional *nsLTP* gene. Then, the identified genomic region of each gene was used for proper mapping of the raw RNA-Seq reads using HISAT2 program^47^. The number of reads mapped to each gene was obtained with HTSeq package^48^. As a result, the expression levels were analyzed for nine *PhnsLTP* genes including the existing accessions and newly annotated *nsLTP* in the following genomic regions: Peaxi162Scf00595g00036 (*Ph19536, PhnsLTP1*), Peaxi162Scf00732g00334 (*Ph18755, PhnsLTP2*), Peaxi162Scf00251:996872-997595 (*Ph18086, PhnsLTP3*), Peaxi162Scf01238g00223 (*Ph13380*), Peaxi162Scf00732g00317 (*Ph3757*), Peaxi162Scf00732g00338 (*Ph45320*), Peaxi162Scf00129g00232 (*Ph25985*), Peaxi162Scf01213g00004 (*Ph33937*) and Peaxi162Scf00395g00012 (*Ph36151*).

### Phylogenetic analysis of PhnsLTPs

Amino acid sequences of nsLTPs were aligned using MUSCLE algorithm in the MEGA7 package^49^. The evolutionary history was inferred by the maximum likelihood method^50^, and the evolutionary distances were computed using the Poisson correction model^50^. The numbers at the nodes correspond to the percentage of replicate trees, which resolve the clade in the bootstrap test (500 replicates for Supplementary Figure 1 and 1000 replicates for **Fig. 1b**). Phylogenetic trees were constructed using MEGA7 package^44^ and included 123 and 15 amino acid sequences for Supplementary Figures 1 and **Fig. 1b**, respectively. All positions with less than 95% site coverage were eliminated. Protein sequences used to generate the figures are available in Supplementary Datasets S1 and S2.

### Quantitative RT-PCR (qRT-PCR) analysis

Sample collection, RNA isolation, and qRT-PCR were conducted as previously described^8^. Briefly, samples were obtained from the tissues indicated in the text and RNA was isolated using the Spectrum™ Plant Total RNA Kit (Sigma-Aldrich). Total RNA (1 μg) was treated with DNase I (Promega) before reverse transcription to cDNA using a 5×All-In-One RT MasterMix (Applied Biological Materials Inc.). *PhnsLTP1, PhnsLTP2*, and *PhnsLTP3* expression was analyzed using Fast SYBR Green Master Mix (Applied Biosystems) with gene-specific primers (Supplementary Table 4) in StepOnePlus PCR system (Applied Biosystems) with StepOne software (v2.2.2). For relative expression quantification, *Elongation factor 1-α* was used as a reference gene (Supplementary Table 4), and calculations were performed using the 2^ΔΔCt^ method^51^. The absolute quantification of transcript levels was carried out based on standard curves generated from qRT-PCR with gene-specific primers using respective purified cDNA fragments diluted to several concentrations between 0.1 fg μl^-1^ and 0.01 ng μl^-1^. Absolute amounts of individual transcripts were calculated and displayed as picograms (pg) of mRNA per 200 ng of total RNA. Each data point represents one biological sample consisting of three flowers. Primers used for the absolute quantification of *PhnsLTP1, PhnsLTP2*, and *PhnsLTP3* transcript levels and generating standard curves are listed in Supplementary Table 4.

### Subcellular localization of PhnsLTP1, PhnsLTP2 and PhnsLTP3

The full-length coding regions of *PhnsLTP1, PhnsLTP2*, and *PhnsLTP3* were amplified using corresponding gene-specific primers (Supplementary Table 4) and the resulting amplicons were inserted into pDONR vector (Invitrogen, Carlsbad, CA). After sequence verification, the generated constructs were transferred by recombination using LR Clonase II (Invitrogen) into pK7RWG2 vector, which expresses fusion proteins with a C-terminal red fluorescent protein (RFP), resulting in constructs expressing PhnsLTP1:RFP, PhnsLTP2:RFP and PhnsLTP3:RFP. *AtPIP1* (aquaporin 1) was used as a plasma membrane marker^28^. It was similarly subcloned into pK7FWG2 vector (Supplementary Table 4) to produce a fusion protein with C-terminal green fluorescent protein (GFP), AtPIP1:GFP. The resulting constructs and an empty RFP vector were transformed into *A. tumefaciens* strain GV3101, infiltrated into *Nicotiana benthamiana* leaves and imaged 48 h after. For plasmolysis, leaf discs were incubated in 20% sucrose for 60 min prior to imaging^52^. Images were acquired using a Zeiss LSM 880 Upright Confocal system and operated using Zeiss Efficient Navigation (ZEN)-Black Edition software (Zen 2.6, Carl Zeiss Inc.). The excitation wavelength and emission bandwidth were optimized by the default settings in the ZEN 2.6 software (Zeiss) and were for GFP as excitation 488 nm, emission 493-556 nm, and for RFP as excitation 561 nm, emission 580-651 nm.

### Heterologous expression of PhnsLTPs

The open reading frames encoding mature PhnsLTP1, PhnsLTP2 and PhnsLTP3 proteins (omitting N-terminal signal peptide) were synthesized (Twist, Bioscience, CA, USA) with codon optimization for expression in *Pichia pastoris*. The coding region of each PhnsLTP was subcloned into the EcoRI and AccI sites of pPICZα vector (Invitrogen) in frame with the N-terminal α factor secretion signal and the C-terminal 6×His-tag.

The resulting plasmids were verified by sequencing and transformed into *P. pastoris*, strain GS115 using the EasySelect™ Pichia Expression Kit (Invitrogen). Transformants were selected on plates containing 100, 200, 500, or 1000 μg/mL Zeocin. Colonies were verified for the presence of insert by PCR with the forward α factor primer 5’-TACTATTGCCAGCATTGCTGC-3’ and the reverse AOX1 primer 5’-GCAAATGGCATTCTGACATCC-3’, the latter is located downstream of 6× His-tag. A single positive colony for each PhnsLTP was inoculated in 10 mL Buffered Glycerol-complex Medium (BMGY, containing 1% yeast extract, 2% peptone, 100 mM potassium phosphate, pH 6.0, 1.34% yeast nitrogen base (YNB), 4 μg L^-1^ biotin, and 1% glycerol) and incubated at 30°C with shaking at 275 rpm until the OD_600_ reached 2-6. This 10 mL starter culture was then inoculated in 1 L BMGY and incubated under the same conditions until OD_600_ = 2-6. Cells were harvested by centrifugation at 2500× *g* for 5 min at room temperature and the pellet was resuspended in 100 mL Buffered Methanol-complex Medium (BMMY, which has a similar composition as BMGY with 0.5% methanol instead of 1% glycerol) to induce expression. The culture was incubated for 48 h at 30°C with shaking at 275 rpm followed by centrifugation at 10,000× *g* for 30 min at room temperature. Ammonium sulfate was then added to the supernatant for 80% saturation and the mixture was gently rotated for 30 min at 4°C in an orbital shaker at 25 rpm followed by centrifugation at 10,000×*g* for 30 min at 4°C. Pellet was resuspended in buffer containing 50 mM NaH_2_PO_4_, 300 mM NaCl, and 20 mM imidazole adjusted to pH 8.0. Purification was performed on nickel-nitrilotriacetic acid (Ni-NTA) agarose using the QIAexpressionist™ kit (Qiagen) according to the manufacturer’s protocol. The purity of proteins was analyzed with 4-15% polyacrylamide precast gels (BIO-RAD) followed by their staining with Coomassie brilliant blue R-250 after electrophoresis.

### TNS binding assays

The fluorescence-based binding assays were performed as previously described^12,53^ with slight modifications. Briefly, assays were performed in the 96-well plates with each well containing a fixed concentration of PhnsLTP (1 μM) and TNS (3 μM) or TNS only in 175 mM mannitol, 0.5 mM K_2_SO_4_, 0.5 mM CaCl_2_, and 5 mM MES (pH 7.0). All mixtures were excited at 320 nm, and the fluorescence emission at 437 nm was recorded using a plate reader (SpectraMax iD3, Molecular Devices) at equilibrium after 5 min of incubation at 25°C giving the background fluorescence 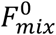and 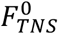. The tested ligands dissolved in 2 μL DMSO were then added to the mixtures and TNS only to different final concentrations, and the fluorescence emissions were recorded at equilibrium after 5 min of incubation (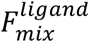and 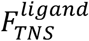). To account for the change in the TNS fluorescence intensity upon adding the solvent DMSO, the background fluorescence of TNS 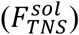 as well as each PhnsLTP with TNS 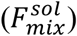 were measured. The ratios 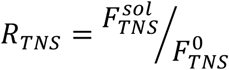and 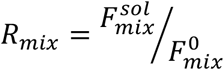were calculated to calibrate the fluorescence. Three measurements were taken for each concentration. The results were expressed as a percentage of fluorescence quenched 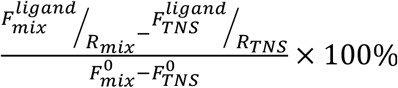. The corresponding dissociation constants *k*_*d*,_ were obtained by nonlinear regression fitting of the binding kinetics to the Michaelis-Menten equation.

### Collection and analysis of plant volatiles and phenylalanine feeding

Floral volatile emission was analyzed by dynamic headspace collection of volatiles at 25°C on columns containing 20 mg Porapak Q (80–100 mesh) (Sigma-Aldrich) and analyzed on an Agilent 6890N-5975B GC–MS system as described previously with minor modifications^7,8,44^. Briefly, emitted volatiles were collected from detached transgenic and wild-type flowers from 19:00 to 22:00 h on day 2 postanthesis. Adsorbed volatiles were eluted from collection columns with 280 μL dichloromethane containing 52 nmol of naphthalene as the internal standard. For analysis of total petal pools, 0.5 g of petal tissue from transgenic and control plants was collected at 22:00 h on day 2 postanthesis, ground to a fine powder in liquid nitrogen, and total petal pools were extracted in 5 mL of dichloromethane containing 52 nmol of internal standard (naphthalene) for 12 h at -20°C. Samples were then warmed up, vortexed and concentrated to 200 μL under a stream of nitrogen gas before injection into GC–MS (Agilent 6890N-5975B). All GC–MS data were collected and analyzed using Agilent MSD ChemStation F.01.03.2357 software. For phenylalanine feeding, flowers from wild type and *PhnsLTP3*-RNAi were collected and placed in 150 mM of phenylalanine solution. Emitted volatiles and their total petal pools were sampled every hour from 16:00 till 19:00 h and from 19:00 till 22:00 h and analyzed as described above.

### Toluidine blue staining and water loss measurements

Toluidine blue staining was carried out as previously described^8,54^. Petunia flowers (day 2 postanthesis) were collected at 15:00, and the adaxial surfaces of the petals were submerged in 0.05% (w/v) toluidine blue aqueous solution for 4 h. After rinsing in distilled water, petals were used for imaging. Water loss measurements were performed as described previously^8^.

### Cuticular wax extraction and analysis

Petals from wild-type and transgenic petunia flowers collected at 18:00 h on day 2 postanthesis were used for wax extractions. The adaxial surfaces of the petals were submerged in 10 mL hexane for 30 s. The solvent was transferred to a glass vial and dried completely under a gentle stream of nitrogen gas. The dried wax residues were derivatized using 200 μL of *N,O*-bis(trimethylsilyl)trifluoroacetamide (BSTFA) (derivatization grade, Sigma-Aldrich)/pyridine mixture (1:1, v/v) for 20 min at 90°C. After evaporation of excess derivatization solution under a stream of nitrogen gas, the derivatized waxes were solubilized in 100 μL hexane. The wax samples were analyzed by GC-MS. A capillary column (HP-5MS, 30 m × 0.25 mm, 0.25 μm, Agilent) was used to separate wax compounds with helium carrier gas at a flow rate of 1.0 mL min^-1^. A 2 μL sample was injected in splitless mode. The initial oven temperature was 40°C for 2 min and ramped up to 200°C at 15°C min^Δ1^, followed by a hold for 2 min at 200°C. Then, the temperature was increased from 200°C to 320°C at 3°C min^Δ1^ and held for 15 min. Compounds were identified based on retention times and fragmentation patterns of authentic standards. Quantification of compounds was done by integrating peak areas and using calibration curves of wax compounds including fatty acids, primary alcohols, and alkanes with varying chain lengths. *n*-Tetracosane (99%, Sigma-Aldrich) was used as an internal standard. The total wax amount was calculated per dry petal weight with eight petals per biological replicate.

### Analysis of volatiles in the cuticle

To analyze the quantity of volatiles in petunia epicuticular waxes, flowers were collected at 22:00 h on day 2 postanthesis, quickly dipped in hexane (<5 s) and lightly dried under the fume hood to remove excess hexane. Care was taken to prevent full submersion of petals so that only the adaxial surface of the flower touched the solvent. Hexane pools were separately profiled for volatiles and wax quantity by GC-MS as described above. Total cuticular volatile fractions were calculated according to the formula as described previously^8^:

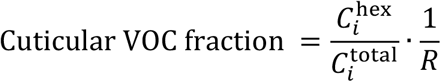

where 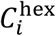is the volatile pool in the hexane fraction, 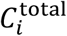is the total volatile pool, and *R* is the fraction of the total wax recovered during dipping in hexane. Absolute VOCs’ amounts in cellular and cuticular fractions were calculated by multiplying total pools shown in Fig. 3c, f and j with corresponding distribution values (Supplementary Table 5).

### Propidium iodide staining and confocal laser scanning microscopy

Petals of wild-type, *PhnsLTP3*-RNAi and *PhnsLTP1+ PhnsLTP3*-RNAi flowers collected at 15:00 h on day 2 postanthesis were stained in propidium iodide solution (10 μg mL^-1^, Invitrogen) for 1 h at room temperature with shaking. Confocal fluorescence microscopy was conducted as described previously^8^.

### PAL, BPBT and BAMT activity

PAL, BPBT and BAMT activities were measured in transgenic *PhnsLTP3*-RNAi and wild-type flowers collected at 20:00 h on day 2 postanthesis. Crude protein extracts were prepared from petal tissues in buffer containing 100 mM Tris pH 7.4, 150 mM NaCl, 1 mM ethylenediaminetetraacetic acid, 1% (v/v) Triton X-100, 10% (v/v) glycerol, 10 mM dithiothreitol, and 1 mM phenylmethanesulfonyl fluoride (3:1 [v/w] buffer/tissue). After centrifugation of slurry at 15,000× *g* for 20 min, the supernatant was used for enzyme assays. PAL activity was analyzed by monitoring the formation of *trans*-cinnamic acid from L-Phe using HPLC as described previously^8^. In brief, 100 μL of reaction mixture containing crude protein extract (4 μg), 0.1 M sodium borate (pH 8.8) and 2 mM L-Phe were incubated for 1 h at room temperature. The reaction was terminated by adding 5 μL 6 M HCl followed by centrifugation. Ten μL of the supernatant were analyzed on HPLC equipped with the Agilent Poroshell 120 EC-C18 column (2.7 μm, 150 × 3.0 mm) held at 35°C using an 8-min linear gradient of 5-70% acetonitrile in 0.1% formic acid at a flow rate of 0.4 mL min^-1^. Cinnamic acid was detected by ultraviolet absorbance at 270 nm and quantified using Open LAB CDS Chemstaion Rev. C.01.08(210) software and authentic standard.

BPBT activity was evaluated by measuring the formation of benzylbenzoate from benzoyl-CoA and benzyl alcohol as previously described^24^, while BAMT activity was determined by measuring the transfer of the methyl group of *S*-methyl-adenosyl-L-methionine to the carboxyl group of benzoic acid^55^. For BPBT activity, 100 μL of reaction mixtures containing crude protein extracts (7.5 μg), 50 mM Tris-HCl (pH 7.5), 0.1 mM EDTA, 0.5 mM benzoyl CoA, 0.5 mM benzyl alcohol were incubated at 25°C for 30 min. For BAMT activity, 200 μL of reaction mixture containing crude protein extract (80 μg), 50 mM Tris-HCl (pH 7.5), 0.5 mM EDTA, 10 mM *S*-(5’-Adenosyl)-L-methionine iodide and 1 mM sodium benzoate were incubated at 25°C for 2 h. Both BPBT and BAMT reactions were terminated by adding 5 μl of 6 M HCl, the corresponding reaction products, benzylbenzoate and methylbenzoate, were extracted twice with 200 μL hexane and 2 μL of the combined organic phases were injected into GC-MS for product analysis.

All enzyme assays were performed at an appropriate protein concentration so that reaction velocity was proportional to enzyme concentration and linear during the incubation time. Control assays included reactions (1) without protein extracts, (2) without substrate, and (3) with boiled protein extracts, and did not produce corresponding products.

### A mathematical model for VOC transport in presence of nsLTPs in the cell wall

The full description of modeling can be found in Supplementary Method 1.

### Non-linear curve fitting of the TNS displacement assay result

The binding curves of PhnsLTP3 with various VOC compounds were obtained by non-linear fitting to the one-binding-site (Equation (1)) and two-binding-site (Equation (2)) models. The equations assume that the ligand binds to multiple independent binding sites and that the binding to each site follows the law of mass action:

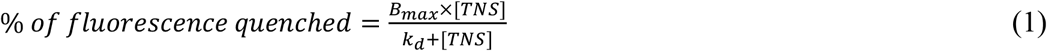

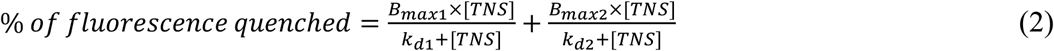

A comparison of the one-site and two-site fits was performed using the F test. The F ratio was calculated as:

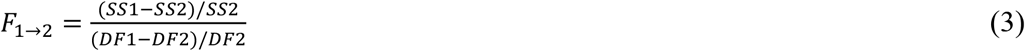

where *F*_1→2_, F ratio of the comparison between the one- and two-binding-site model; SS1, sum of squares of the data fit to the one-site model; SS2, sum of squares of the data fit to the two-site model; DF1, degrees of freedom of the data fit to the one-site model; DF2, degrees of freedom of the data fit to the two-site model.

The corresponding *P*-value was determined using the table with *DF*_*n*_ =(*DF*1− *DF*2) and *DF*_*d*_ = *DF*2. Fitting of the data to the two-site model was significantly better than to the one-site model if the *P*-value < 0.05.

### Calculation of the volatile emission factor

The volatile emission factor (VEF) was calculated as previously described^8^. In brief, the time-dependent accumulation of total volatile internal pool was estimated by linear regression. The total biosynthetic flux was estimated by the mass balance:

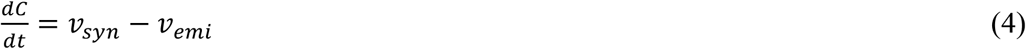

where *C*, the total volatile internal pool; *v*_*syn*_, the total volatile biosynthetic flux; *v*_*emi*_, the total volatile emission flux. The VEF is the ratio of the total emission flux over the total biosynthetic flux of the volatiles:

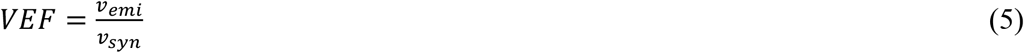

## Data availability

The data supporting the findings of this study are available within the article and its Supplementary Information files. The sequences reported in this paper have been deposited in GenBank database with the following accession numbers: PhnsLTP1, ON228071; PhnsLTP2, ON228072, and PhnsLTP3, ON228073. *Petunia axillaris* genomic data were obtained from http://solgenomics.net using *Petunia axillaris* v1.6.2 genome database.

## Acknowledgments

This work was supported by grant from the National Science Foundation IOS-1655438 to N.D. and J.A.M., by the USDA National Institute of Food and Agriculture Hatch Project number 177845 to ND, and by a United States-Israel Binational Agricultural Research and Development Postdoctoral Fellowship FI-588-2019 to I.M. The authors acknowledge the use of the Zeiss LSM-880 confocal microscope for collection of microscopy images at the imaging facilities of the Bindley Bioscience Center, a core facility of the NIH-funded Indiana Clinical and Translational Sciences Institute. We thank Ryan Benke for a generation of *PhnsLTP1*-RNAi construct.

## Author Contributions

I.M., J.A.M., and N.D. conceived the project and designed the research; P.L. performed phylogenetic analysis, water loss determination, enzyme activity assays, propidium iodide staining experiments, expression analysis, and feeding experiments. I.M. generated *PhnsLTP1-, PhnsLTP2-, PhnsLTP3-* and *PhnsLTP1*+*PhnsLTP3-*RNAi lines, performed expression analysis, and metabolic profiling of transgenic plants. M-L.S. produced recombinant PhnsLTP3 and analyzed its binding activity, performed modeling, and estimated VEF. P.L. and J.H.L. performed wax profiling and VOC distribution. J.H.L. performed toluidine staining experiments, expression analysis, and metabolic profiling at earlier time points. X-Q.H. analyzed *PhnsLTPs* expression in RNA-seq datasets. I.M., P.L., and X-Q.H. analyzed the subcellular localization of PhnsLTPs. P.L., I.M, M-L.S., J.H.L., X-Q.H., J.A.M., and N.D. analyzed data; N.D., P.L., X-Q.H., and J.A.M. wrote the manuscript with contributions from all authors. All authors read and edited the manuscript.

