## Supplementary material for "Emission of floral volatiles is facilitated by cell-wall non-specific lipid transfer proteins": Supplementary Material Final ND.pdf

Liao *et al.*

### Supplementary Method 1

**A mathematical model for VOC transport in presence of nsLTPs in the cell wall.** The extent of the enhanced transport was estimated using a steady-state facilitated transport model[1,2]. A set of reaction-diffusion equations were used to describe the facilitated transport by nsLTPs in the cell wall. The equations for the facilitated transport of nsLTPs are analogous to that of $O_2$ transport by myoglobin, which is the carrier protein in muscles that facilitates transport in the aqueous environment[3]. Considering the simplest reaction between the protein and its ligand with one-to-one binding stoichiometry:

$$VOC + nsLTP \overset{k_d}{\leftrightarrow} nsLTP \cdot VOC \quad (1)$$

$$k_d = \frac{k_r}{k_f} = \frac{S \cdot E}{C} \quad (2)$$

where $S$, concentration of the free ligands (VOCs); $E$, concentration of the free carrier proteins (nsLTPs); $C$, concentration of the protein-ligand ($nsLTP \cdot VOC$) complex; $k_f$, rate constant of the forward reaction; $k_r$, rate constant of the reverse reaction.

Assuming the system is under steady-state and the diffusion coefficient is constant, the reaction-diffusion equations can be written for the VOCs, nsLTPs (the carriers), and the $nsLTP \cdot VOC$ complex across the cell wall of length L:

$$D_S \frac{\partial^2 S}{\partial x^2} = k_f SE - k_r C \quad (3)$$

$$D_E \frac{\partial^2 E}{\partial x^2} = k_f SE - k_r C \quad (4)$$

$$D_C \frac{\partial^2 C}{\partial x^2} = -k_f SE + k_r C \quad (5)$$

where $D_S$, diffusion coefficient of the free VOCs; $D_E$, diffusion coefficient of the free nsLTPs; $D_C$, diffusion coefficient of the $\mathrm{nsLTP} \cdot \mathrm{VOC}$ complex.

The boundary conditions on VOC concentrations are considered to be constant based on the solubility-diffusion model (Equation (6)) and no release of nsLTPs and the $nsLTP \cdot VOC$ complex (Equation (7)) because they are nonvolatile and thus are contained in the cell wall:

$$S|_{x=0} = S_0 \quad , \quad S|_{x=L} = S_L = {C_{Cut}}/{K_{c/w}} \quad (6)$$

$$\frac{dE}{dx}|_{x=0} = \frac{dC}{dx}|_{x=0} = 0 \quad , \quad \frac{dE}{dx}|_{x=L} = \frac{dC}{dx}|_{x=L} = 0 \tag{7}$$

where $C_{Cut}$, VOC concentrations in the cuticle; $K_{c/w}$, cuticle-water partition coefficient of VOC.

It is worth mentioning that for the boundary condition (Equation (6)), unlike $S_L$ which can be determined from $C_{Cut}$ and $K_{c/w}$ based on the equilibrium assumption between the cuticle and cell wall phases, the $S_0$ value cannot be calculated in the same manner because of the plasma membrane-localized ABC transporters. ABC transporters will continuously pump VOCs out of the plasma membrane thus driving the condition on the left boundary of the cell wall layer away from equilibrium with the plasma membrane. Lacking the enzyme kinetic information of the ABC transporters, the VOC levels built up by the transporters in the cell wall ($S_0$) is hard to be estimated for simulation. Since the facilitated diffusion relies on the concentration gradient, as a first approach, a range of $S_0$, which was arbitrarily set to be larger than $S_L$, was implemented to study the effect of different levels of concentration gradient on the facilitation of flux.

The equations can be analytically solved[1] and obtain the total flux:

$$J = \frac{D_S}{L}(S_0 - S_L) + \frac{D_C}{L}E_T\left(\frac{S_0}{S_0+k_d} - \frac{S_L}{S_L+k_d}\right) = \frac{D_S}{L}(S_0 - S_L)[1 + \frac{D_C}{D_S}\frac{E_T k_d}{(S_0+k_d)(S_L+k_d)}] \tag{8}$$

where $E_T$, total concentration of the carrier protein (LTP) in the cell wall, and $J = J_{emi} = J_{Free} + J_{Facilitated} = J_{syn}$ (see Supplementary Figure 18). From Equation (8), a dimensionless parameter, the extent of the flux enhancement (E.o.F), can be defined as the ratio of the total flux over VOC flux by only free diffusion can be written as equation (9):

$$\mathrm{E.o.F} = 1 + \frac{D_C}{D_S}\frac{E_T k_d}{(S_0+k_d)(S_L+k_d)} \tag{9}$$

Considering the scenario that the carrier protein (nsLTP) has two independent binding sites for ligands:

$$VOC + nsLTP_1 \overset{k_{d1}}{\leftrightarrow} nsLTP_1 \cdot VOC \tag{10}$$

$$VOC + nsLTP_2 \overset{k_{d2}}{\leftrightarrow} nsLTP_2 \cdot VOC \tag{11}$$

$$k_{d1} = \frac{k_{r,1}}{k_{f,1}} = \frac{S \cdot E_1}{C_1} \tag{12}$$

$$k_{d2} = \frac{k_{r,2}}{k_{f,2}} = \frac{S \cdot E_2}{C_2} \tag{13}$$

where $E_1$, concentration of the free carrier protein with the number 1 binding site vacant (nsLTP1); $E_2$, concentration of the free carrier protein with the number 2 binding site vacant (nsLTP2); $C_1$, concentration of the $nsLTP_1 \cdot VOC$ complex; $C_2$, concentration of the $nsLTP_2 \cdot VOC$ complex; $k_{f,1}$, rate constant of the first forward reaction; $k_{r,1}$, rate constant of the first reverse reaction; $k_{f,2}$, rate constant of the second forward reaction; $k_{r,2}$, rate constant of the second reverse reaction.

The equations can be solved as detailed previously[4], and the total VOC flux can be expressed as:

$$J = \frac{D_S}{L}(S_0 - S_L) + \frac{D_{C_1}}{L}(C_{1,0} - C_{1,L}) + \frac{D_{C_2}}{L}(C_{2,0} - C_{2,L}) \tag{14}$$

where $C_{1,0}$, $nsLTP_1 \cdot VOC$ concentration on the left boundary; $C_{1,L}$, $nsLTP_1 \cdot VOC$ concentration on the right boundary; $C_{2,0}$, $nsLTP_2 \cdot VOC$ concentration on the left boundary; $C_{2,L}$, $nsLTP_2 \cdot VOC$ concentration on the right boundary; $D_{C_1}$, diffusion coefficient of the $nsLTP_1 \cdot VOC$ complex; $D_{C_2}$, diffusion coefficient of the $nsLTP_2 \cdot VOC$ complex.

Assuming that VOCs and nsLTPs are at quasi-steady state throughout the cell wall, $nsLTP_1 \cdot VOC$ and $nsLTP_1 \cdot VOC$ concentrations can be written as:

$$C_1 = \frac{S}{S + k_{d1}} E_T \tag{15}$$

$$C_2 = \frac{S}{S + k_{d2}} E_T \tag{16}$$

Equation (15) and Equation (16) can be substituted into Equation (14), and the corresponding extent of the flux enhancement (E.o.F) can be defined as previously stated. The values of the parameters used are provided in Supplementary Table 6.

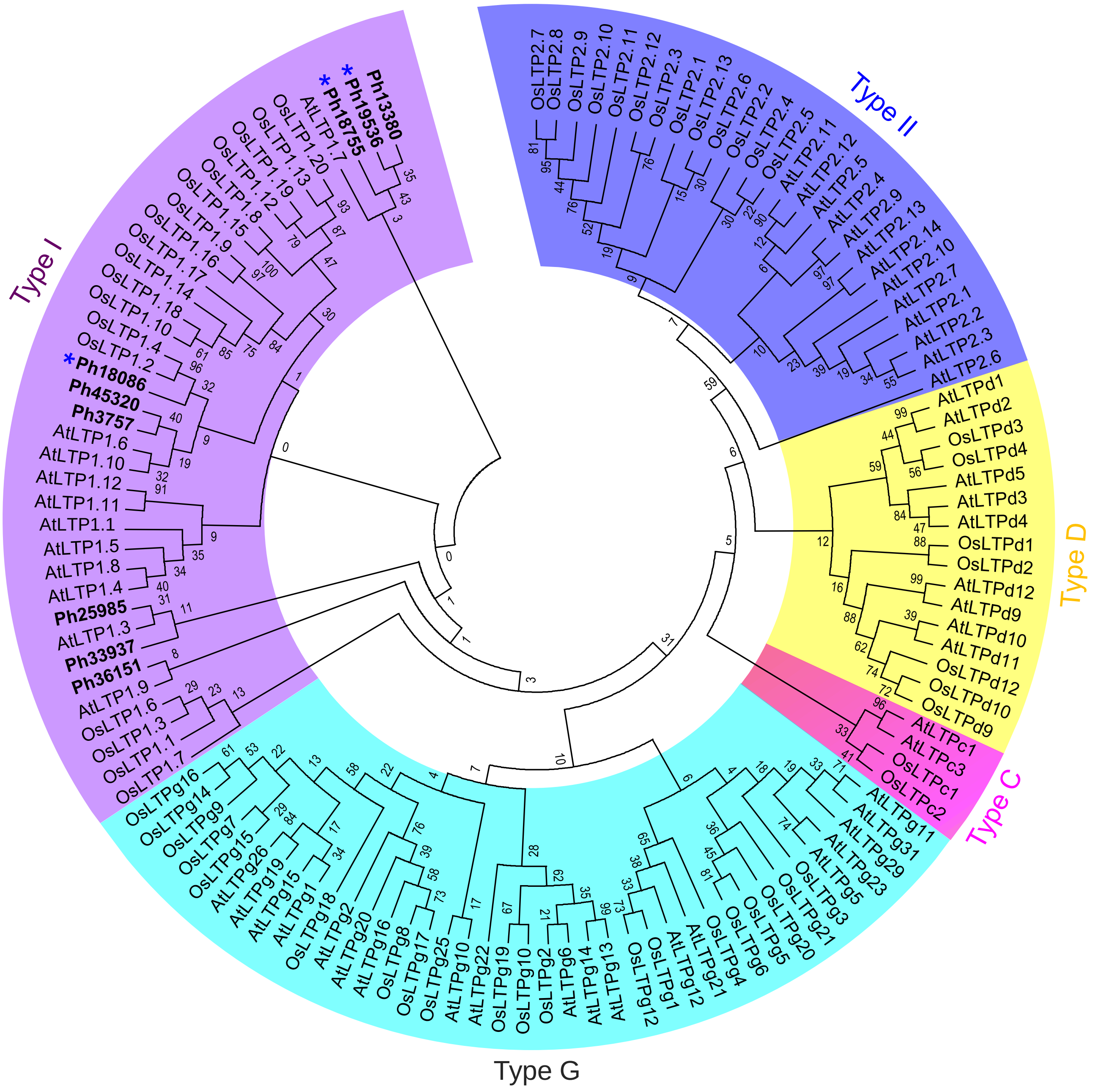


**Supplementary Figure 1 | Phylogenetic tree of nsLTP sequences from *Petunia hybrida*, *Arabidopsis thaliana*, and *Oryza sativa*.** The phylogenetic tree was built by the maximum-likelihood method in the MEGA7 package from alignment of mature proteins using MUSCLE algorithm. Different colors highlight different nsLTP subfamilies: purple, Type I; blue, Type II; pink, Type C; yellow, Type D; and green, Type G. All *P. hybrida* (Ph) LTPs are shown in bold and Ph19536, Ph18755 and Ph18086 corresponding to PhLTP1, PhLTP2 and PhLTP3, respectively, are marked with asterisks. All the sequences from *A. thaliana* (At) and *O. sativa* (Os) were obtained from Edstam et al. (2011). The percentage of replicate trees in which the associated taxa clustered together in the bootstrap test (500 replicates) are shown next to the branches.

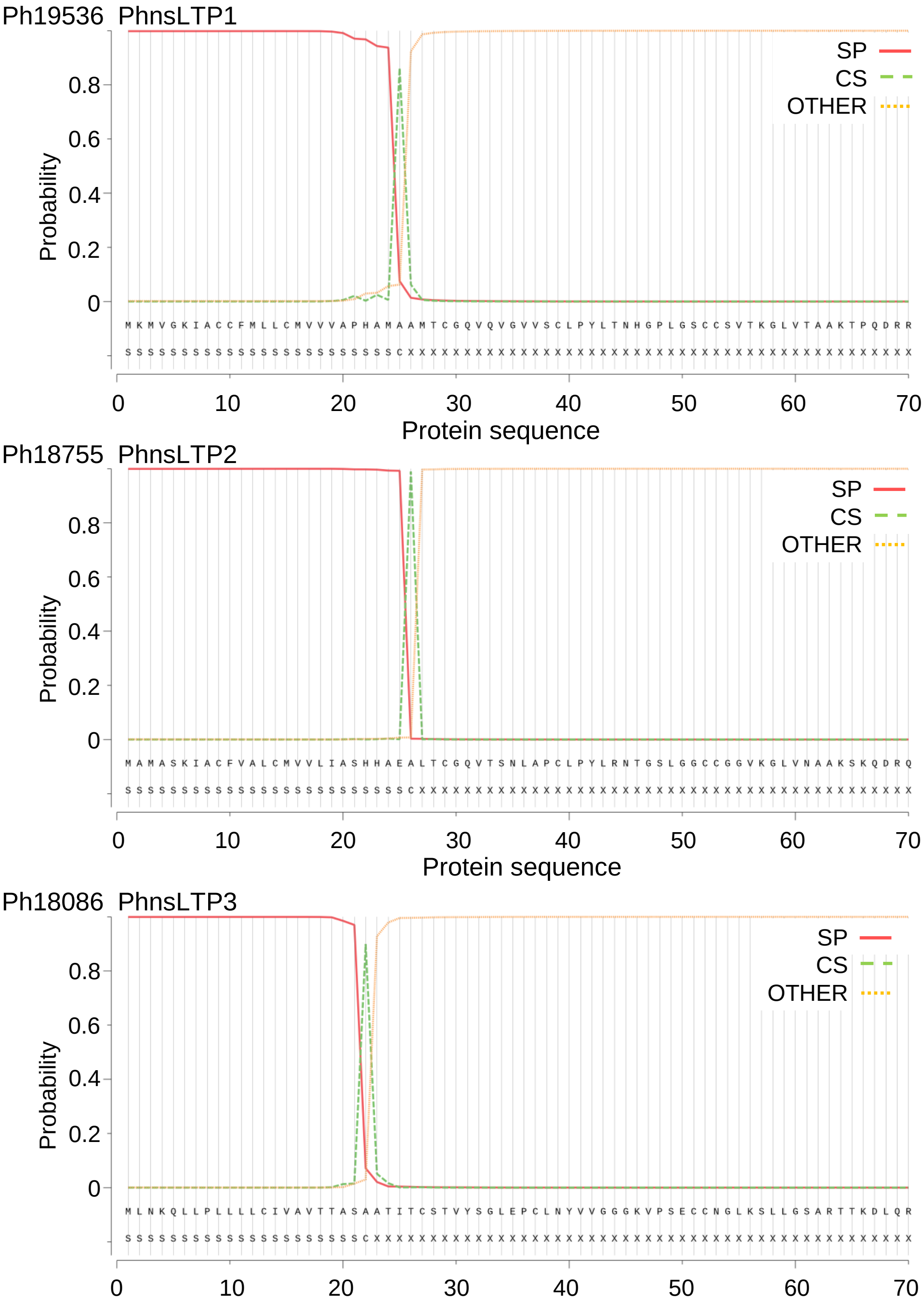


**Supplementary Figure 2 | Prediction of signal peptide in PhnsLTP1, PhnsLTP2 and PhnsLTP3.** Analysis was performed using SignalP-5.0. SP, secretory signal peptide; CS: the cleavage site; OTHER: the probability that the sequence does not have any kind of signal peptide.

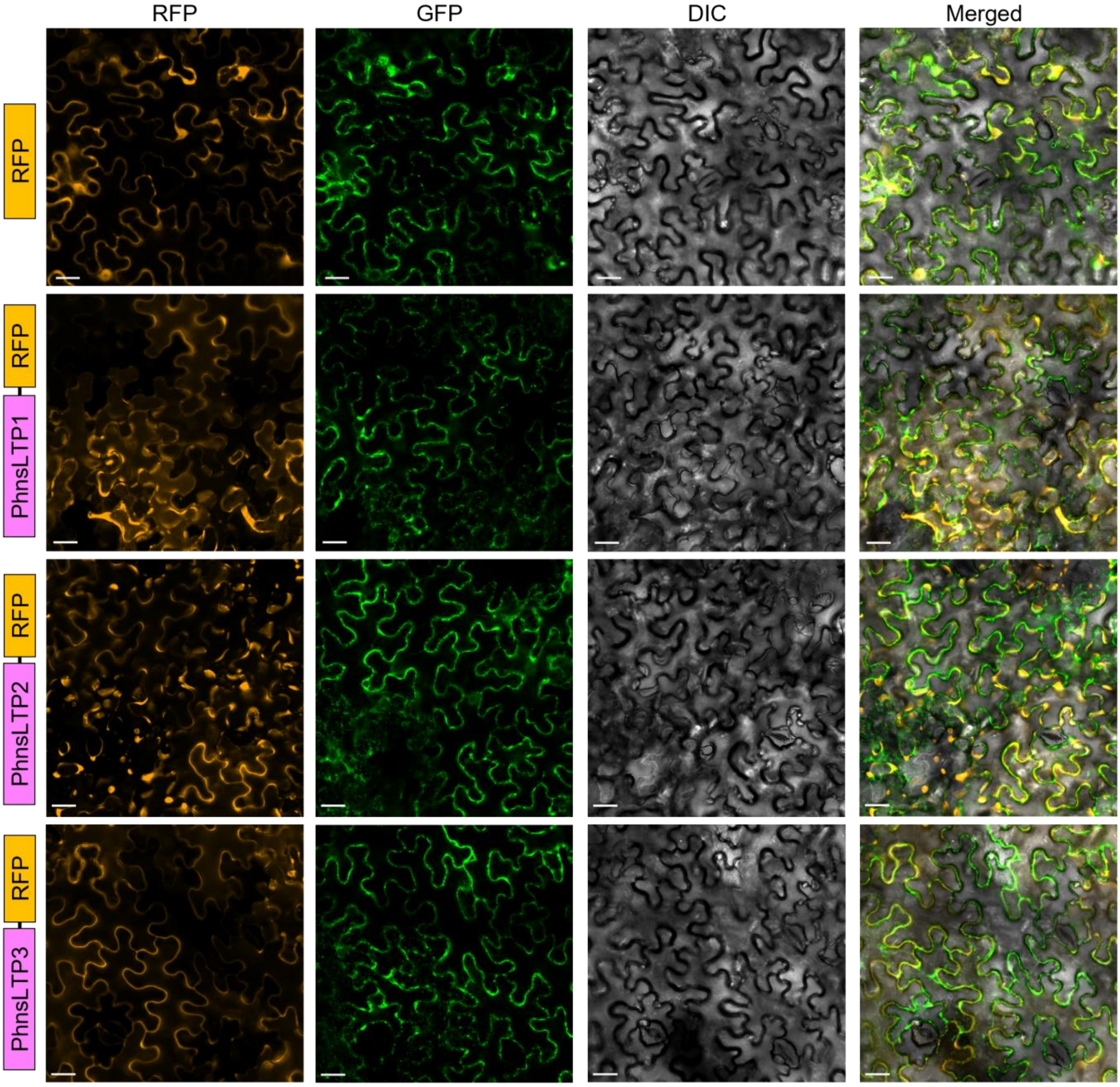


**Supplementary Figure 3 | Subcellular localization of PhnsLTP1, PhnsLTP2 and PhnsLTP3.** PhnsLTP1, PhnsLTP2, and PhnsLTP3 fusion constructs were expressed in *N. benthamiana* leaves and their corresponding transient expression was detected by confocal laser scanning microscopy. "RFP" panels (orange) represent signals of PhnsLTP1, PhnsLTP2, and PhnsLTP3 fused fluorescence proteins; the "GFP" panels (green) represent signals of the plasma membrane marker protein, AtPIP1:GFP; the DIC panels (dark grey) show images of differential-interference-contrast (DIC) microscopy; and the "Merged" panels show merged RFP, GFP and DIC signals. Schematic diagrams of the PhnsLTP-RFP fusion proteins for each experiment are shown on the left. Images were captured using 20× objective lens. Scale bars are 10 µm. The experiments were repeated three times with similar results.

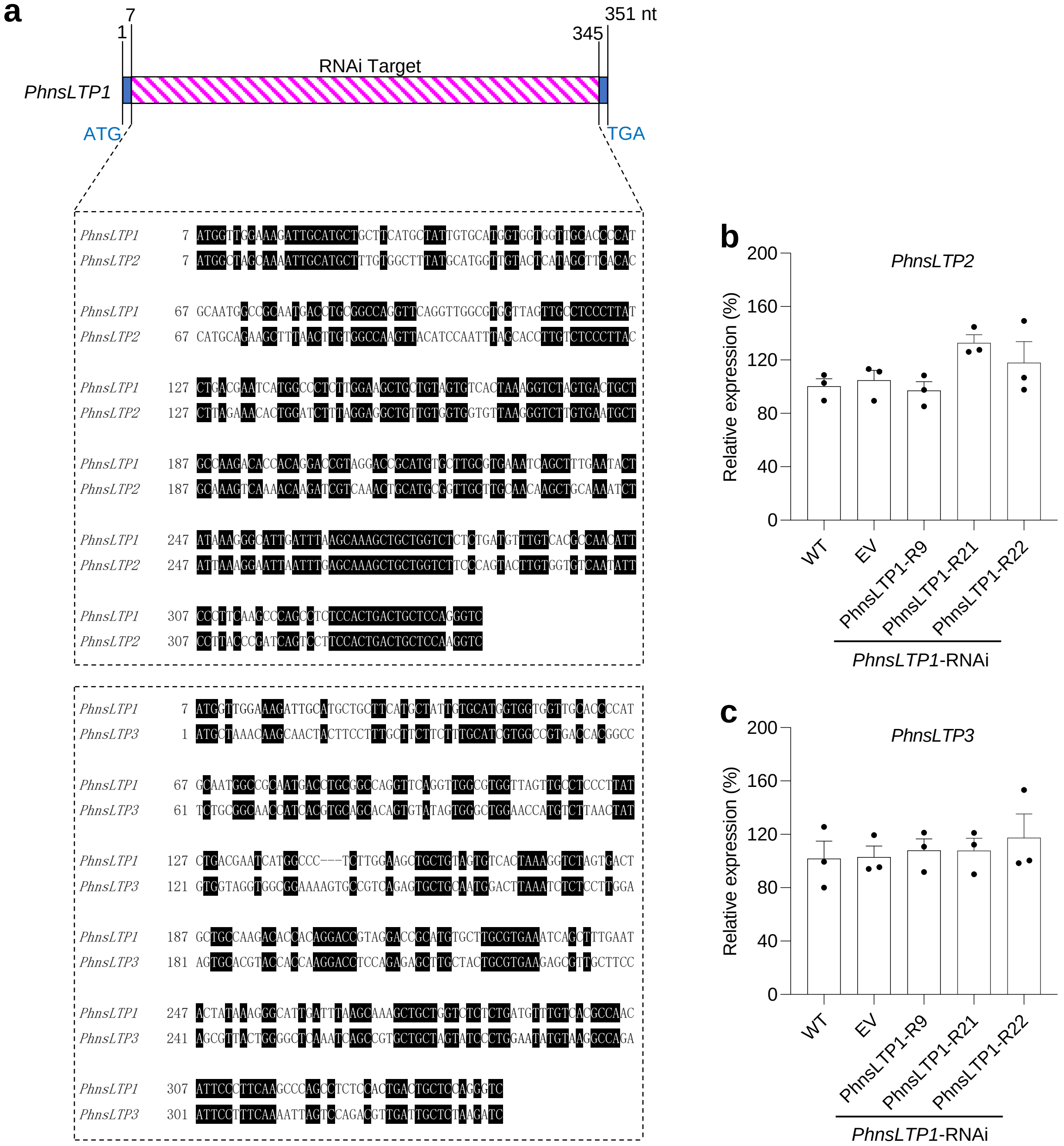


**Supplementary Figure 4 | Effect of *PhnsLTP1* dsRNA trigger on expression of *PhnsLTP2* and *PhnsLTP3*. a,** Diagram showing location of a region used to generate the *PhnsLTP1* dsRNA trigger and a nucleotide alignment of the *PhnsLTP1* region targeted by the RNAi trigger with *PhnsLTP2* and *PhnsLTP3*. This *PhnsLTP1* region is 66% and 52% identical to the corresponding regions of *PhnsLTP2* and *PhnsLTP3*. No stretches greater than 17 and 8 uninterrupted nucleotides are shared between *PhnsLTP1* and *PhnsLTP2* and between *PhnsLTP1* and *PhnsLTP3*, respectively. Thus, the *PhnsLTP1*-RNAi construct specifically triggers *PhnsLTP1* downregulation. Effect of *PhnsLTP1* downregulation on expression of *PhnsLTP2* (**b**) and *PhnsLTP3* (**c**) in *PhnsLTP1*-RNAi lines. Transcript levels were determined by qRT-PCR with gene specific primers in 2-day-old flowers from wild type (WT), empty vector control (EV) and 3 independent *PhnsLTP1*-RNAi lines (R9, R21 and R22) harvested at 15:00 h and presented relative to the corresponding levels in WT, set as 100%. Data are means ± S.E. (n = 3 biological replicates). *P* values in (**b**) and (**c**) were determined by one-way ANOVA with the Dunnett's multiple comparisons test relative to the WT and EV controls.

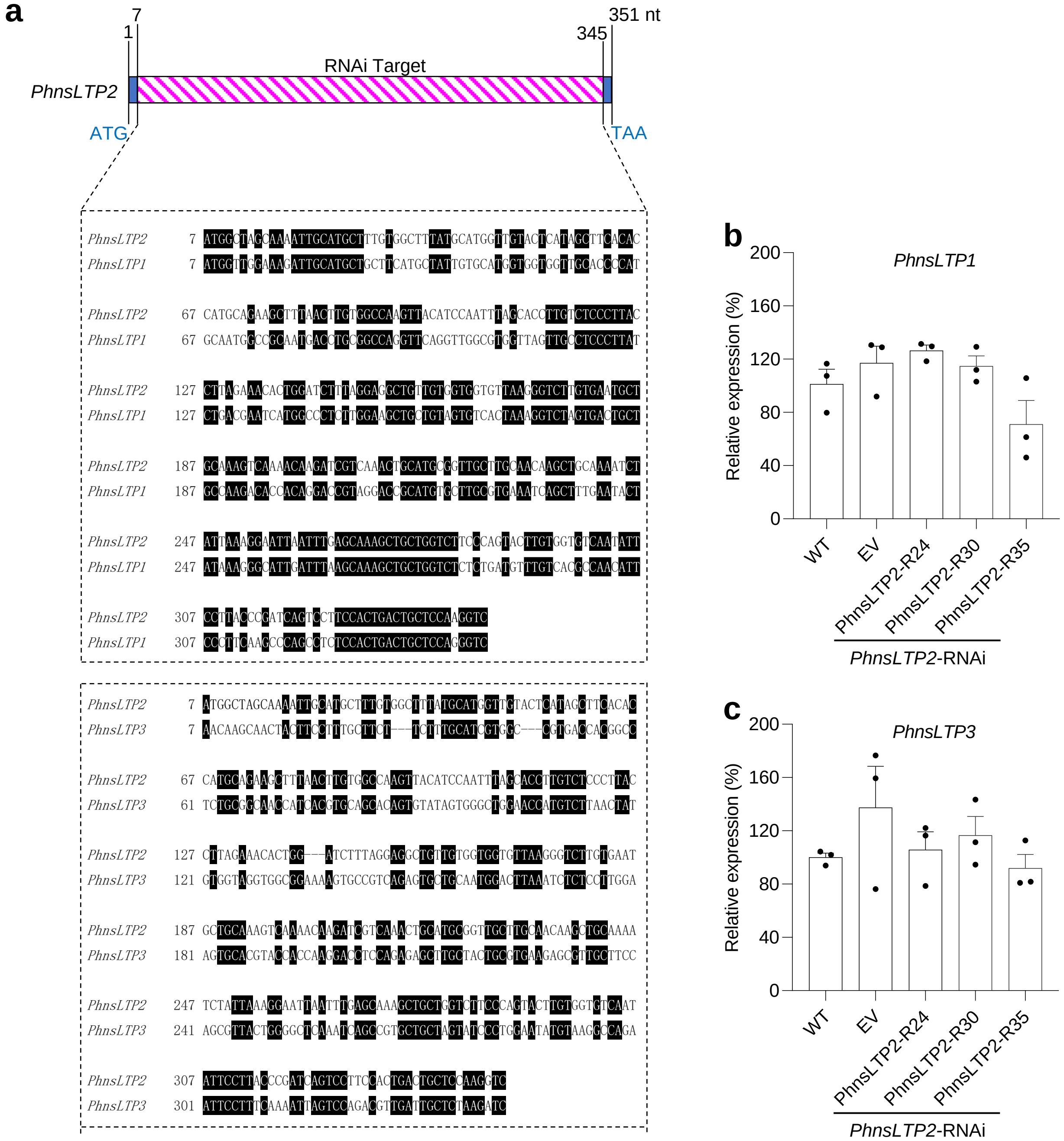


**Supplementary Figure 5 | Effect of *PhnsLTP2* dsRNA trigger on expression of *PhnsLTP1* and *PhnsLTP3*. a,** Diagram showing location of a region used to generate the *PhnsLTP2* dsRNA trigger and a nucleotide alignment of the *PhnsLTP2* region targeted by the RNAi trigger with *PhnsLTP1* and *PhnsLTP3*. This *PhnsLTP2* region is 66% and 50% identical to the corresponding regions of *PhnsLTP1* and *PhnsLTP3*. No stretches greater than 17 and 7 uninterrupted nucleotides are shared between *PhnsLTP2* and *PhnsLTP1* and between *PhnsLTP2* and *PhnsLTP3*, respectively. Thus, the *PhnsLTP2*-RNAi construct specifically triggers *PhnsLTP2* downregulation. Effect of *PhnsLTP2* downregulation on expression of *PhnsLTP1* (**b**) and *PhnsLTP3* (**c**) in *PhnsLTP2*-RNAi lines. Transcript levels were determined by qRT-PCR with gene specific primers in 2-day-old flowers from wild type (WT), empty vector control (EV) and 3 independent *PhnsLTP2*-RNAi lines (R24, R30 and R35) harvested at 15:00 h and presented relative to the corresponding levels in WT, set as 100%. Data are means ± S.E. (n = 3 biological replicates). *P* values in (**b**) and (**c**) were determined by one-way ANOVA with the Dunnett's multiple comparisons test relative to the WT and EV controls.

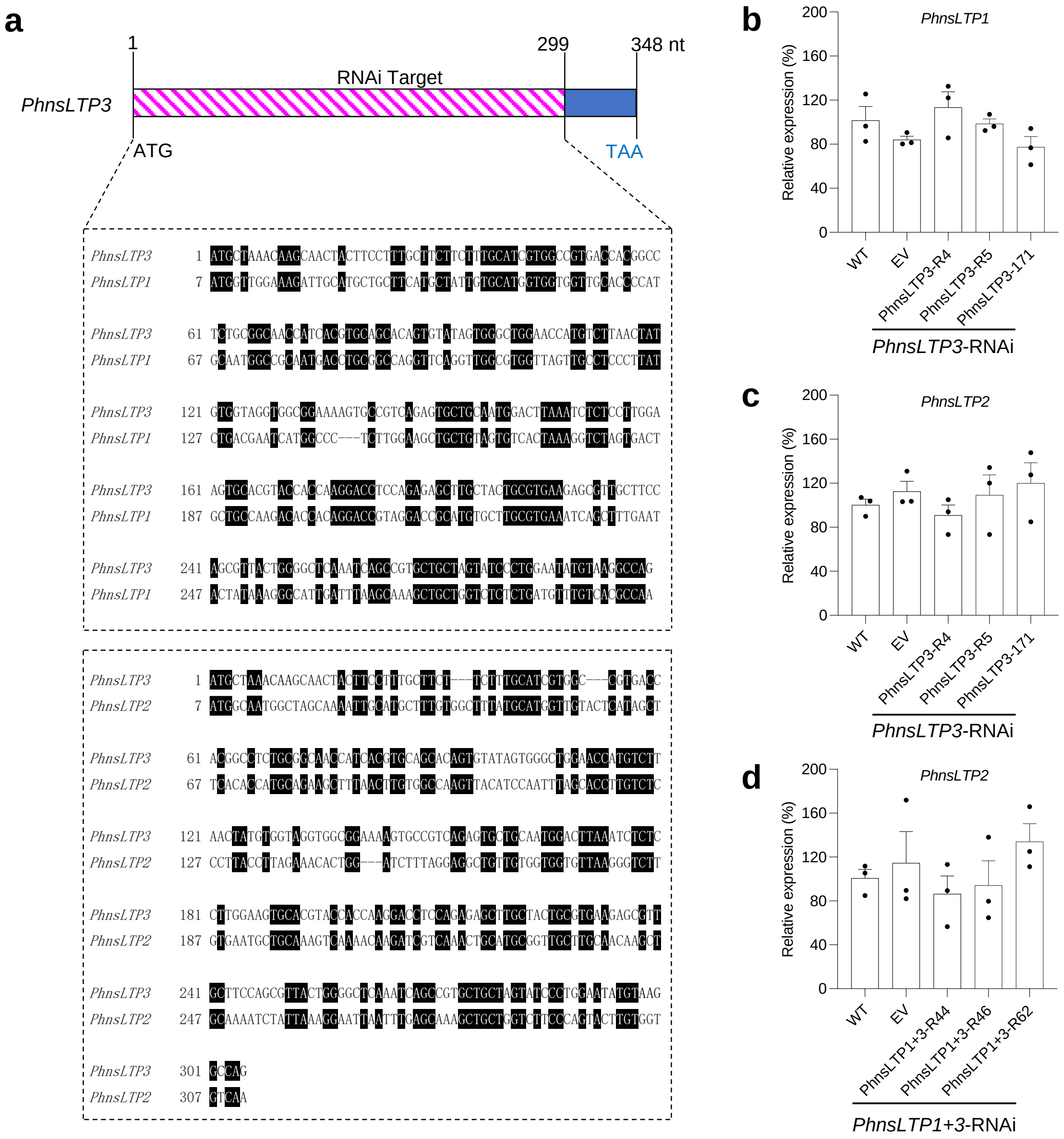


**Supplementary Figure 6 | Effect of *PhnsLTP3* and *PhnsLTP1*+ *PhnsLTP3* dsRNA triggers on expression of *PhnsLTP1* and *PhnsLTP2*. a,** Diagram showing location of a region used to generate the *PhnsLTP3* dsRNA trigger and a nucleotide alignment of the *PhnsLTP3* region targeted by the RNAi trigger with *PhnsLTP1* and *PhnsLTP2*. This *PhnsLTP3* region is 51% and 47% identical to the corresponding regions of *PhnsLTP1* and *PhnsLTP2*. No stretches greater than 8 and 6 uninterrupted nucleotides are shared between *PhnsLTP3* and *PhnsLTP1* and between *PhnsLTP3* and *PhnsLTP2*, respectively. Thus, the *PhnsLTP3*-RNAi construct specifically triggers *PhnsLTP3* downregulation. Effect of *PhnsLTP3* downregulation on expression of *PhnsLTP1* (**b**) and *PhnsLTP2* (**c**) in *PhnsLTP3*-RNAi lines. **d**, Effect of simultaneous *PhnsLTP1* and *PhnsLTP3* downregulation on *PhnsLTP2* expression in *PhnsLTP1*+*PhnsLTP3*-RNAi lines. Transcript levels in (**b** - **d**) were determined by qRT-PCR with gene specific primers in 2-day-old flowers from wild type (WT), empty vector control (EV), 3 independent *PhnsLTP3*-RNAi lines (R4, R5 and 171) (**b**, **c**) and 3 independent *PhnsLTP1*+*PhnsLTP3*-RNAi lines (R44, R46 and R62) (**d**) harvested at 15:00 h and presented relative to the corresponding levels in WT, set as 100%. Data are means ± S.E. (n = 3 biological replicates). *P* values in (**b** - **d**) were determined by one-way ANOVA with the Dunnett's multiple comparisons test relative to the WT and EV controls.

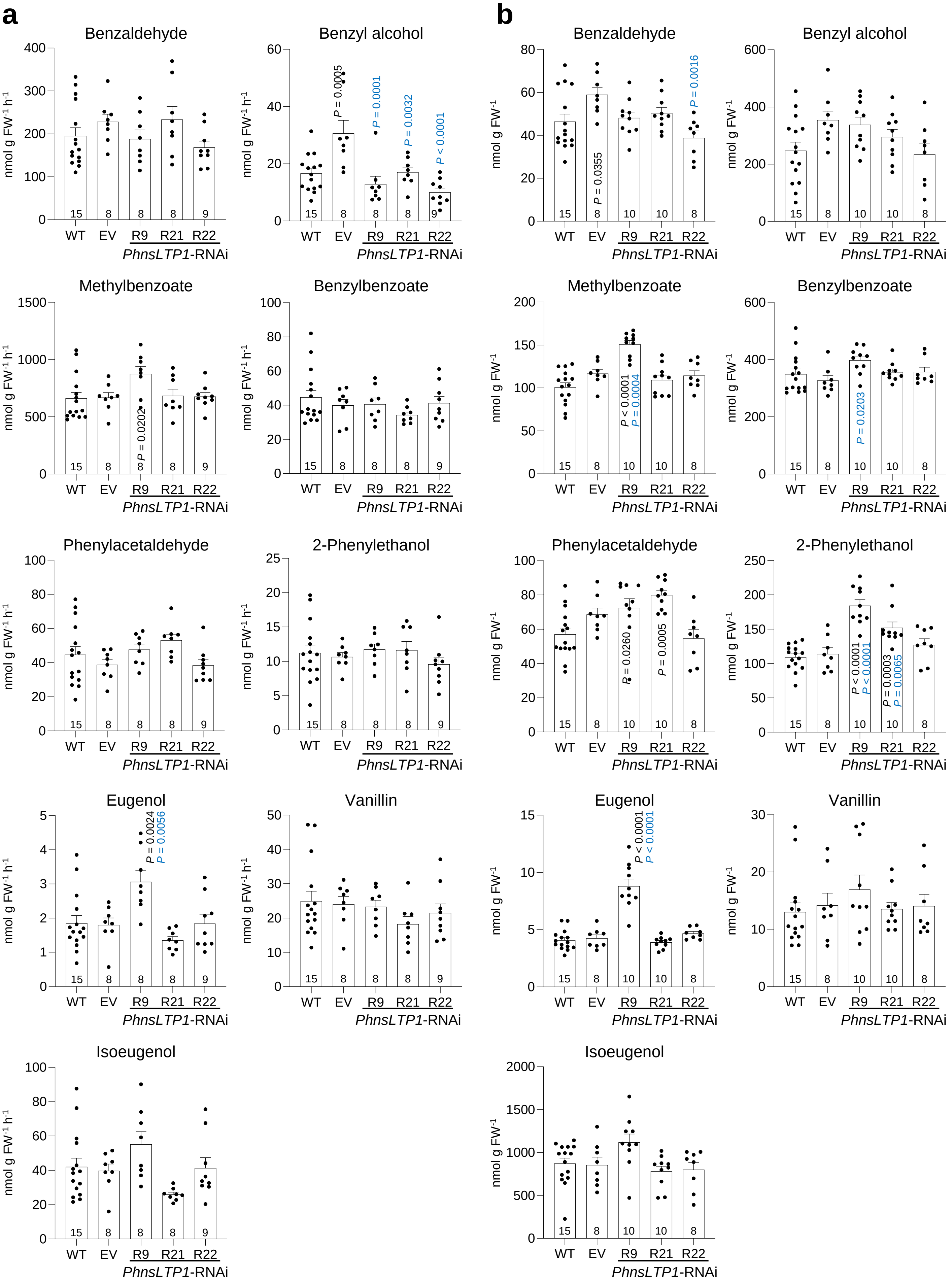
a
b
Benzaldehyde
Benzyl alcohol
Methylbenzoate
Benzylbenzoate
Phenylacetaldehyde
2-Phenylethanol
Eugenol
Vanillin
Isoeugenol
nmol g FW-1 h-1
nmol g FW-1
WT
EV
R9
R21
R22
PhnsLTP1-RNAi
P = 0.0005
P = 0.0001
P = 0.0032
P < 0.0001
P = 0.0355
P = 0.0016
P = 0.0202
P < 0.0001
P = 0.0004
P = 0.0203
P = 0.0260
P = 0.0005
P < 0.0001
P < 0.0001
P = 0.0003
P = 0.0065
P = 0.0024
P = 0.0056
P < 0.0001
P < 0.0001

**Supplementary Figure 7 | Effect of *PhnsLTP1* downregulation on emission and total pools of individual benzenoid/phenylpropanoid volatiles in 2-day-old petunia flowers.** Emission **(a)** and total pools **(b)** of individual VOCs are shown for wild type (WT), empty vector control (EV) and three independent *PhnsLTP1-RNAi* lines (R9, R21, and R22). Data are means ± S.E. Numbers at the bottoms of the columns represent the numbers of biological replicates. *P* values were determined by one-way ANOVA with the Dunnett's multiple comparisons test relative to the WT (black) and EV (blue) controls. FW, fresh weight.

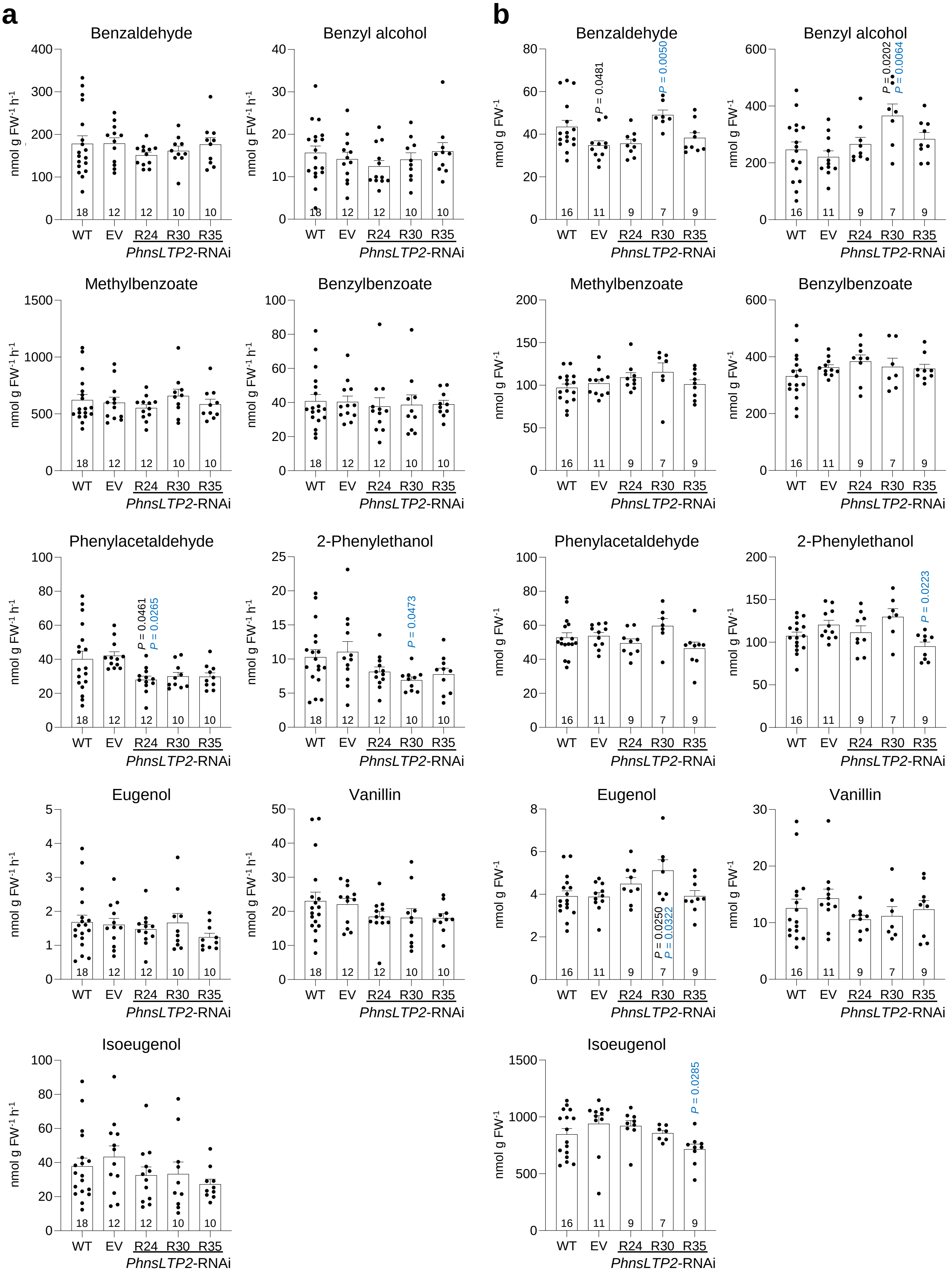
a
b
Benzaldehyde
Benzyl alcohol
Methylbenzoate
Benzylbenzoate
Phenylacetaldehyde
2-Phenylethanol
Eugenol
Vanillin
Isoeugenol
nmol g FW-1 h-1
nmol g FW-1
WT
EV
R24
R30
R35
PhnsLTP2-RNAi
P = 0.0481
P = 0.0050
P = 0.0202
P = 0.0064
P = 0.0461
P = 0.0265
P = 0.0473
P = 0.0223
P = 0.0250
P = 0.0322
P = 0.0285

**Supplementary Figure 8 | Effect of *PhnsLTP2* downregulation on emission and total pools of individual benzenoid/phenylpropanoid volatiles in 2-day-old petunia flowers.** Emission **(a)** and total pools **(b)** of individual VOCs are shown for wild type (WT), empty vector control (EV) and three independent *PhnsLTP2-RNAi* lines (R24, R30, and R35). Data are means ± S.E. Numbers at the bottoms of the columns represent the numbers of biological replicates. *P* values were determined by one-way ANOVA with the Dunnett's multiple comparisons test relative to the WT (black) and EV (blue) controls. FW, fresh weight.

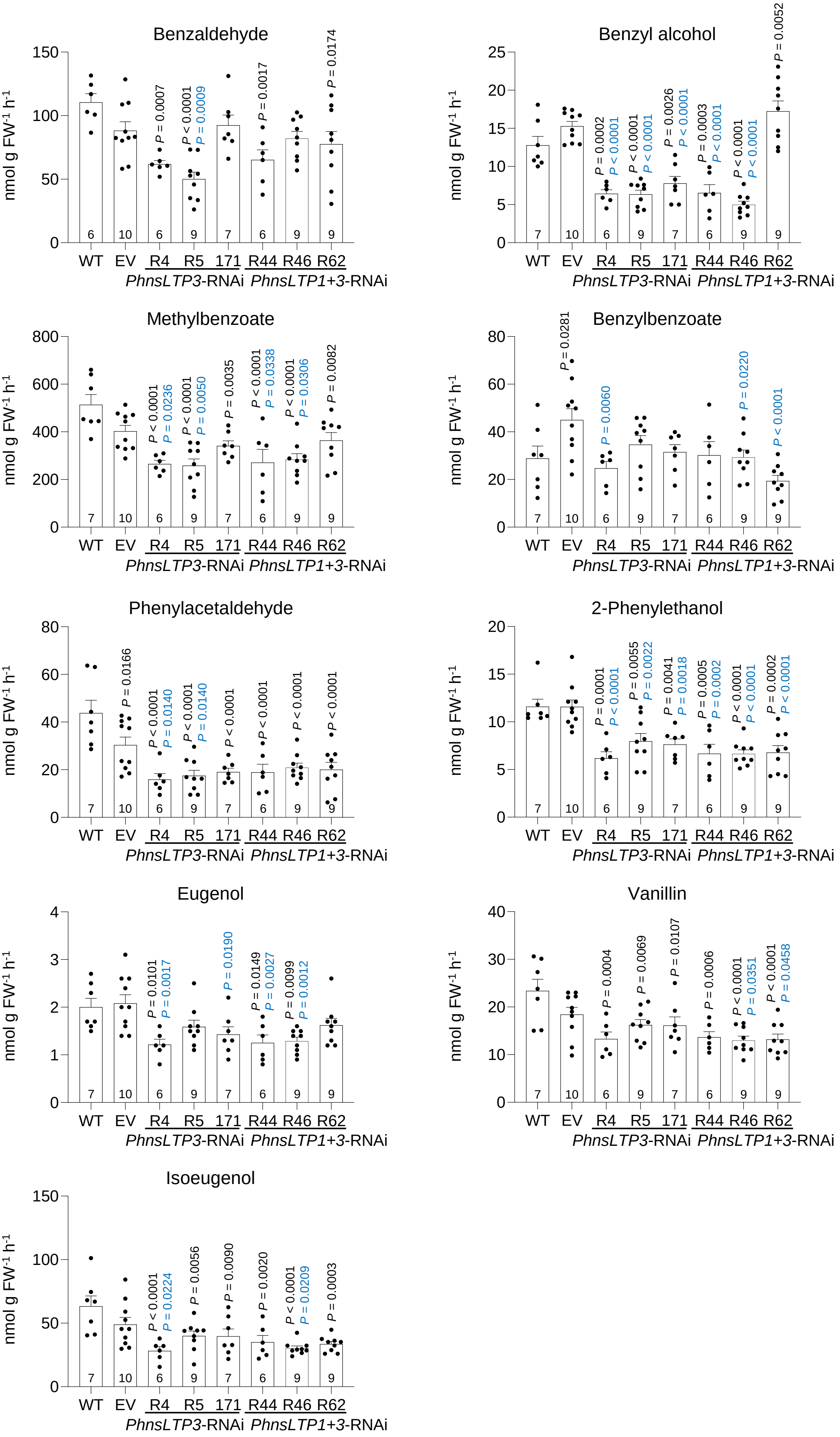

Benzaldehyde
nmol g FW-1 h-1
P = 0.0007
P < 0.0001
P = 0.0009
P = 0.0017
P = 0.0174
6 10 6 9 7 6 9 9
WT EV R4 R5 171 R44 R46 R62
PhnsLTP3-RNAi PhnsLTP1+3-RNAi
Benzyl alcohol
nmol g FW-1 h-1
P = 0.0002
P < 0.0001
P < 0.0001
P < 0.0001
P = 0.0026
P < 0.0001
P = 0.0003
P < 0.0001
P < 0.0001
P < 0.0001
P = 0.0052
7 10 6 9 7 6 9 9
WT EV R4 R5 171 R44 R46 R62
PhnsLTP3-RNAi PhnsLTP1+3-RNAi
Methylbenzoate
nmol g FW-1 h-1
P < 0.0001
P = 0.0236
P < 0.0001
P = 0.0050
P = 0.0035
P < 0.0001
P = 0.0338
P < 0.0001
P = 0.0306
P = 0.0082
7 10 6 9 7 6 9 9
WT EV R4 R5 171 R44 R46 R62
PhnsLTP3-RNAi PhnsLTP1+3-RNAi
Benzylbenzoate
nmol g FW-1 h-1
P = 0.0281
P = 0.0060
P = 0.0220
P < 0.0001
7 10 6 9 7 6 9 9
WT EV R4 R5 171 R44 R46 R62
PhnsLTP3-RNAi PhnsLTP1+3-RNAi
Phenylacetaldehyde
nmol g FW-1 h-1
P = 0.0166
P < 0.0001
P = 0.0140
P < 0.0001
P = 0.0140
P < 0.0001
P < 0.0001
P < 0.0001
P < 0.0001
7 10 6 9 7 6 9 9
WT EV R4 R5 171 R44 R46 R62
PhnsLTP3-RNAi PhnsLTP1+3-RNAi
2-Phenylethanol
nmol g FW-1 h-1
P = 0.0001
P < 0.0001
P = 0.0055
P = 0.0022
P = 0.0041
P = 0.0018
P = 0.0005
P = 0.0002
P < 0.0001
P < 0.0001
P = 0.0002
P < 0.0001
7 10 6 9 7 6 9 9
WT EV R4 R5 171 R44 R46 R62
PhnsLTP3-RNAi PhnsLTP1+3-RNAi
Eugenol
nmol g FW-1 h-1
P = 0.0101
P = 0.0017
P = 0.0190
P = 0.0149
P = 0.0027
P = 0.0099
P = 0.0012
7 10 6 9 7 6 9 9
WT EV R4 R5 171 R44 R46 R62
PhnsLTP3-RNAi PhnsLTP1+3-RNAi
Vanillin
nmol g FW-1 h-1
P = 0.0004
P = 0.0069
P = 0.0107
P = 0.0006
P < 0.0001
P = 0.0351
P < 0.0001
P = 0.0458
7 10 6 9 7 6 9 9
WT EV R4 R5 171 R44 R46 R62
PhnsLTP3-RNAi PhnsLTP1+3-RNAi
Isoeugenol
nmol g FW-1 h-1
P < 0.0001
P = 0.0224
P = 0.0056
P = 0.0090
P = 0.0020
P < 0.0001
P = 0.0209
P = 0.0003
7 10 6 9 7 6 9 9
WT EV R4 R5 171 R44 R46 R62
PhnsLTP3-RNAi PhnsLTP1+3-RNAi

**Supplementary Figure 9 | Effect of *PhnsLTP3* and *PhnsLTP1+ PhnsLTP3* downregulation on emission of individual benzenoid/phenylpropanoid volatiles in 2-day-old petunia flowers.** Emission of individual VOCs are shown for wild type (WT), empty vector control (EV) and three independent *PhnsLTP3*-RNAi lines (R4, R5, and 171) and *PhnsLTP1+PhnsLTP3* RNAi lines (R44, R46 and R62). Data are means ± S.E. Numbers at the bottoms of the columns represent the numbers of biological replicates. *P* values were determined by one-way ANOVA with the Dunnett's multiple comparisons test relative to the WT (black) and EV (blue) controls. FW, fresh weight.

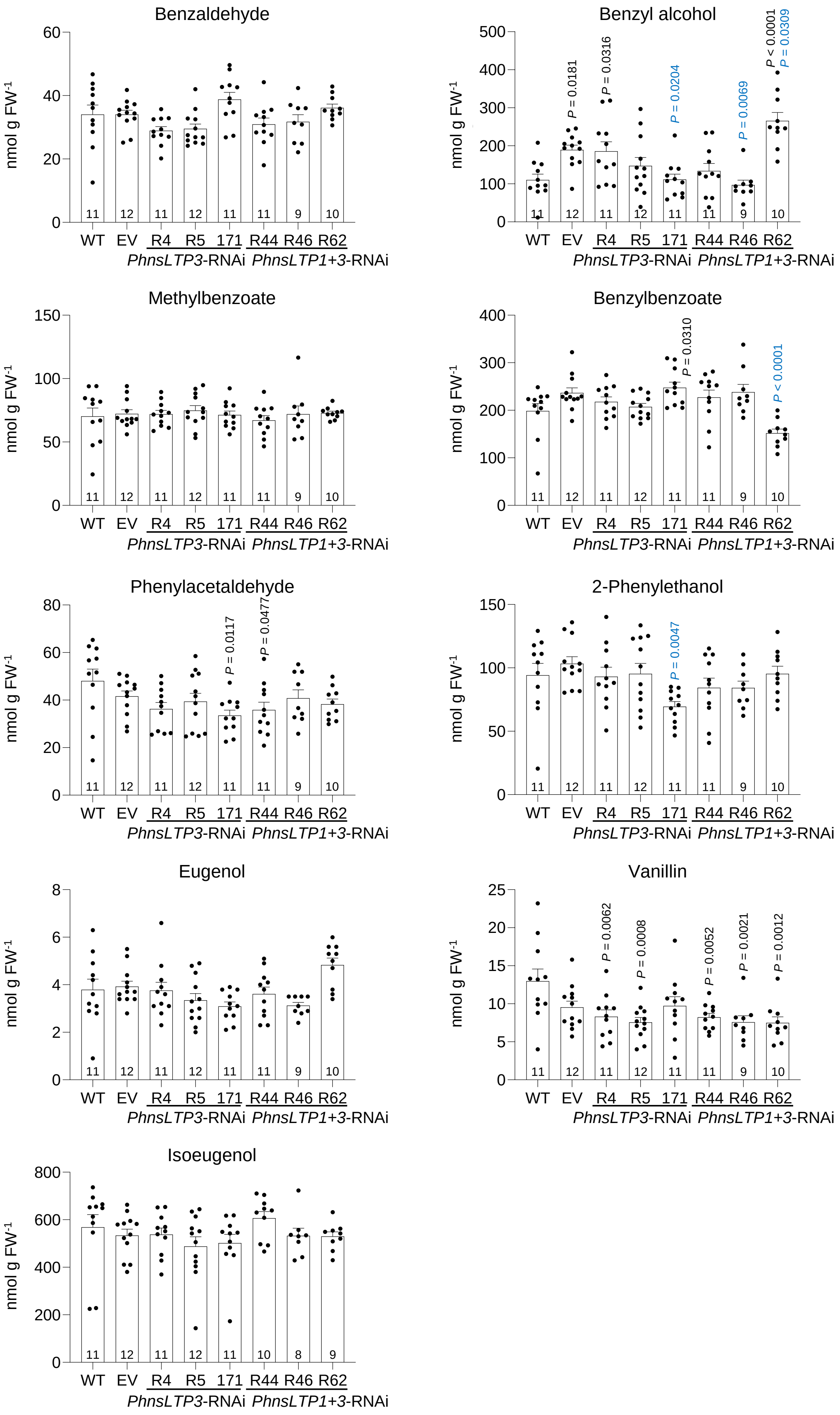
Benzaldehyde
Benzyl alcohol
P = 0.0181
P = 0.0316
P = 0.0204
P = 0.0069
P < 0.0001
P = 0.0309
Methylbenzoate
Benzylbenzoate
P = 0.0310
P < 0.0001
Phenylacetaldehyde
P = 0.0117
P = 0.0477
2-Phenylethanol
P = 0.0047
Eugenol
Vanillin
P = 0.0062
P = 0.0008
P = 0.0052
P = 0.0021
P = 0.0012
Isoeugenol
nmol g FW-1
WT EV R4 R5 171 R44 R46 R62
PhnsLTP3-RNAi PhnsLTP1+3-RNAi

**Supplementary Figure 10 | Effect of *PhnsLTP3* and *PhnsLTP1+ PhnsLTP3* downregulation on total pools of individual benzenoid/phenylpropanoid volatiles in 2-day-old petunia flowers.** Total pools of individual VOCs are shown for wild type (WT), empty vector control (EV) and three independent *PhnsLTP3-RNAi* lines (R4, R5, and 171) and *PhnsLTP1+PhnsLTP3* RNAi lines (R44, R46 and R62). Data are means ± S.E. Numbers at the bottoms of the columns represent the numbers of biological replicates. *P* values were determined by one-way ANOVA with the Dunnett's multiple comparisons test relative to the WT (black) and EV (blue) controls. FW, fresh weight.

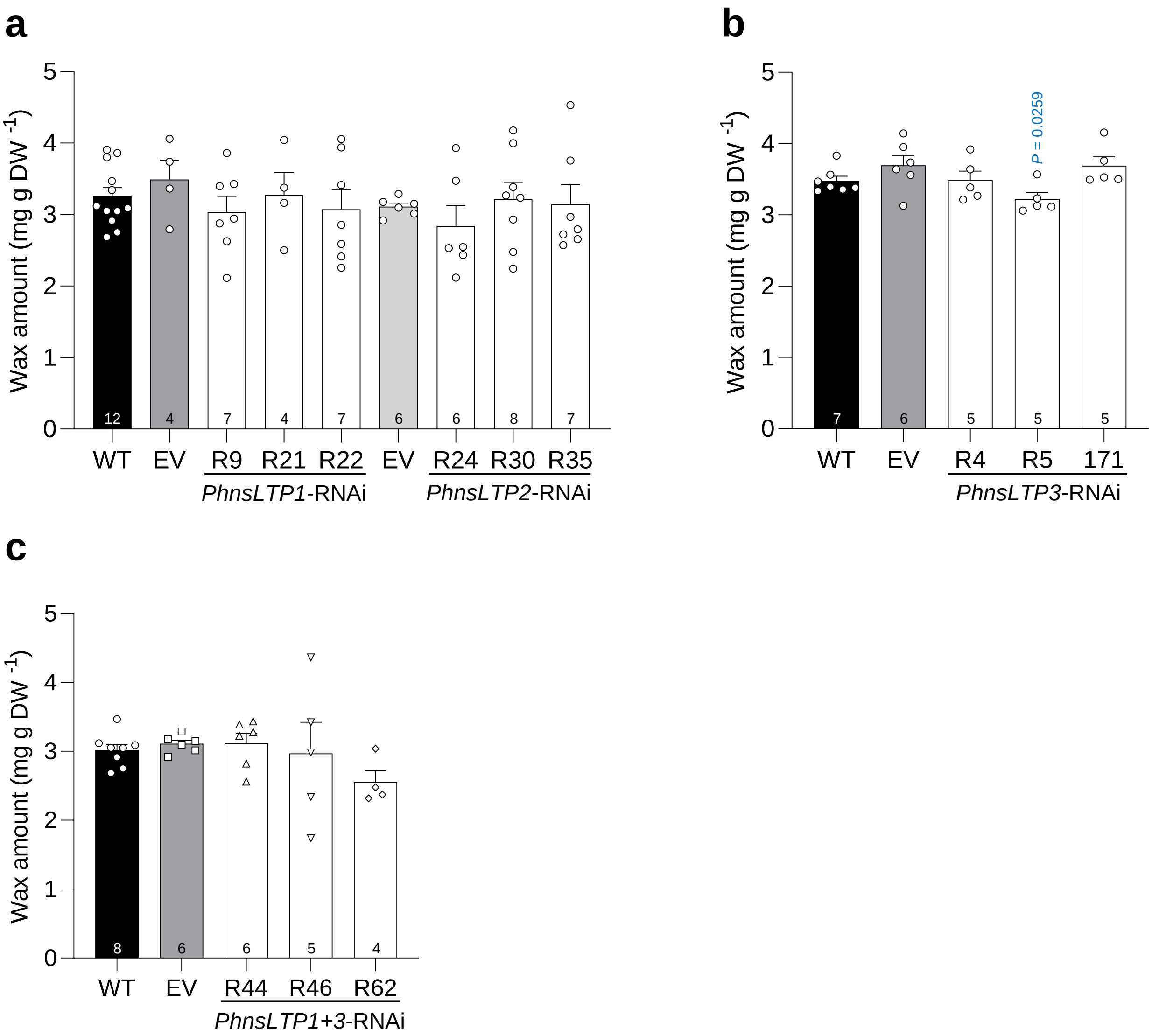


**Supplementary Figure 11 | Total wax amount in petunia petals.** Amount of total wax in 2-day-old flowers of wild type (WT), empty vector control (EV), three *PhnsLTP1*-RNAi lines (R9, R21, and R22), and three *PhnsLTP2*-RNAi lines (R24, R30, and R35) (**a**), three *PhnsLTP3*-RNAi lines (R4, R5, and 171) (**b**) and three *PhnsLTP1*+*PhnsLTP3*-RNAi lines (R44, R46, and R62) (**c**). The total wax amount was normalized by the dry weight (DW). Data are means ± S.E. Numbers at the bottoms of the columns represent the numbers of biological replicates. *P* values were determined by one-way ANOVA with the Dunnett's multiple comparisons test relative to the WT and EV (blue) controls.

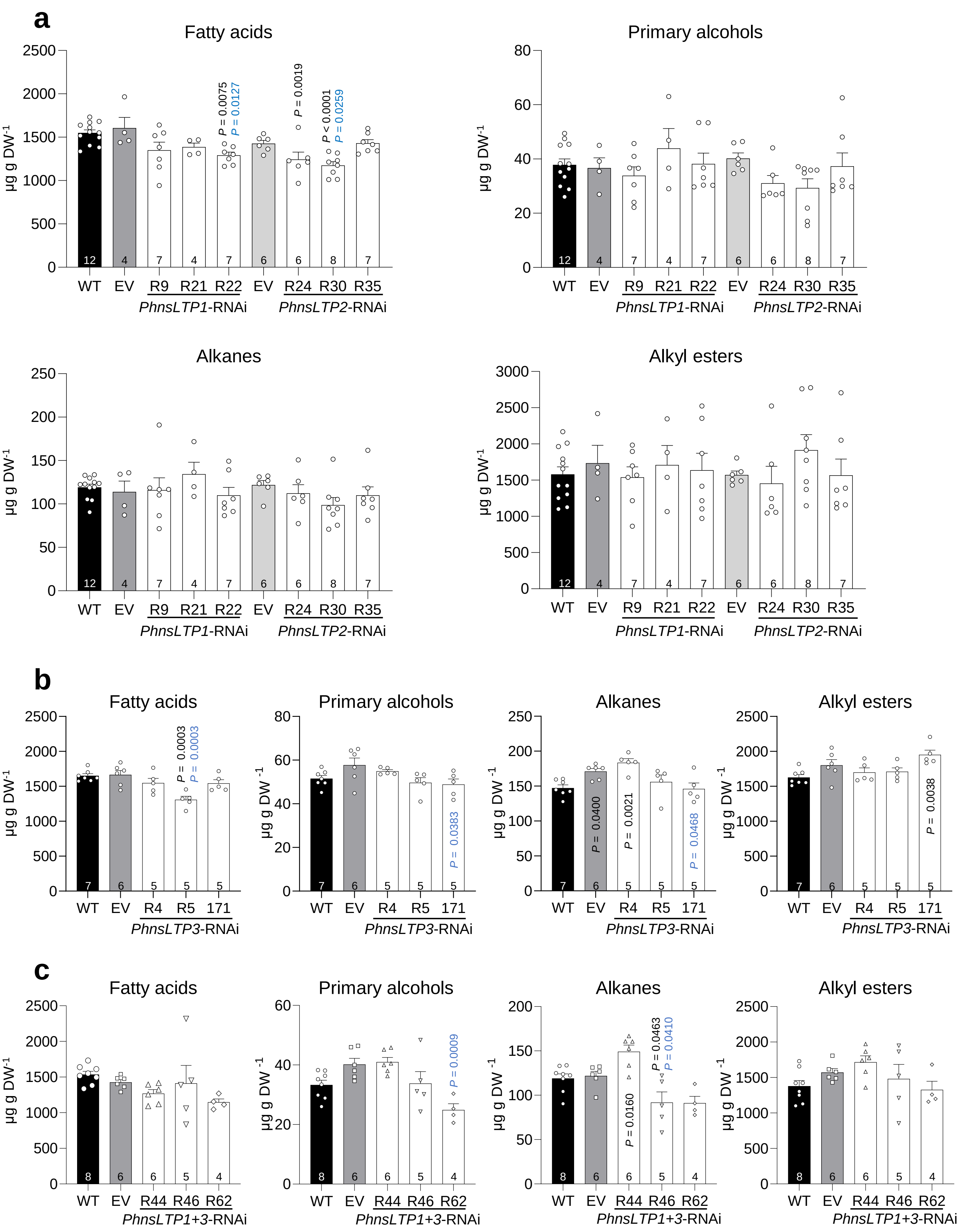
a
Fatty acids
Primary alcohols
Alkanes
Alkyl esters
μg g DW-1
WT EV R9 R21 R22 EV R24 R30 R35
PhnsLTP1-RNAi
PhnsLTP2-RNAi
12 4 7 4 7 6 6 8 7
P = 0.0075
P = 0.0127
P = 0.0019
P < 0.0001
P = 0.0259
b
Fatty acids
Primary alcohols
Alkanes
Alkyl esters
μg g DW-1
WT EV R4 R5 171
PhnsLTP3-RNAi
7 6 5 5 5
P = 0.0003
P = 0.0003
P = 0.0383
P = 0.0400
P = 0.0021
P = 0.0468
P = 0.0038
c
Fatty acids
Primary alcohols
Alkanes
Alkyl esters
μg g DW-1
WT EV R44 R46 R62
PhnsLTP1+3-RNAi
8 6 6 5 4
P = 0.0009
P = 0.0160
P = 0.0463
P = 0.0410

**Supplementary Figure 12 | Wax amount of major constituent classes in petunia petals.** Amounts of fatty acids, primary alcohols, alkanes, and alkyl esters are shown in 2-day-old flowers of wild type (WT), empty vector control (EV), three *PhnsLTP1*-RNAi lines (R9, R21, and R22), and three *PhnsLTP2*-RNAi lines (R24, R30, and R35) (**a**), three *PhnsLTP3*-RNAi lines (R4, R5, and 171) (**b**) and three *PhnsLTP1+PhnsLTP3*-RNAi lines (R44, R46, and R62) (**c**). Amounts of fatty acids, primary alcohols, alkanes, and alkyl esters were normalized by the dry weight (DW). Data are means ± S.E. Numbers at the bottoms of the columns represent the numbers of biological replicates. *P* values were determined by one-way ANOVA with the Dunnett's multiple comparisons test relative to the WT (black) and EV (blue) controls.

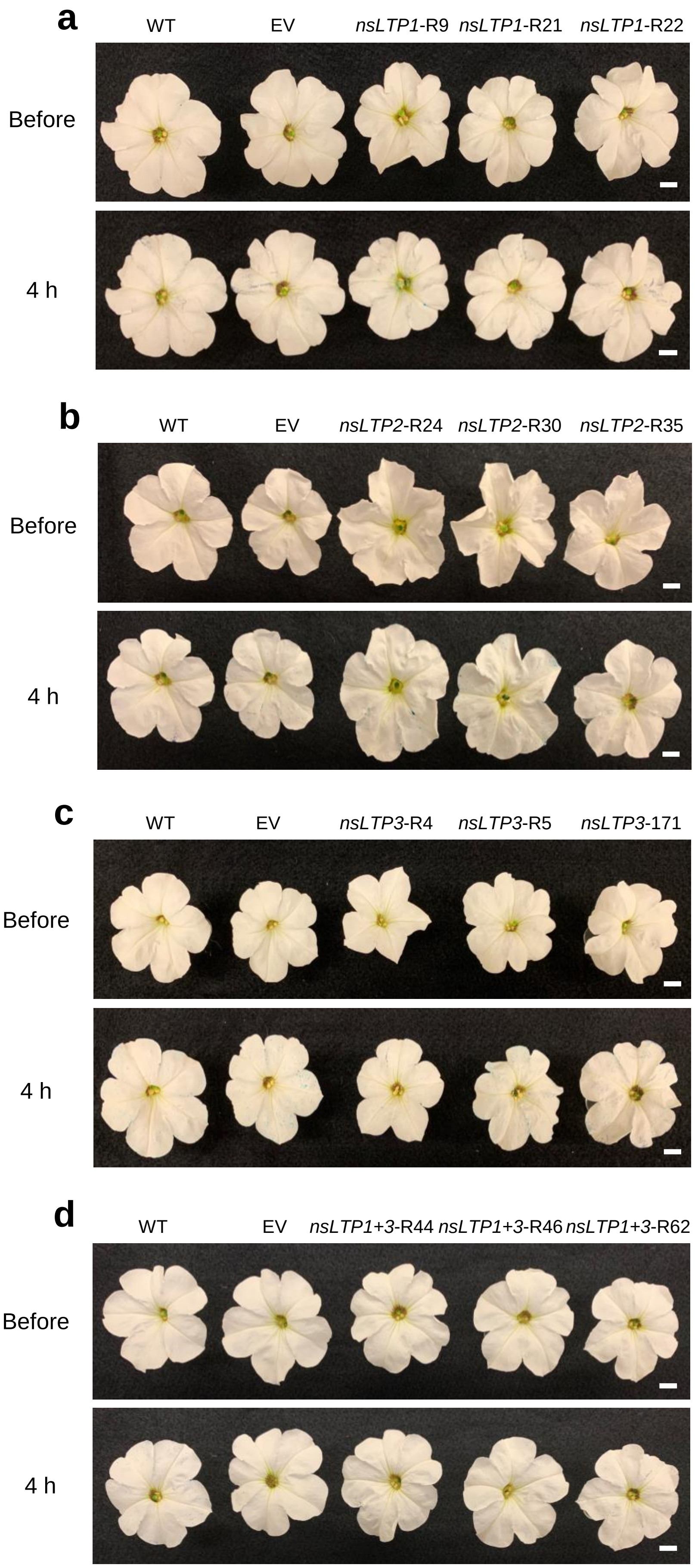

a
WT
EV
nsLTP1-R9
nsLTP1-R21
nsLTP1-R22
Before
4 h
b
WT
EV
nsLTP2-R24
nsLTP2-R30
nsLTP2-R35
Before
4 h
c
WT
EV
nsLTP3-R4
nsLTP3-R5
nsLTP3-171
Before
4 h
d
WT
EV
nsLTP1+3-R44
nsLTP1+3-R46
nsLTP1+3-R62
Before
4 h

**Supplementary Figure 13 | Effect of *PhnsLTP1, PhnsLTP2, PhnsLTP3* and *PhnsLTP1+ PhnsLTP3* downregulation on cuticle permeability in petunia petals.** Representative 2-day-old petunia flowers from wild type (WT), empty vector control (EV), three independent *PhnsLTP1*-RNAi lines (R9, R21, and R22) (**a**), three independent *PhnsLTP2*-RNAi lines (R24, R30, and R35) (**b**), three independent *PhnsLTP3*-RNAi lines (R4, R5, and 171) (**c**), and three independent *PhnsLTP1+PhnsLTP3*-RNAi lines (R44, R46, and R62) (**d**), before (upper panel) and 4 h after staining with toluidine blue (lower panel). Scale bar, 1 cm.

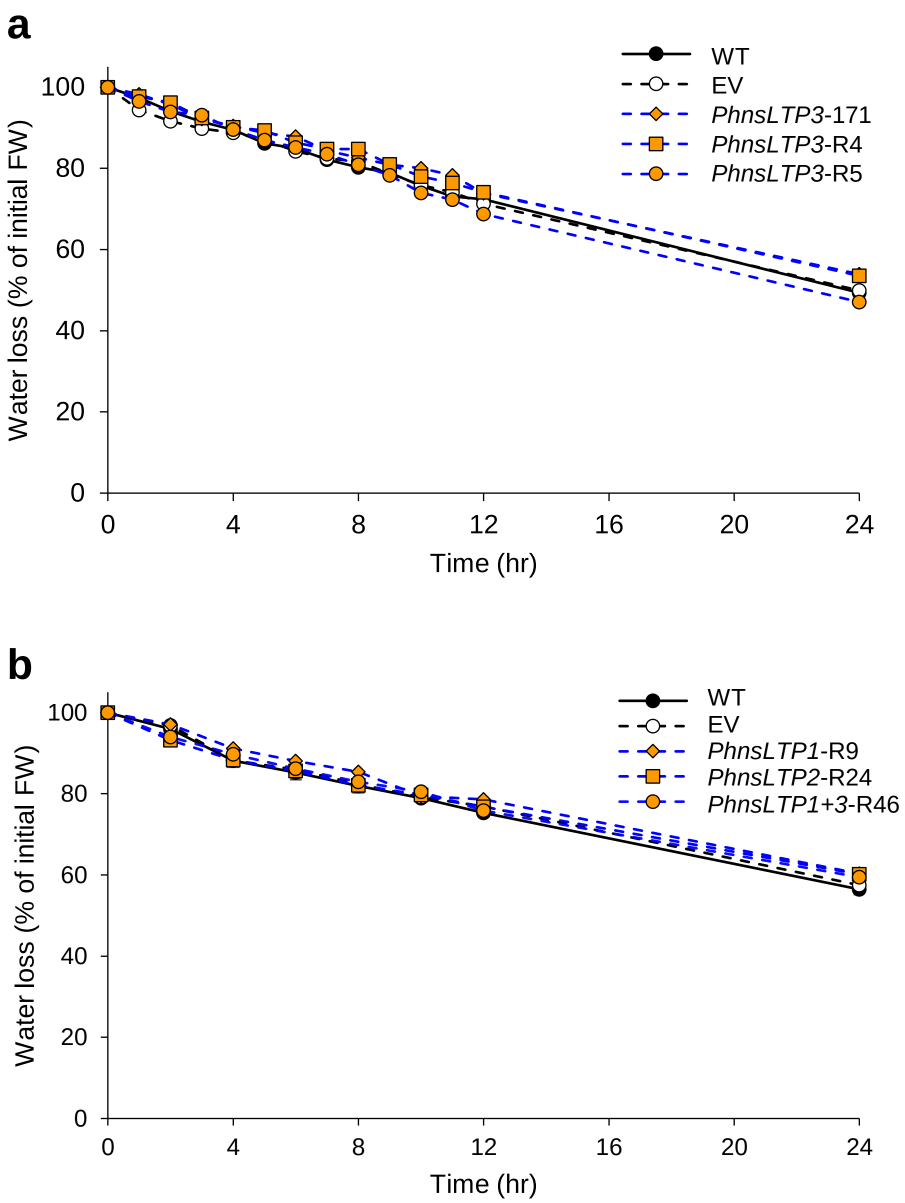


**Supplementary Figure 14 | Effect of *PhnsLTP3, PhnsLTP1, PhnsLTP2 and PhnsLTP1+PhnsLTP3* downregulation on water loss in petunia corollas. a,** Water loss in detached 2-day-old flowers from wild-type (WT), empty vector control (EV), and three independent *PhnsLTP3*-RNAi lines (R4, R5 and 171) were measured over 24 h and expressed relative to initial weight. **b**, Water loss in detached 2-day-old flowers from WT, EV, and one representative of *PhnsLTP1*-RNAi line (R9), *PhnsLTP2*-RNAi line (R24), and *PhnsLTP1+PhnsLTP3*-RNAi line (R46) were measured as in (**a**). Data are means ± S.E. (n = 3 and 5 biological replicates of three flowers each in **a** and **b**, respectively). FW, fresh weight.

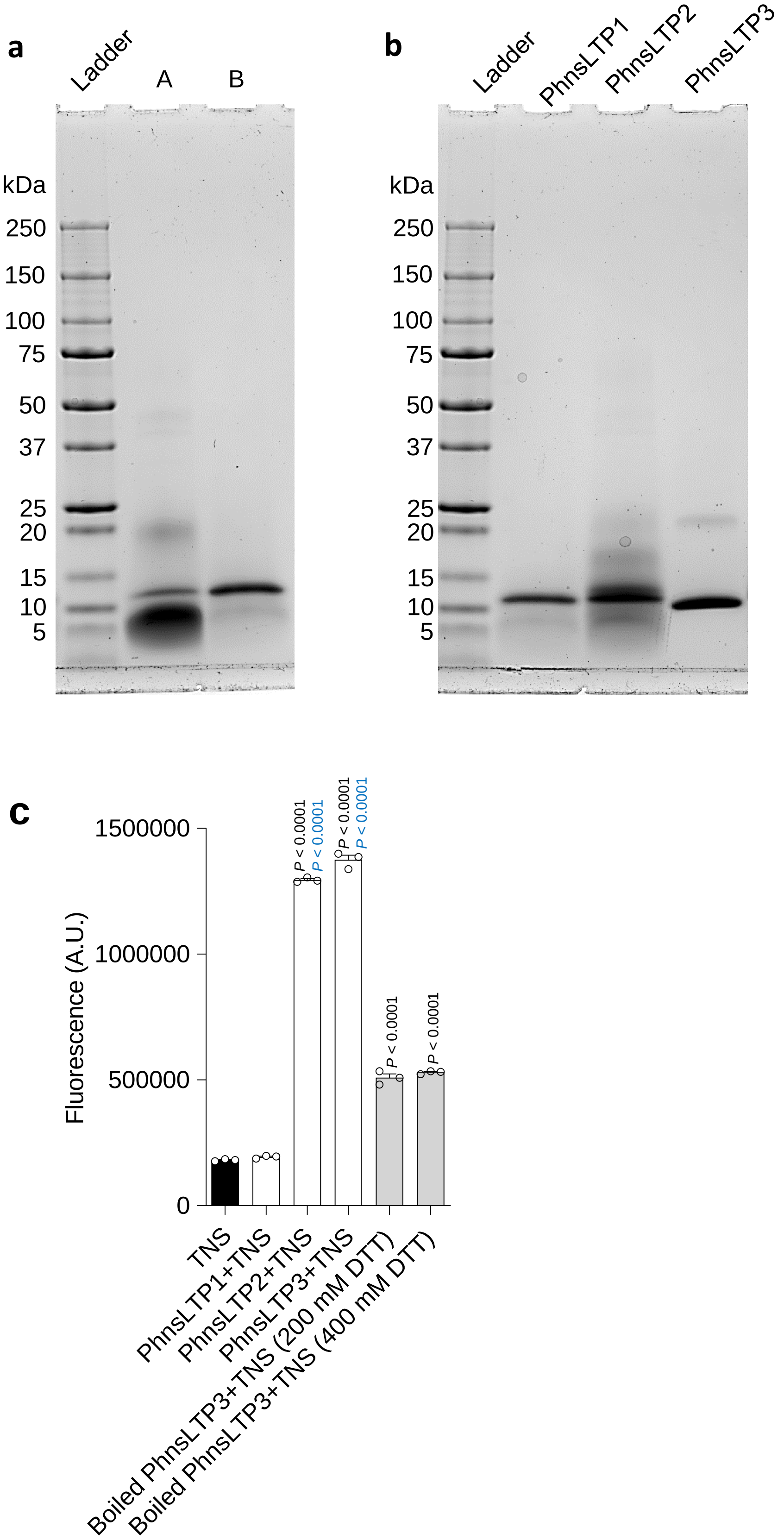

a
Ladder
A
B
kDa
250
150
100
75
50
37
25
20
15
10
5
b
Ladder
PhnsLTP1
PhnsLTP2
PhnsLTP3
kDa
250
150
100
75
50
37
25
20
15
10
5
c
Fluorescence (A.U.)
1500000
1000000
500000
0
P < 0.0001
P < 0.0001
P < 0.0001
P < 0.0001
P < 0.0001
P < 0.0001
TNS
PhnsLTP1+TNS
PhnsLTP2+TNS
PhnsLTP3+TNS
Boiled PhnsLTP3+TNS (200 mM DTT)
Boiled PhnsLTP3+TNS (400 mM DTT)

**Supplementary Figure 15 | Protein purification and characterization of PhnsLTP1, PhnsLTP2, and PhnsLTP3 activity with TNS. a,** SDS-PAGE analysis of PhnsLTP1 after affinity chromatography on Ni-NTA agarose (lane A) and subsequent cation exchange chromatography on HiTrap$^{TM}$ SP HP-Cytiva (lane B). 5.3 µg protein were loaded per each lane. **b,** SDS-PAGE analysis of 8.75 µg of purified PhnsLTP1, PhnsLTP2, and PhnsLTP3. The experiments in **a** and **b** were repeated at least four times with similar results. **c,** Activity of PhnsLTP1, PhnsLTP2 and PhnsLTP3 with TNS. Only PhnsLTP2 and PhnsLTP3 show TNS binding. TNS only, as well as TNS with boiled PhnsLTP3 protein and 200 mM or 400 mM DTT were used as negative controls. All data are means ± S.E. (n = 3 biological replicates). *P* values were determined by one-way ANOVA with the Tukey's multiple comparisons test relative to TNS background (black) and boiled PhnsLTP3 in presence 200 mM DTT (blue).

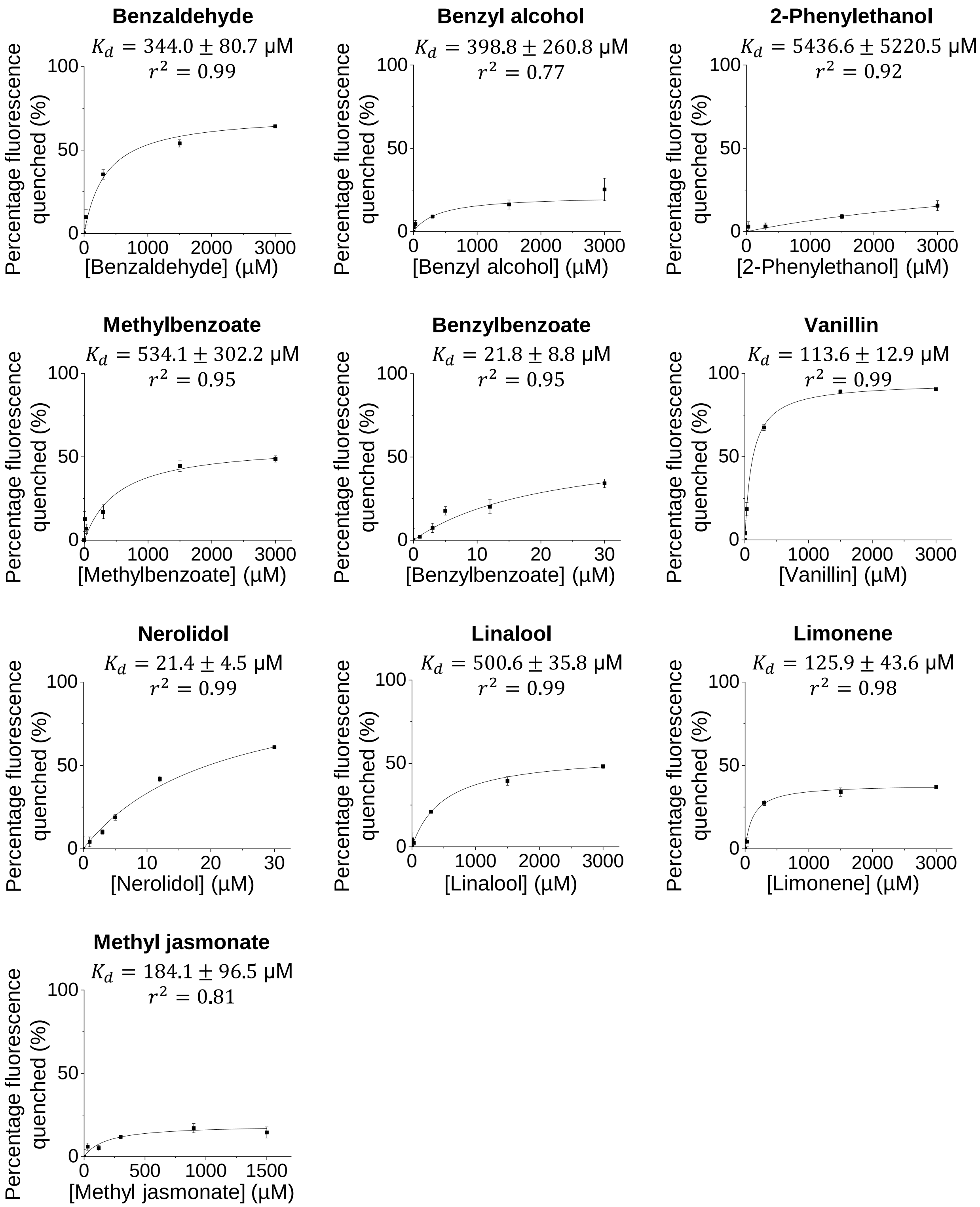
Benzaldehyde
$K_d = 344.0 \pm 80.7$ μM
$r^2 = 0.99$
Percentage fluorescence quenched (%)
[Benzaldehyde] (μM)
Benzyl alcohol
$K_d = 398.8 \pm 260.8$ μM
$r^2 = 0.77$
Percentage fluorescence quenched (%)
[Benzyl alcohol] (μM)
2-Phenylethanol
$K_d = 5436.6 \pm 5220.5$ μM
$r^2 = 0.92$
Percentage fluorescence quenched (%)
[2-Phenylethanol] (μM)
Methylbenzoate
$K_d = 534.1 \pm 302.2$ μM
$r^2 = 0.95$
Percentage fluorescence quenched (%)
[Methylbenzoate] (μM)
Benzylbenzoate
$K_d = 21.8 \pm 8.8$ μM
$r^2 = 0.95$
Percentage fluorescence quenched (%)
[Benzylbenzoate] (μM)
Vanillin
$K_d = 113.6 \pm 12.9$ μM
$r^2 = 0.99$
Percentage fluorescence quenched (%)
[Vanillin] (μM)
Nerolidol
$K_d = 21.4 \pm 4.5$ μM
$r^2 = 0.99$
Percentage fluorescence quenched (%)
[Nerolidol] (μM)
Linalool
$K_d = 500.6 \pm 35.8$ μM
$r^2 = 0.99$
Percentage fluorescence quenched (%)
[Linalool] (μM)
Limonene
$K_d = 125.9 \pm 43.6$ μM
$r^2 = 0.98$
Percentage fluorescence quenched (%)
[Limonene] (μM)
Methyl jasmonate
$K_d = 184.1 \pm 96.5$ μM
$r^2 = 0.81$
Percentage fluorescence quenched (%)
[Methyl jasmonate] (μM)

**Supplementary Figure 16 | Binding kinetics of recombinant PhnsLTP2 with phenylpropanoids/benzenoids by petunia flowers, terpenoids and methyl jasmonate.** TNS displacement assays were performed at fixed concentrations of PhnsLTP2 (1 µM) and TNS (3 µM) and the fluorescence was measured before and after the tested compounds were added at various concentrations. The results are expressed as percentage of fluorescence quenched with respect to the concentration of added compound. The binding curves were obtained by non-linear fitting to the Michaelis-Menten equation (solid line in the graph) and the dissociation constants $K_d$ [µM] were obtained. Phenylacetaldehyde, eugenol, and isoeugenol had high fluorescence background with TNS, which did not allow obtaining the corresponding binding curves. All data are means ± S.E. (n = 3 biological replicates).

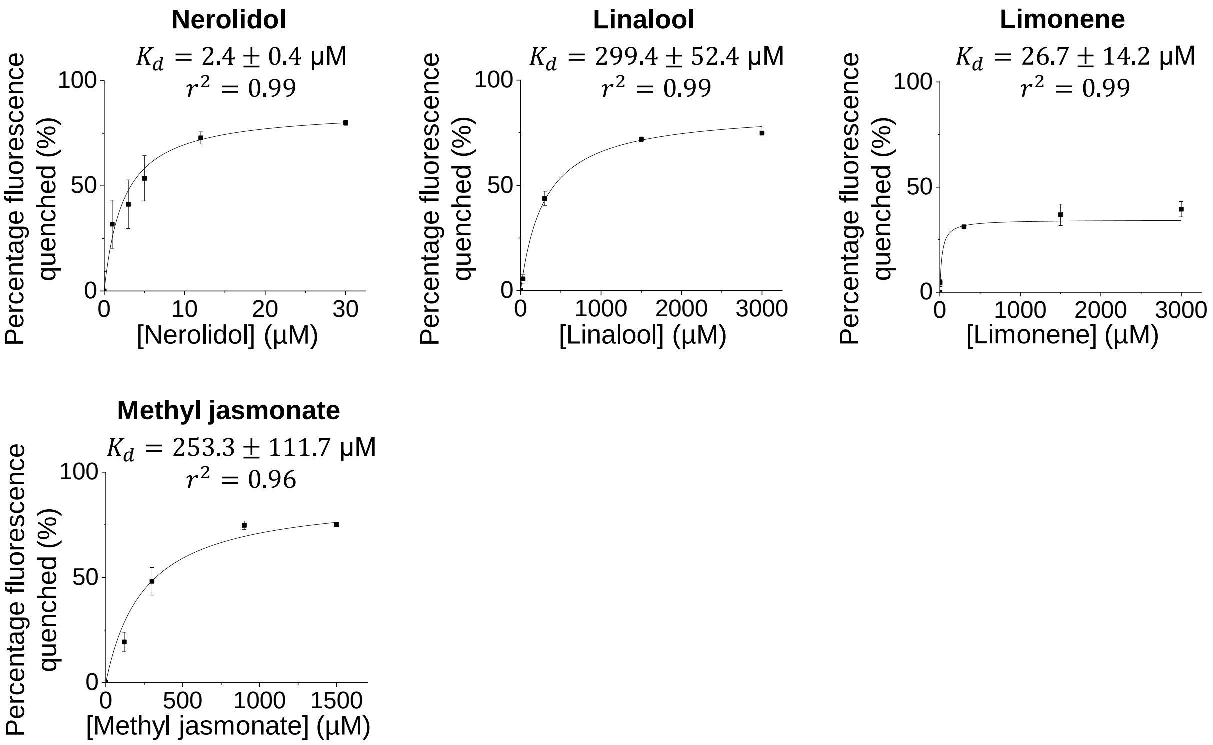


**Supplementary Figure 17 | Binding kinetics of recombinant PhnsLTP3 with monoterpenes and methyl jasmonate.** TNS displacement assays were performed at fixed concentrations of PhnsLTP3 (1 μM) and TNS (3 μM) and the fluorescence was measured before and after the tested compounds were added at various concentrations. The results are expressed as percentage of fluorescence quenched with respect to the concentration of added compound. The binding curves were obtained by non-linear fitting to the Michaelis-Menten equation (solid line in the graph) and the dissociation constants $K_d$ [μM] were obtained. All data are means ± S.E. (n = 3 biological replicates).

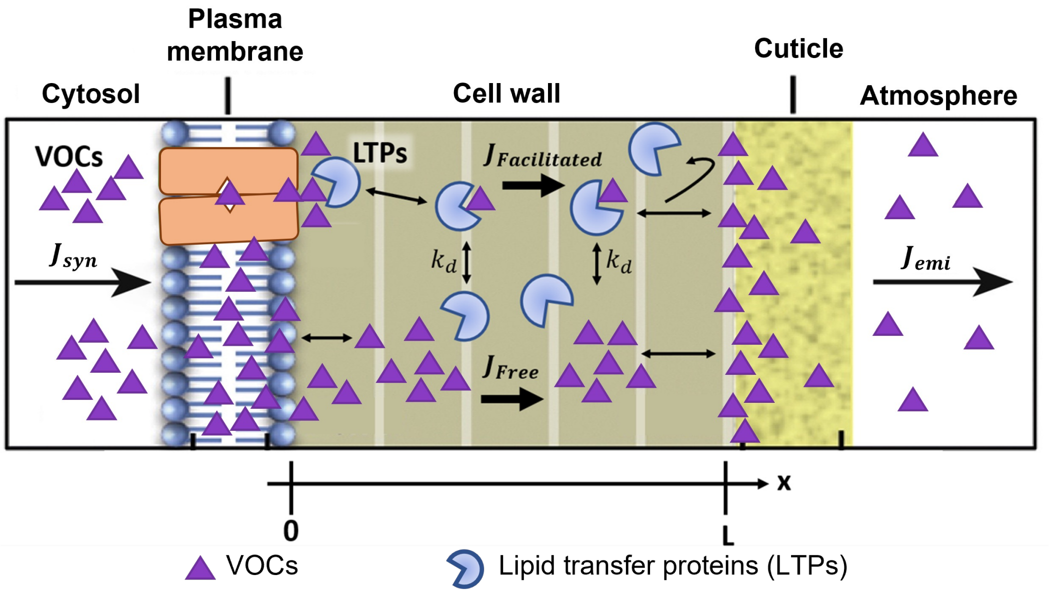


**Supplementary Figure 18 | Proposed role of nsLTPs in VOC emission.** Facilitated diffusion was used to describe the nsLTP-mediated transport of VOCs across the cell wall. nsLTPs are hypothesized to spontaneously uptake the VOCs from the left boundary of the cell wall layer (at x=0). Both the free VOCs and the nsLTP-VOC complexes then move in the positive x direction down the concentration gradient to the right boundary, where nsLTPs unload the cargo to the cuticle. The total flux across the cell wall is the sum of the free VOC flux and the facilitated flux of VOCs in the complex with LTPs. $J_{syn}$, VOC biosynthetic flux; $J_{Facilitated}$, facilitated VOC flux by nsLTPs; $J_{Free}$, free VOC diffusion flux; $J_{emi}$, VOC emission flux; $k_d$, equilibrium dissociation constant; $S_0$, concentration of total VOCs at the apical facing site of the plasma membrane; $S_L$, concentration of total VOCs at the inner site of the cuticle.

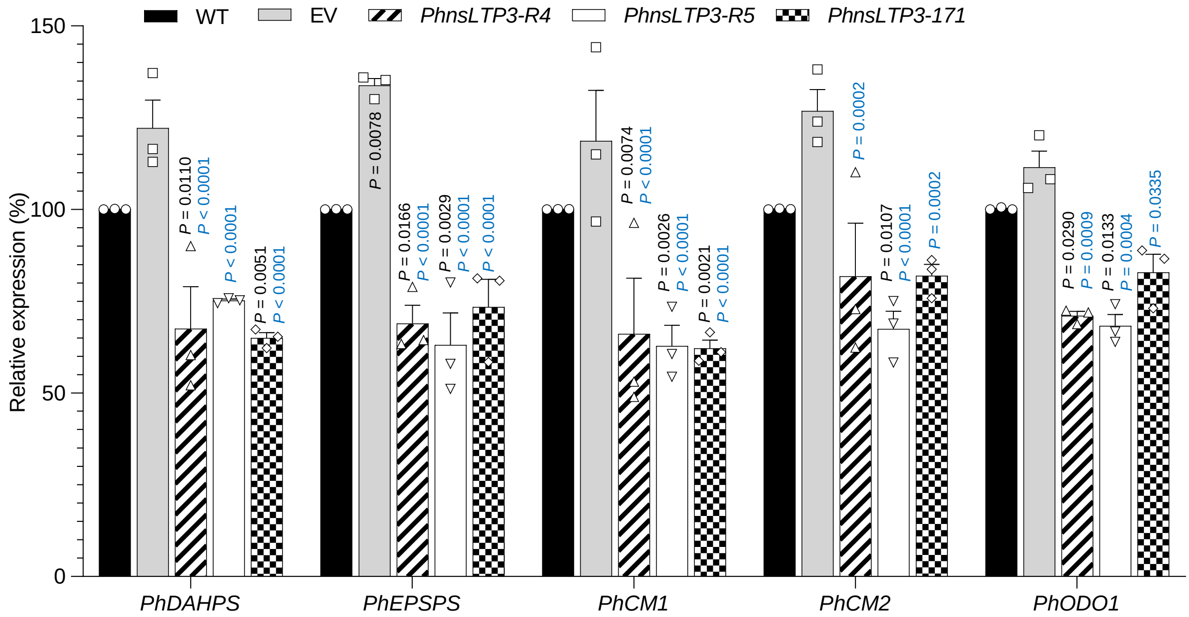


**Supplementary Figure 19 | Effect of *PhnsLTP3* downregulation on expression of *PhDAHPS*, *PhEPSPS*, *PhCM1*, *PhCM2* and *PhODO1* in petunia flowers.**

Transcript levels were determined by qRT-PCR with gene specific primers in 2-day-old flowers from wild type (WT), empty vector control (EV) and three independent *PhnsLTP3*-RNAi lines (R4, R5 and 171) harvested at 15:00 h and presented relative to the corresponding levels in WT, set as 100%. Data are means ± S.E. (n = 3 biological replicates). *P* values were determined by two-way ANOVA with the Tukey's multiple comparisons test relative to the corresponding WT (black) and EV (blue) controls. Displayed gene identifiers encode the following proteins: *PhDAHPS*, 3-deoxy-D-arabino-heptulosonate 7-phosphate synthase; *PhEPSPS*, 5-enolpyruvylshikimate 3-phosphate synthase; *PhCM1* and *PhCM2*, chorismate synthase 1 and 2, respectively; and *PhODO1*, ODORANT1 transcription factor.

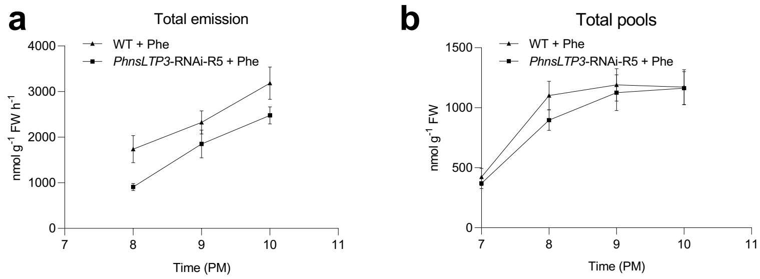


**Supplementary Figure 20 | Total VOC emission and their total pools in control and *PhnsLTP3*-RNAi petunia flowers fed with 150 mM Phe.** Total VOC emission (**a**) and total pools (**b**) in wild-type (WT) and *PhnsLTP3*-RNAi-R5 line fed with 150 mM Phe from 19:00 to 22:00 h. Data are means ± S.E. (n = 3 biological replicates). In (**a**) the overall emission in transgenics is significantly lower than in WT as determined by two-way ANOVA with $P = 0.0086$.

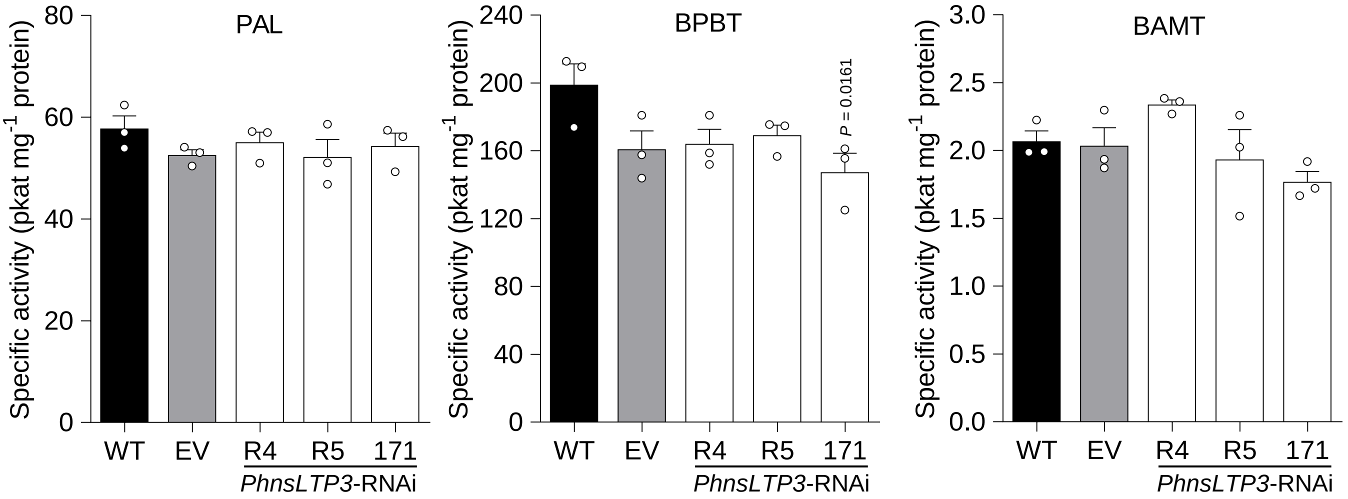


**Supplementary Figure 21 | Effect of *PhnsLTP3* downregulation on enzyme activities of phenylalanine ammonia lyase (PAL), benzoyl-CoA:benzylalcohol/2-phenylethanol benzoyltransferase (BPBT) and benzoic acid carboxyl methyltransferase (BAMT).** Enzyme activities of PAL, BPBT and BAMT were measured in 2-day-old flowers of wild type (WT), empty vector control (EV), and three *PhnsLTP3*-RNAi lines (R4, R5, and 171) harvested at 20:00 h. All data are means ± S.E. (n = 3 biological replicates). *P* values were determined by one-way ANOVA with the Dunnett's multiple comparisons test relative to the WT and EV controls.

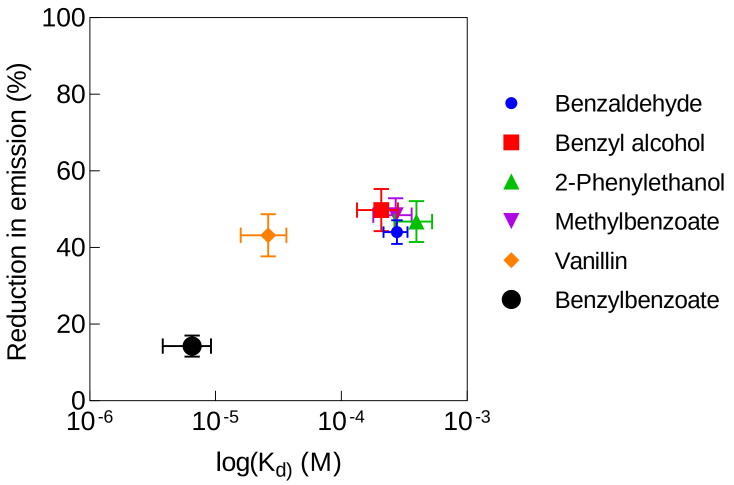


**Supplementary Figure 22 | Relationship between the $K_d$ values for individual VOCs and a reduction in the emission of the corresponding compound upon *PhnsLTP3* downregulation.** Reductions in emission (%) of individual VOCs are shown for *PhnsLTP3*-RNAi line R5 (see Supplementary Figure 9); log ($K_d$) values were calculated based on data in Figure 5.

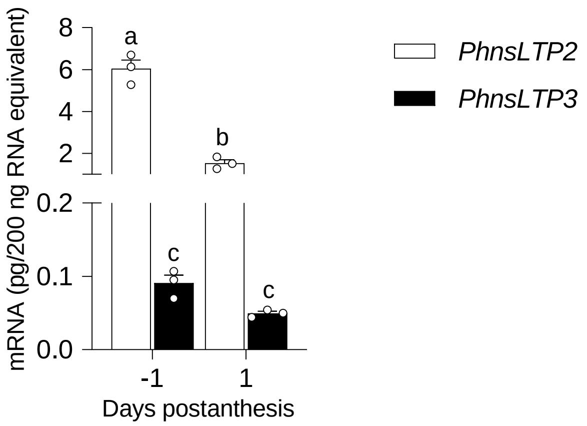


**Supplementary Figure 23 | Expression analysis of *PhnsLTP2* and *PhnsLTP3* in petunia stigma.** Expression of *PhnsLTP2* and *PhnsLTP3* in petunia stigma at 15:00 h on days -1 and 1 postanthesis. Absolute mRNA levels were determined by qRT-PCR with gene-specific primers and are shown as pg/200 ng total RNA. Data are means ± S.E. (n = 3 biological replicates). Different letters indicate statistically significant differences ($P < 0.05$) determined by one-way ANOVA with the Tukey's multiple comparisons test.

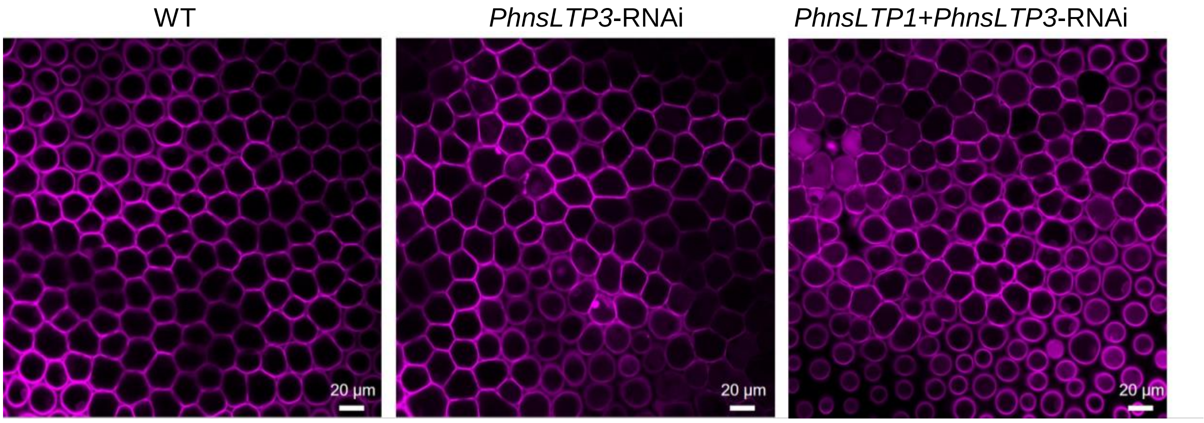


**Supplementary Figure 24 | Effect of *PhnsLTP3* and *PhnsLTP1+PhnsLTP3* downregulation on cell membrane integrity in petunia flowers.** Representative confocal microscopy images of conical epidermal cells of 2-day-old wild type (WT), *PhnsLTP3*-171 RNAi and *PhnsLTP1+PhnsLTP3*-R44 RNAi flowers stained with propidium iodide. Scale bars, 20 µm. Similar results were obtained in three independent samples.

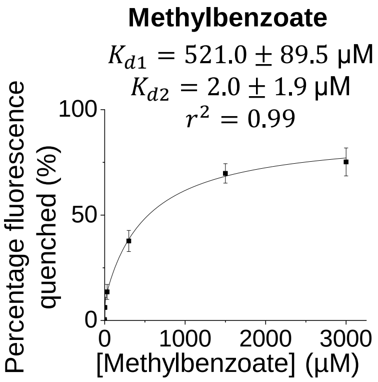


**Supplementary Figure 25 | Analysis of number of binding sites of PhnsLTP3.** TNS displacement assay was performed at fixed concentrations of PhnsLTP3 (1 μM) and TNS (3 μM) and the fluorescence was measured before and after methyl benzoate was added at various concentrations. The results are expressed as percentage of fluorescence quenched with respect to the concentration of added compound. The binding curve was obtained by non-linear fitting to the two-binding-site model (solid line in the graph) and the dissociation constants $K_d$ [μM] were obtained. All data are means ± S.E. (n = 3 biological replicates).

**Supplementary Table 1. Features of PhLTP1, PhLTP2 and PhLTP3 proteins.**

| Protein name | Size of signal peptide (aa*) | Size of mature protein (aa*) | Predicted mass for mature protein (kDa) |
|---|---|---|---|
| PhLTP1 | 25 | 91 | 9.35 |
| PhLTP2 | 26 | 90 | 9.09 |
| PhLTP3 | 22 | 93 | 9.65 |

* aa, amino acids.

**Supplementary Table 2. Protein sequence identity and similarity between mature LTPs from *Petunia hybrida* (Ph), *Nicotiana tabacum* (Nt) and *Artemisia annua* (Aa).**

| | PhLTP1 | PhLTP2 | PhLTP3 | NtLTP1 | AaLTP3 |
|---|---|---|---|---|---|
| PhLTP1 | 100 (100) | 58 (69) | 40 (60) | 57 (68) | 46 (60) |
| PhLTP2 | | 100 (100) | 41 (60) | 69 (80) | 59 (70) |
| PhLTP3 | | | 100 (100) | 46 (63) | 46 (64) |
| NtLTP1 | | | | 100 (100) | 58 (70) |
| AaLTP3 | | | | | 100 (100) |

Shown are percentages of amino acid identity and similarity (in brackets).

**Supplementary Table 3. VOC emission factors (VEFs) in wild-type and *PhnsLTP3*-RNAi petunia flowers from 17:00 to 22:00 h.**

| | Time (h) | Wild type | *PhnsLTP3*-RNAi | *P* value* |
|---|---|---|---|---|
| Figure 7e | 17:00 | 0.2236 ± 0.0143<br>n = 9 | 0.2155 ± 0.0178<br>n = 5 | 0.9848 |
| Figure 7f | 18:00 | 0.3491 ± 0.0230<br>n = 9 | 0.3239 ± 0.0107<br>n = 5 | 0.7003 |
| | 19:00 | 0.4892 ± 0.0119<br>n = 9 | 0.3797 ± 0.0134<br>n = 5 | 0.0004 |
| | 20:00 | 0.6525 ± 0.0048<br>n = 8 | 0.5916 ± 0.0199<br>n = 4 | 0.0057 |
| | 21:00 | 0.7011 ± 0.0126<br>n = 7 | 0.6239 ± 0.0220<br>n = 5 | 0.0003 |
| | 22:00 | 0.7183 ± 0.0066<br>n = 8 | 0.6388 ± 0.0093<br>n = 5 | 0.0001 |

Data are means ± S.E. (n = biological replicates). * *P* values were determined by two-way ANOVA with the Sidak's multiple comparisons test relative to the corresponding wild type at each time point.

### Supplementary Table 4. Primers used in this study.

| Primer name | Direction | Sequence | Purpose |
|---|---|---|---|
| PhnsLTP1_RNAi_F | Forward | 5'-AAAAAGCAGGCTTCAGATCTGAATTCATGGTT-3' | Cloning |
| PhnsLTP1_RNAi_R | Reverse | 5'-AGAAAGCTGGGTCACTAGTGGATCCATGGTT-3' | Cloning |
| PhnsLTP1:RFP | Forward | 5'-AAAAAGCAGGCTTCATGAAAATGGTTGGAAAGATT-3' | Cloning |
| PhnsLTP1:RFP | Reverse | 5'-AGAAAGCTGGGTCGTTGACCCTGGAGCAGTCAGT-3' | Cloning |
| PhnsLTP2:RFP | Forward | 5'-AAAAAGCAGGCTTCATGGCAATGGCTAGCAAAATT-3' | Cloning |
| PhnsLTP2:RFP | Reverse | 5'-AGAAAGCTGGGTCCTGGACCTTGGAGCAGTCAGT-3' | Cloning |
| PhnsLTP3:RFP | Forward | 5'-AAAAAGCAGGCTTCATGCTAAACAAGCAACTACTT-3' | Cloning |
| PhnsLTP3:RFP | Reverse | 5'-AGAAAGCTGGGTCCTTGATCTTAGAGCAATCAAC-3' | Cloning |
| AtPIP1:GFP | Forward | 5'-GGGGACAAGTTTGTACAAAAAAGCAGGCTTAATGGCAAAGGATGTG-3' | Cloning |
| AtPIP1:GFP | Reverse | 5'-GGGGACCACTTTGTACAAGAAAGCTGGGTTGACGTTGGCAGC-3' | Cloning |
| PhnsLTP1_piczalp_NS_F | Forward | 5'-AGAGAGGCTGAAGCTGCAATGACCTGCGGCC-3' | Cloning |
| PhnsLTP1_piczalp_NS_R | Reverse | 5'-ATGATGATGATGATGGTTGACCCTGGAGCAGTCAG-3' | Cloning |
| PhnsLTP2_piczalp_NS_F | Forward | 5'-AGAGAGGCTGAAGCTTTAACTTGTGGCCAAGTTACATCCAAT-3' | Cloning |
| PhnsLTP2_piczalp_NS_R | Reverse | 5'-ATGATGATGATGATGCTGGACCTTGGAGCAGTCAG-3' | Cloning |
| PhnsLTP3_piczalp_NS_F | Forward | 5'-AGAGAGGCTGAAGCTGCAACCATCACGTGCAGC-3' | Cloning |
| PhnsLTP3_piczalp_NS_R | Reverse | 5'-ATGATGATGATGATGCTTGATCTTAGAGCAATCAACGTCTGG-3' | Cloning |
| PhnsLTP1_qPCR_F | Forward | 5'-GAATCATGGCCCTCTTGGAA-3' | qRT-PCR |
| PhnsLTP1_qPCR_R | Reverse | 5'-TCCTACGGTCCTGTGGTGTCT-3' | qRT-PCR |
| PhnsLTP2_qPCR_F | Forward | 5'-TGGATCTTTAGGAGGCTGTTGT-3' | qRT-PCR |
| PhnsLTP2_qPCR_R | Reverse | 5'-TGACACCACAAGTACTGGGAAG-3' | qRT-PCR |
| PhnsLTP3_qPCR_F | Forward | 5'-GCTTCTTCTTTGCATCGTGGC-3' | qRT-PCR |
| PhnsLTP3_qPCR_R | Reverse | 5'-ATACTAGCAGCACGGCTGATTT-3' | qRT-PCR |
| PhDAHPS_qPCR_F | Forward | 5'-CAAAGCTCCGTGTGGTCTTAAA-3' | qRT-PCR |
| PhDAHPS_qPCR_R | Reverse | 5'-TCCTGGGTGGCTTCCTTCTT-3' | qRT-PCR |
| PhEPSPS_qPCR_F | Forward | 5'-CACCCCACCGGAGAAACTAA-3' | qRT-PCR |
| PhEPSPS_qPCR_R | Reverse | 5'-TGACGGGAACATCTGCACAA-3' | qRT-PCR |
| PhCM1_qPCR_F | Forward | 5'-CCTGCTGTTGAAGAGGCTATCA-3' | qRT-PCR |
| PhCM1_qPCR_R | Reverse | 5'-CAGGGTCACCTCCATTTTCTG-3' | qRT-PCR |
| PhCM2_qPCR_F | Forward | 5'-TGCAACTACTGCTGCCTGTGAT-3' | qRT-PCR |
| PhCM2_qPCR_R | Reverse | 5'-TGCAACTACTGCTGCCTGTGAT-3' | qRT-PCR |
| PhODO1_qPCR_F | Forward | 5'-CCACTCAAGAAAGAAGCAAATCTAAGT-3' | qRT-PCR |
| PhODO1_qPCR_R | Reverse | 5'-TGCCGCTGTGACATTTGTACTC-3' | qRT-PCR |
| PhEF1-α | Forward | 5'-CCTGGTCAAATTGGAAACGG-3' | qRT-PCR |
| PhEF1-α | Reverse | 5'-CAGATCGCCTGTCAATCTTGG-3' | qRT-PCR |
| α-Factor | Forward | 5'-TACTATTGCCAGCATTGCTGC-3' | PCR |
| 3'AOX1 | Reverse | 5'-GCAAATGGCATTCTGACATCC-3' | PCR |

**Supplementary Table 5. Absolute amount of total VOCs in cellular and cuticular fractions shown in Fig. 4.**

| | | Cellular VOC amount (nmol g $FW^{-1}$) | Cuticular VOC amount (nmol g $FW^{-1}$) |
|---|---|---|---|
| Figure 4a | Wild type | 1069.7 ± 60.2 | 730.4 ± 51.4 |
| | Empty vector | 1197.8 ± 82.2 | 716.5 ± 52.6 |
| | *PhnsLTP1*-RNAi-R9 | 1521.3 ± 127.2 | 814.1 ± 108.3 |
| | *PhnsLTP1*-RNAi-R21 | 1142.1 ± 80.1 | 700.9 ± 65 |
| | *PhnsLTP1*-RNAi-R22 | 1126.4 ± 99.1 | 619.6 ± 70.7 |
| | Empty vector | 1180.8 ± 51.9 | 671.4 ± 39.2 |
| | *PhnsLTP2*-RNAi-R24 | 1209.4 ± 76.9 | 682.5 ± 58.9 |
| | *PhnsLTP2*-RNAi-R30 | 1359.9 ± 69.6 | 597.9 ± 41.9 |
| | *PhnsLTP2*-RNAi-R35 | 1076.3 ± 55.4 | 578.2 ± 36.2 |
| Figure 4b | Wild type | 717.8 ± 29.8 | 474.4 ± 21.9 |
| | Empty vector | 769.6 ± 35.4 | 454.7 ± 24.4 |
| | *PhnsLTP3*-RNAi-R4 | 813.8 ± 44.7 | 369.7 ± 27.8 |
| | *PhnsLTP3*-RNAi-R5 | 789.9 ± 49.8 | 302.3 ± 23.4 |
| | *PhnsLTP3*-RNAi-171 | 781.7 ± 41.6 | 304.6 ± 24.6 |

All data are means ± S.E. FW, fresh weight.

**Supplementary Table 6. The values of parameters used in the model.**

| Parameter | Symbol [unit] | Value |
|---|---|---|
| Diffusion coefficient of the free VOCs | $D_S$ [m$^2$ s$^{-1}$] | $7.58 \times 10^{-10}$ [a] |
| Diffusion coefficient of the free nsLTPs and the nsLTP·VOC complexes | $D_E$ and $D_C$ [m$^2$ s$^{-1}$] | $1.75 \times 10^{-10}$ [b] |
| Cuticle-water partition coefficient of VOC* | $K_{c/w}$ [-] | $10^{2.69}$ [c] |
| VOC concentrations in the cuticle* | $C_{Cut}$ [mol m$^{-3}$] | 12 [d] |
| The cell wall length | $L$ [nm] | 400 [d] |
| Concentration of LTP in the cell wall | $E_T$ [mM] | 1 for Fig. 6b-d |
| Dissociation constants | $K_d$ [μM] | Variable values [e] |

[a] Value for methyl benzoate obtained from Yaws[5] at 25 ºC.

[b] Value estimated based on Stokes–Einstein equation at 25 ºC. The radius of PhnsLTP3 was estimated by the correlation from Erickson[6] assuming a spherical protein structure.

[c] Value for methyl benzoate obtained from Ray et al.[7].

[d] Value for methyl benzoate obtained from Widhalm et al.[8].

* These parameters were used to calculate $S_L$ using equation 6 in Supplementary Material. A range of $S_0$, which was arbitrarily set to be larger than $S_L$.

[e] Values were obtained from Figure 5 and Supplementary Figure 17.
